# Targeting a novel chloroquine derivative to lysosomes induces massive and irreversible damage to lysosomes and suppresses autophagosomes and lysosomes assembly in cancer

**DOI:** 10.1101/2025.09.07.674530

**Authors:** Nitish Chauhan, Ananda Guha Majumdar, Sujit Kumar Bhutia, Papiya Dey, Mahesh Subramanian, Birija Sankar Patro

**Affiliations:** Bio-Organic Division, Bhabha Atomic Research Centre, Mumbai, Maharashtra, India – 400085; Homi Bhabha National Institute, Anushaktinagar, Mumbai, Maharashtra, India – 400094; Department of Life Science, National Institute of Technology, Rourkela, Odisha 769008, India

**Keywords:** Chloroquine, *trans*-4, 4′-dihydroxystilbene, Lysostilbenes, TFEB, lysosome targeting, pancreatic cancers

## Abstract

Pancreatic ductal adenocarcinoma (PDAC) exhibits profound therapy resistance driven by lysosome dependent nutrient recycling, metabolic adaptation, and stress tolerance. Current lysosome-targeting agents such as chloroquine (CQ) and hydroxychloroquine (HCQ) show limited efficacy due to transient activity and dose-limiting toxicities. To overcome these limitations, we developed Lysostilbenes, a new class of hybrid small molecules combining the CQ pharmacophore with lysosome-disrupting *trans*-4,4′-dihydroxystilbene. Lysostilbene-4 emerged as the lead candidate, demonstrating ∼30–40-fold greater cytotoxicity against PDAC cells than parental compounds, while sparing non-malignant cells. At nanomolar concentrations, Lysostilbene-4 induced rapid, irreversible lysosomal membrane permeabilization (LMP), initiating a lysosome mitochondria apoptotic cascade *via* cathepsin-B release, BID cleavage, BAX activation, and caspase-mediated apoptosis. In parallel, it abrogated lysosomal recovery by impairing repair, lysophagy, autophagosome maturation, and uncoupling TFEB-driven transcriptional programs from effective lysosome biogenesis. TFEB knockout further sensitized PDAC cells, underscoring TFEB as a key determinant of lysosomal resilience and a potential predictive biomarker. Importantly, Lysostilbene-4 was well tolerated in preclinical mouse models at supra-therapeutic doses without systemic toxicity. These findings position Lysostilbene-4 as a first-in-class lysosome-targeting therapeutic that enforces sustained lysosomal collapse while disabling adaptive recovery mechanisms, providing a mechanistically precise and safe strategy against PDAC.

## 1. Introduction

Accumulating evidence implicates lysosomes as central mediators of radio- and chemoresistance in cancers, primarily through drug sequestration, efflux *via* exocytosis, and regulation of key metabolic signaling pathways [1,2]. As the hub of endosomal trafficking and a critical regulator of tumor growth, progression, and therapy resistance, lysosomes have emerged as attractive therapeutic targets [3]. Current strategies largely focus on either disrupting lysosomal membrane integrity to induce lysosomal membrane permeabilization (LMP) or targeting lysosome-associated signaling regulators such as TFEB, AMPK, and cell-cycle effectors [4,5].

Lysosomal damage triggers a tripartite adaptive program, <u>R</u>epair, <u>R</u>emoval (lysophagy) and <u>R</u>egeneration (biogenesis) termed the “3R response of lysosomes” [6,7]: (1) ESCRT-dependent or -independent membrane repair, (2) galectin-mediated sensing and ubiquitin-driven removal through lysophagy, and (3) TFEB-driven lysosomal regeneration through biogenesis. While lysosome-targeting agents show therapeutic promise, their clinical translation is hindered by two major challenges: (a) activation of 3R responses that counteract lysosomal disruption and diminish cytotoxic efficacy, and (b) off-target toxicities in normal tissues due to insufficient tumor selectivity [8,9]. For instance, L-leucyl-L-leucine methyl ester (LLOMe) induces rapid lysosomal disruption in cancer cells; however, this damage is often rapidly repaired by the 3R machinery [7], limiting cell death induction [10]. Notably, a recent study demonstrated that LLOMe-induced lysosomal injury can promote programmed survival and regenerative responses, highlighting its potential application in regenerative medicine [11]. Currently, no lysosome targeting drug is available, which can cause lysosome specific damage and simultaneously overcome the lysosome repair, removal and regeneration to induce potent killing of cancer cells with minimum or no effects on normal cells.

Chloroquine (CQ) and hydroxychloroquine (HCQ) currently represent the only clinically approved autophagy and lysosomal inhibitors, with over 40 ongoing clinical trials assessing their therapeutic efficacy either as monotherapies or in combination with as surgery, radiotherapy, or chemotherapy [8,9]. Their perioperative application has been linked to improved pathological responses and survival outcomes [12,13]. A meta-analysis of seven clinical trials integrating CQ or HCQ with standard-of-care therapies across a spectrum of malignancies-including glioblastoma, brain metastases from non-small cell lung cancer and breast cancer, non-Hodgkin lymphoma, and pancreatic adenocarcinoma-demonstrated consistent improvements in overall response rate (ORR), progression-free survival (PFS), and overall survival (OS) [14]. Pancreatic ductal adenocarcinoma (PDAC), one of the most treatment-refractory solid tumors, exhibits constitutively elevated lysosomal activity, which supports tumor cell survival and progression. Pharmacological inhibition of lysosomal function has been shown to attenuate PDAC cell proliferation and tumor growth. In a randomized clinical trial (NCT01978184), preoperative administration of HCQ in combination with gemcitabine and nab-paclitaxel significantly enhanced pathological responses in patients with resectable pancreatic adenocarcinoma [13]. Despite these encouraging findings, the clinical implementation of CQ and HCQ faces major limitations: absence of predictive biomarkers for patient selection, and dose-limiting toxicities including cardiotoxicity, retinopathy, and lethality at high CQ doses (≥50 mg/kg orally) [8]. Taken together, while CQ and HCQ provide a valuable proof-of-concept for targeting the autophagy-lysosome axis in cancer, their clinical limitations underscore the urgent need for the development of next-generation CQ derivatives with enhanced potency, reduced toxicity, and improved patient stratification strategies—particularly for overcoming therapeutic resistance in PDAC.

In this study, we aimed to develop a novel class of chloroquine-based derivatives capable of inducing lysosome-specific damage to selectively sensitize cancer cells while minimizing toxicity to normal cells. Building on our previous work demonstrating that *trans*-4,4’-dihydroxystilbene (DHS) and resveratrol induces potent lysosomal membrane permeabilization (LMP) and suppress tumor burden [15,16], we hypothesized that conjugation of the lysosomotropic chloroquine pharmacophore with the lysosome-disrupting DHS moiety *via* an alkyl linker could enhance therapeutic efficacy. Accordingly, we designed, synthesized, and characterized a new class of hybrid molecules termed “Lysostilbenes”. Among these, Lysostilbene-4 exhibited a 30-40-fold increase in cytotoxic potency against pancreatic ductal adenocarcinoma cells compared to its parent compounds, with minimal impact on non-malignant cells. Notably, Lysostilbene-4 selectively disrupts lysosomes and concurrently inhibits the 3R responses for lysosomal recovery: membrane <u>r</u>epair, <u>r</u>emoval (lysophagy), and <u>r</u>egeneration/biogenesis of lysosomes. Furthermore, acute administration of Lysostilbene-4 was well tolerated *in vivo* in murine models. These findings introduce Lysostilbenes as a promising new class of lysosome-targeting agents with significant therapeutic potential against treatment-refractory pancreatic ductal adenocarcinoma cells.

## 2. Results

### 2.1. Design and synthesis of lysosome targeting chloroquine-stilbene hybrids

For targeting lysosome, a series of molecules have been prepared by combining the pharmacophores of Chloroquine i.e., 7-chloro-4-(4-diethylamino-1-methylbutylamino)-quinoline and 4,4’-dihydroxystilbene (DHS). 4-amino-7-chloroquinoline pharmacophore of Chloroquine, is known to be primarily protonated and trapped in the acidic lumen of lysosomes (Fig. 1A). DHS, a potent natural resveratrol analogue, is known to induce lysosome permeabilization, albeit at high concentration (Fig. 1B). We anticipated that conjugating DHS to Chloroquine pharmacophore may specifically target DHS to lysosome damage with no major impact on other cellular organelles (Fig. 1C). We named these probable <u>lyso</u>somes targeting 4,4’-dihydroxy<u>stilbenes</u> as Lysostilbenes (LS). We used 3 different approaches to make lysostilbenes: (1) Class-I Lysostilbenes: 4-amino group of chloroquine is modified and conjugated with DHS. Here, the conjugated molecules cannot be protonated, (2) Class-II Lysostilbenes: 4-amino group of chloroquine is conjugated with DHS. Here, the conjugated molecules retain protonation ability. (3) Class-III Lysostilbenes: 4-amino group of chloroquine is conjugated with DHS while 7-chlorine group is modified.

**Figure 1.**
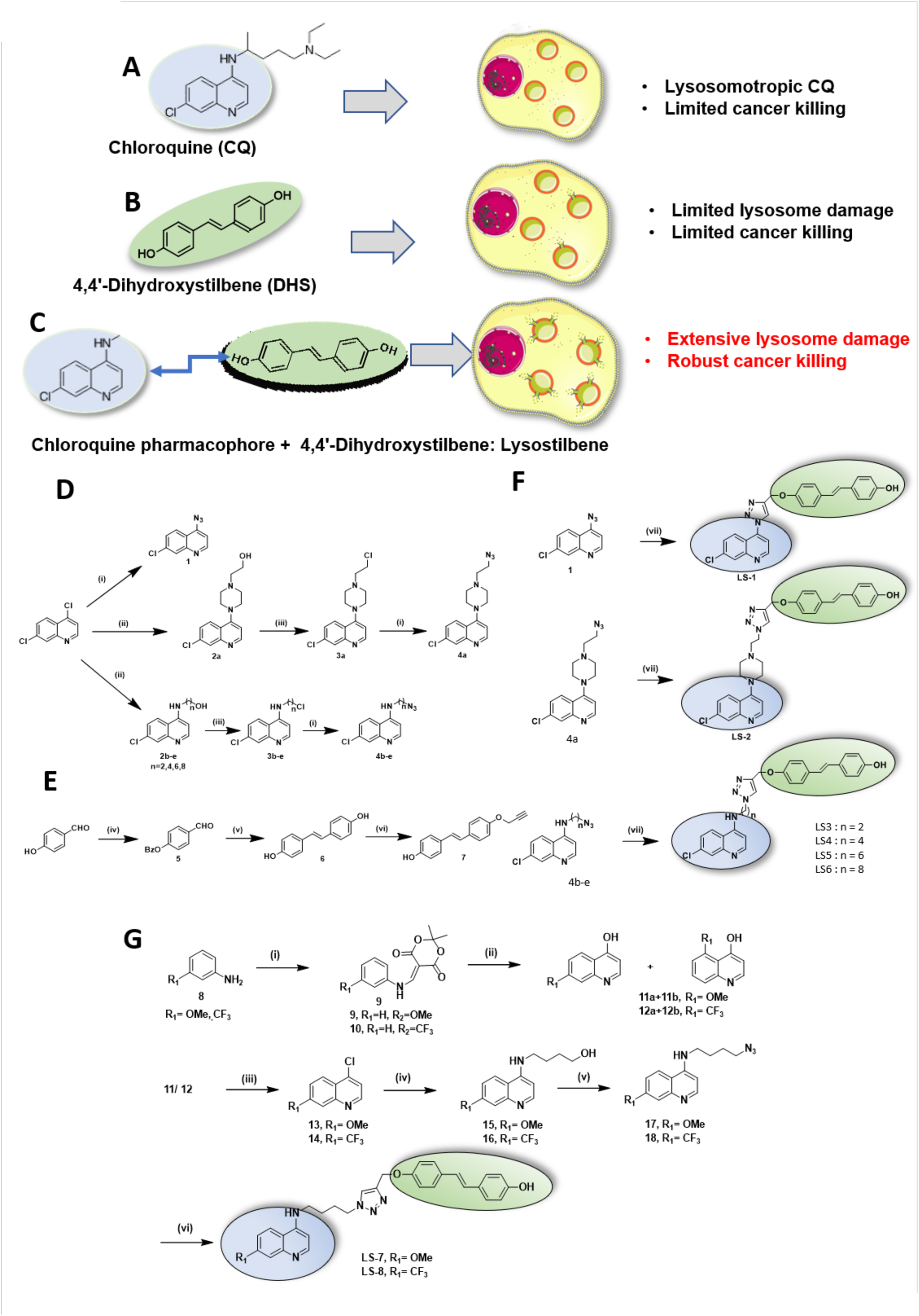
Design and synthesis strategy of Lysostilbenes, hybrid molecules for sensitizing pancreatic cancers. (A-C) Schematics showing structure and properties of chloroquine (CQ), *trans*-4,4′-Dihydroxystilbene (DHS) structure, and hybrid conjugates. (D) Scheme for Synthesis of azides and monopropargyl-DHS. Reagents and conditions: (i) NaN_3_/ DMF/ 100°C/ 6 h; (ii) 1- (2-Hydroxyethyl)piperazine (for 2a) or H_2_N(CH_2_)_n_OH (for 2b-e)/ 120°C/ 5 h; (iii) SOCl_2_/ cat. DMF/ CH_2_Cl_2_/ 24 h; (iv) BzCl/ Et_3_N/ 0°C-RT/ 2.5 h, (v) TiCl_4_/ Zn/ THF/ 0°C-reflux/ 8 h; (vi) propargyl bromide/ K_2_CO_3_/ acetone/ 45°C/ 18 h. (E) Scheme for synthesis of 1,2,3-triazole tethered lysostilbenes (LS1-6). Reagents and conditions: (vii) 7/ CuSO_4_.5H_2_O/ Na-ascorbate/ethanol-H_2_O/ 16-18 h. (F) Scheme for synthesis of LS-7 and LS-8. Reagents and conditions: (i) Meldrum’s acid, CH(OEt)_3_, reflux 1 h, then appropriate aniline, reflux 2 h; (ii) diphenyl ether/ 240°C/ 15-45 min; (iii) POCl_3_/ reflux, 3 h; (iv) a) H_2_N(CH_2_)_n_OH/ 120°C/ 5 h; b) SOCl_2_/ cat. DMF/ CH_2_Cl_2_; (v) NaN_3_/ DMF/ 100°C/ 6 h; (vi) 7/ CuSO_4_.5H_2_O/ Na-ascorbate/ ethanol-H_2_O/16-18 h

For synthesis of these lysostilbenes, two pharmacophores were joined with 1,2,3-triazole linkers using well-known Cu(I)-catalysed click chemistry. Detailed process of synthesis and extensive chemical characterization of these lysostilbenes molecules are given in materials and methods section (Fig. S1-S60). Briefly, 4,7-dichloroquinoline, as a starting material, was either directly converted to the corresponding azide **1** or treated with neat aminoalcohols (120 °C; 5 h) to get compounds **2a-e** (Fig. 1D). Then chlorination of the hydroxyl groups of **2a-e** followed by azidation gave the azides **4a**-**e** (Fig. 1D). Another partner for click synthesis was prepared by monopropargylation of *trans*-4,4’-dihydroxystilbene (Fig. 1E). For synthesis of class-I Lysostilbenes, compound **1** and **4a** were coupled with monopropargylated DHS (**7**) to get the Lysostilbenes-1, 2 (**LS1, LS2**) *via* Cu(I)-catalysed 1,3-dipolar cycloaddition (Fig. 1F), For class-II stilbenes, compound **4b-e** were coupled with monopropargylated DHS (**7**) to get the Lysostilbenes-3-6 (**LS3-6**) *via* Cu(I)-catalysed 1,3-dipolar cycloaddition (Fig. 1F). **LS3-6** have varied carbon numbers in the spacer between the pharmacophores of Chloroquine and DHS. For class-III Lysostilbenes, the precursors (compound **8-18;** Fig. 1G) were synthesized as per protocol described by Madrid *et al.*[17]. Finally, compound **17** and **18** were coupled with monopropargylated DHS (**7**) using click chemistry to get the Lysostilbenes-7, 8 (**LS7**, **LS8)** with 1,2,3-triazole linker (Fig. 1G).

### 2.2 Lysostilbene-4 induces potent and selective cytotoxicity in pancreatic cancer cells

The antiproliferative activity of newly synthesized Lysostilbenes (LSs) was evaluated against two pancreatic ductal adenocarcinoma (PDAC) cell lines, MIA PaCa-2 and PANC-1, using a clonogenic assay (Fig. 2A, B). Parent compounds, Chloroquine and DHS, were also tested for comparison with Lysostilbenes. Our findings show that Chloroquine and DHS exhibit IC_50_ values in the range of 1-10 μM for both PDAC cell lines (Fig. 2A-D). In contrast, class-I lysostilbenes (Lysostilbene-1 and Lysostilbene-2) did not display enhanced efficacy up to 1 μM (Fig. 2A-D), indicating that blocking the protonation function of the 4-amino group in chloroquine pharmacophores does not confer additional benefit in cancer cell killing. Previously, the significance of 4-amino group in the chloroquine scaffold, demonstrating its crucial role in chloroquine potency was reported [18]. Class-II Lysostilbenes (Lysostilbenes-3-6) displayed pronounced potency, with Lysostilbene-4 showing exceptional efficacy at nano molar concentrationapproximately 30-40-fold greater than the parent compounds (Chloroquine and DHS) (Fig. 2A-F). IC_50_ values of Lysostilbene-4 are ∼105 nM for both MIA PaCa-2 and PANC-1 cells. The enhanced anticancer activity within this class appears to depend on (a) the protonation capacity of the 4-amino group in the CQ pharmacophore and (b) the alkyl chain length linking the CQ and DHS pharmacophores. The remarkable potency of Lysostilbene-4 may be attributed to its four-carbon alkyl spacer, which likely provides optimal flexibility between the two conjugated pharmacophores, thereby facilitating its biological activity. To evaluate the contribution of additional chemical substituents in Lysostilbene-4, we synthesized Class-III lysostilbenes by replacing its 7-chloro group with either a methoxy (Lysostilbene-7) or trifluoromethyl (Lysostilbene-8) moiety. Both Lysostilbene-7 and Lysostilbene-8 exhibited markedly reduced antiproliferative activity compared with Lysostilbene-4 (Fig. 2A-D), underscoring the critical role of the 7-chloro substituent in mediating the efficacy of Lysostilbene-4. The anticancer activity of lysostilbenes was further evaluated using the MTT assay. Consistent with the clonogenic assay, Lysostilbene-4 emerged as the most potent compound among those tested (Fig. 2G, H). To examine its selectivity, we compared the cytotoxic effects of Lysostilbene-4 on pancreatic cancer cells (PDAC) versus non-malignant cell lines (MCF10A and Vero). Notably, Lysostilbene-4 induced significantly greater cytotoxicity in PDAC cells than non-malignant cells (Fig. 2I). Together, these findings highlight the strong therapeutic potential of Lysostilbene-4 against pancreatic cancer and provide proof-of-concept for the novel design of pharmacophore-conjugated molecules with enhanced potency. To further delineate its mechanism of action, subsequent studies were performed with Lysostilbene-4 in MIA PaCa-2 pancreatic cancer cells.

**Figure 2.**
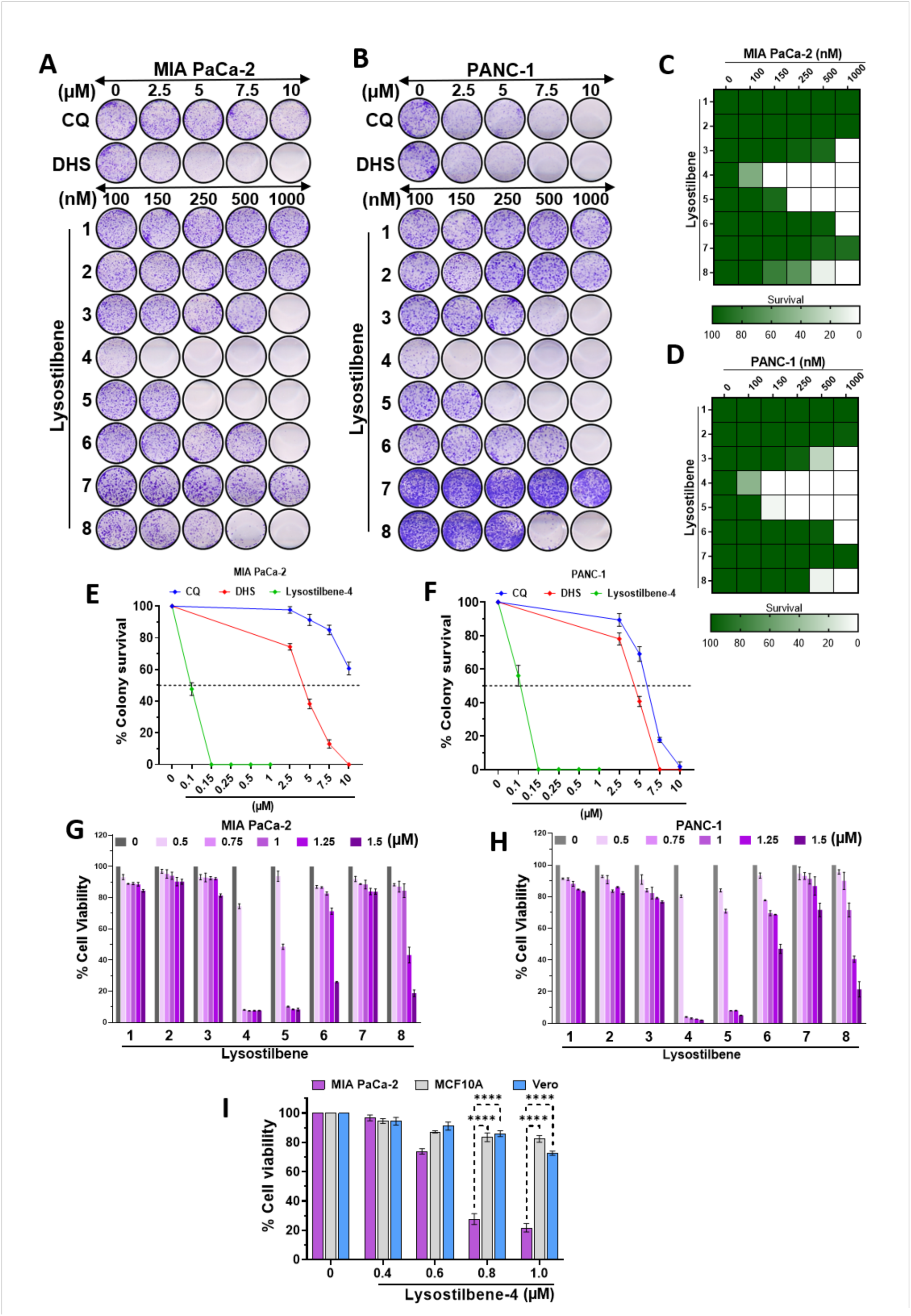
Lysostilbene-4 potently suppresses colony formation and viability of pancreatic cancer cells. (A, B) Clonogenic survival assay of MIA PaCa-2 and PANC-1 cells treated with increasing concentrations of Lysostilbenes (100-1000 nM) along with parent molecule as CQ (2.5-10 µM) or DHS (2.5-10 µM) for 8-10 days. Representative images of crystal violet–stained colonies are shown. (C, D) Heat-maps of clonogenic survival corresponding to above assays, where dark green indicates 100% colony survival and white represents 0%. (E, F) Dose–response curves showing percentage of colony survival in MIA PaCa-2 and PANC-1 in above assays. Error bars represent mean ± SEM, N = 3 biological replicates. (G, H) Percentage cell viability of MIA PaCa-2 and PANC-1 cells treated with Lysostilbene-4 (0.5-1.5 µM) for 72 h, evaluated by MTT assay. Error bars represent mean ± SEM, N = 3 biological replicates. (I) Percentage cell viability of pancreatic cancer cells (MIA PaCa-2) and non-cancerous cells (MCF10A and Vero), treated with Lysostilbene-4 for 72 h and followed by MTT assay. Error bars represent mean ± SEM, N= 3 biological replicates. (\*\*\*\**p* < 0.0001).

### 2.3 Lysostilbene-4 selectively disrupts lysosomes and induces irreversible lysosomal damage

Numerous agents have been reported to perturb lysosomal function; however, most act non-specifically, only a few including LLOMe [10,19], glycyl-L-phenylalanine 2-naphthylamide (GPN) [19], Triamterene [20] have been reported to selectively target lysosomes with minimal effects on other organelles. However, these agents typically elicit lysosomal disruption only at millimolar concentrations (0.5-2.0 mM), limiting their translation potential as lysosome-specific anticancer therapeutics. Given the potent anticancer activity of Lysostilbene-4 at nanomolar doses, we next evaluated its ability to selectively target lysosomes. To assess lysosomal integrity, we employed an immunofluorescence-based galectin-3 (Gal-3) assay, a sensitive and specific marker of lysosomal damage [21]. Normally, cytosolic Gal-3 translocates into damaged lysosomes and binds β-galactoside residues on luminal glycoproteins of the lysosomal membrane (Fig. 3A). Dose-standardization experiments revealed that Lysostilbene-4 rapidly induced Gal-3 puncta formation at 500 and 750 nM, with prominent foci at 750 nM (Fig. 3B, Fig. S61). Accordingly, subsequent studies were performed at 750 nM to capture its lysosome-specific effects. LLOMe (1 mM) served as a positive control and comparator for potential mechanistic differences. Strikingly, Lysostilbene-4 induced Gal-3 puncta within 2 h at nanomolar concentrations, whereas LLOMe required >1000-fold higher doses (Fig. 3B-D). Although Gal-3 puncta frequency and the percentage of Gal-3-positive cells were higher with LLOMe at 2 h compared to 4 h, Lysostilbene-4 elicited equal or greater Gal-3 accumulation at 4 h (Fig. S62). Live-cell imaging using acridine orange (AO), which fluoresces red in intact acidic lysosomes and green in cytoplasm, further confirmed robust lysosomal membrane permeabilization (LMP). Significant loss of functional lysosomes was observed within 2 h of treatment with LLOMe (1 mM), CQ (10 μM), or DHS (10 μM), while Lysostilbene-4 induced a similar extent of LMP at only 750 nM (Fig. 3E, F). Consistent with prior findings that LLOMe-induced LMP is transient and reversible within 24 h [22]. We next compared the temporal dynamics of lysosomal damage using Magic Red cathepsin B substrate (MRCB) co-stained with LysoTracker Green (LTG), demonstrating rapid but recoverable lysosomal impairment in LLOMe-treated cells (Fig. 3G-K). In contrast, Lysostilbene-4 induced only modest early damage (30-60 min), but this progressively escalated to extensive and irreversible lysosomal disruption by 24 h, with complete loss of lysosomal integrity and no evidence of lysosomal repair or regeneration (Fig. 3H-K). To confirm lysosomal selectivity and exclude major off-target effects, flow cytometry analyses were performed using AO, MitoTracker Red (MTR), and ER-Tracker Green (ETG) to monitor lysosomal, mitochondrial, and ER functions, respectively (Fig. S63A-F). Lysostilbene-4 selectively impaired lysosomes within 2 h, with no discernible effects on mitochondria or ER up to 6 h. Similarly, its parent molecules, CQ and DHS, exhibited negligible effects on mitochondria or ER under these conditions (Fig. S63A-F). Collectively, these results demonstrate that Lysostilbene-4 acts as a highly selective lysosomal disruptor, inducing persistent and irreversible lysosomal membrane permeabilization at nanomolar concentrations, thereby distinguishing it from classical lysosomotropic agents.

**Figure 3.**
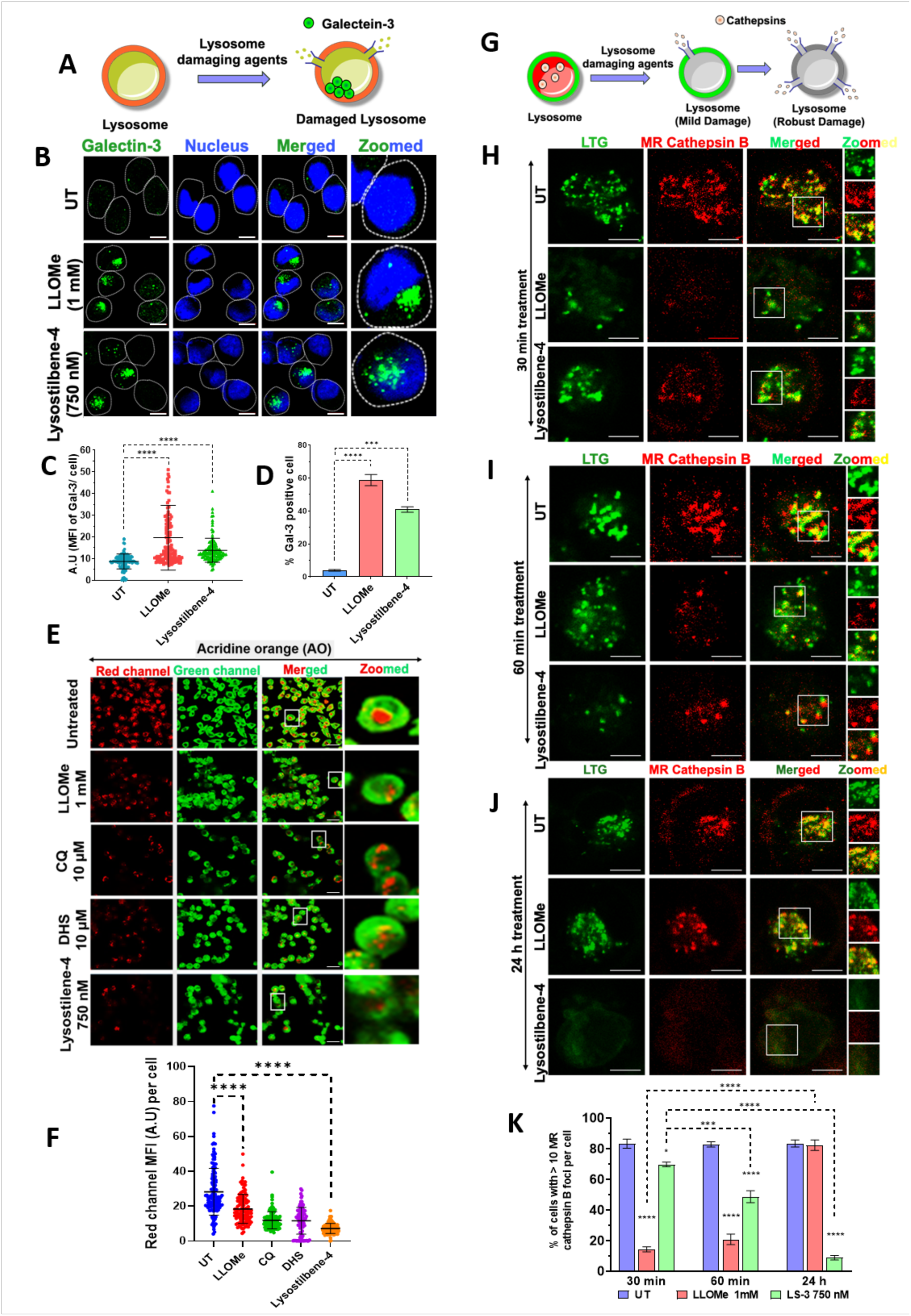
Lysostilbene-4 induces persistent lysosomal permeabilization (LMP). (A) Schematic illustration of Galectin-3 recruitment assay for LMP detection. (B) Representative confocal images showing Galectin-3 puncta formation (green) in untreated (UT), LLOMe or Lysostilbene-4 treatment for 2 h. Scale bars = 10 µm. (C, D) Quantification of Galectin-3 mean fluorescence intensity (MFI) and percentage of Galectin-3–positive cells in the above assay. Error bars represent mean ± SD; N= 3 biological replicates with >130 cells per sample. (E, F) Representative live cell microscopy images for assessment of functional lysosome (red fluorescence) status using Acridine orange (AO) dye with the indicated treatments for 2 h. Scale bars = 20 µm. Error bars represent mean ± SD; N= 3 biological replicates with >120 cells per sample. (G) Schematic of lysosomal damage progression leading to cathepsin release. (H-J) Confocal images showing Cathepsin B activity (Magic Red, MR, red) and lysosomal localization (Lysotracker Green, LTG, green) following LLOMe or Lysostilbene-4 treatment for 30 min, 60 min, 24 h. Scale bars = 10 µm. Error bars represent mean ± SEM, N= 3 biological replicates. (\*\*\**p* < 0.001; \*\*\*\**p* < 0.0001).

### 2.4 Lysostilbene-4 induces lysosomal membrane permeabilization-driven apoptosis signalling in pancreatic cancer cells

Several studies have demonstrated that LMP-associated cathepsin release promotes proteolytic BID cleavage and subsequent caspase activation, thereby initiating mitochondria-mediated intrinsic apoptosis in cancer cells [16,23,24]. Given the pronounced lysosomal membrane damage induced by Lysostilbene-4, it is plausible that the initial leakage of lysosomal proteases into the cytoplasm contributes to apoptosis at later stages (24 h). To assess LMP-mediated cathepsin B release, we employed a modified version of a previously established protocol [22,25] (Fig. 4A). In this assay, limited plasma membrane permeabilization with digitonin allows cytoplasmic cathepsins to be released into the extracellular medium while minimally affecting lysosomes. Cathepsin activity is subsequently detected using the Magic Red cathepsin B (MRCB) substrate (Fig. 4A).

**Figure 4.**
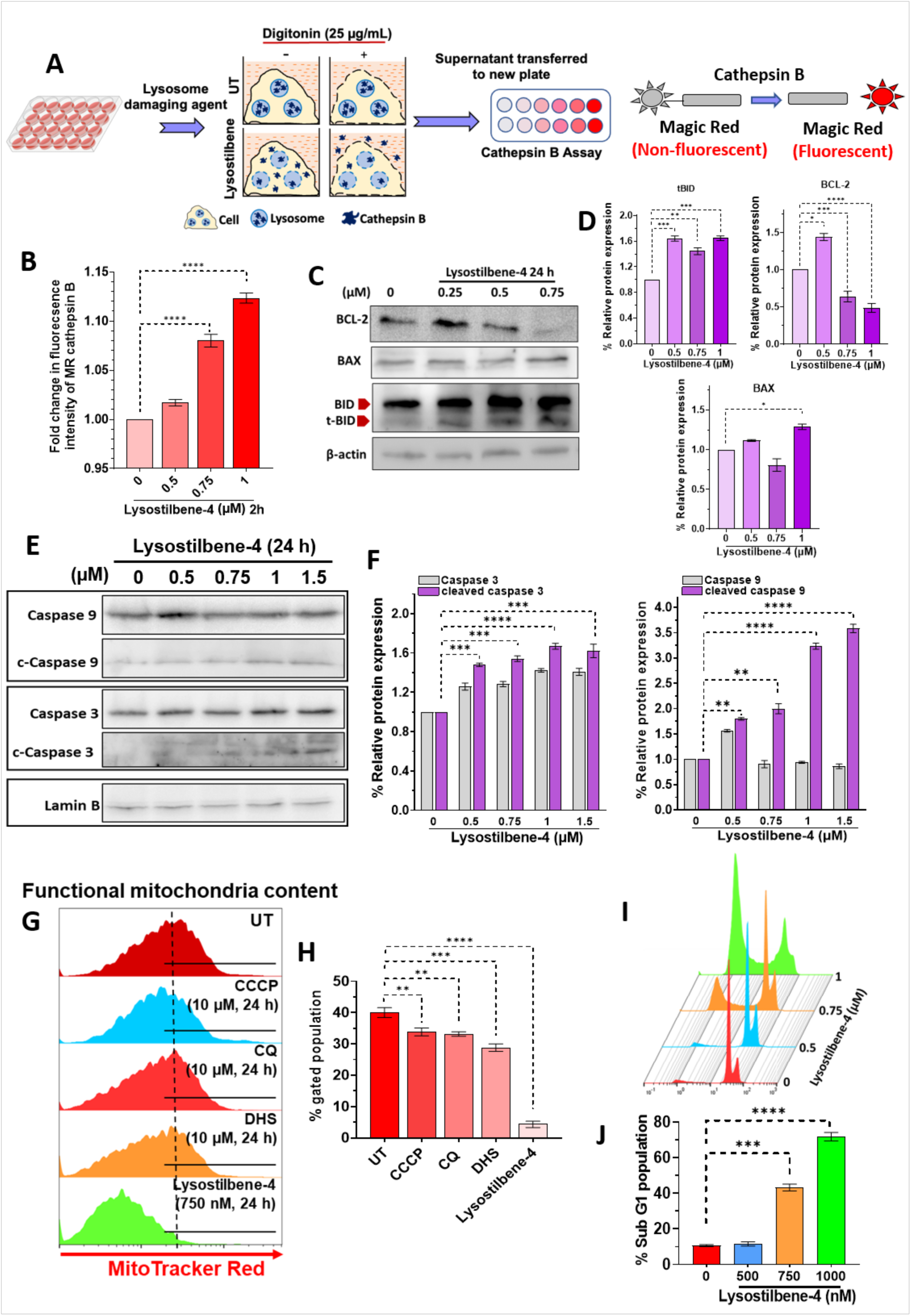
Lysostilbene-4–induced LMP triggers mitochondrial dysfunction and apoptotic signalling. (A, B) (A) Schematic representation of the Cathepsin B release assay. Cells were treated with Lysostilbene-4 for 2 h, followed by selective plasma membrane permeabilization with digitonin. The resulting supernatants were transferred to a fresh plate and Cathepsin B activity was measured using the Magic Red substrate, which generates fluorescence upon enzymatic cleavage (non-fluorescent → fluorescent). Fold change in fluorescence intensity of Magic Red substrate in the supernatant was quantified. Error bars represent mean ± SEM, N= 4 biological replicates. (C, D) Immunoblot and densitometric quantification of BCL-2, BAX, BID, and truncated BID from cell extracts, prepared after cells were treated with Lysostilbene-4 for 24 h. β-actin served as a loading control. Error bars represent mean ± SEM, N= 3 biological replicates. (E, F) Immunoblot and quantification of Caspase 3 and Caspase 9 activation from cell extracts, prepared after cells were treated with Lysostilbene-4 for 24 h. Lamin B served as loading control. Error bars represent mean ± SEM, N= 3 biological replicates. (G, H) Flow cytometry analysis of functional mitochondrial content using MitoTracker Red in the indicated treatment conditions for 24 h. Error bars represent mean ± SEM, n=4 biological replicates. (I, J) Flow cytometry-based cell cycle profile for Sub-G1 of MIA PaCa-2 cells treated with Lysostilbene-4 (500, 750 and 1000 nM) for 72 h. Error bars represent mean ± SEM, N= 4 biological replicates. (\*\**p* < 0.01; \*\*\**p* < 0.001; \*\*\*\**p* < 0.0001).

Our results showed that Lysostilbene-4 treatment (2 h) significantly increased cytosolic cathepsin B release in a dose-dependent manner (Fig. 4B) consistent with early lysosomal leakage. Moreover, prolonged exposure (24 h) led to enhanced BID cleavage into its truncated form (t-BID), accompanied by decreased BCL-2 and increased BAX expression (Fig. 4C, D), suggesting a delayed mitochondrial engagement in apoptosis. These events further supported by elevated levels of cleaved caspase-3 and caspase-9 (Fig. 4E, F), as well as a marked loss of mitochondrial membrane potential (MMP) at later time point (24 h, Fig. 4G, H). Strikingly, the extent of MMP collapse in Lysostilbene-4-treated cancer cells (24 h) was far greater than that observed with the parent compounds (CQ and DHS) or the positive control CCCP (Fig. 4G, H), underscoring its potent activation of the intrinsic apoptotic pathway. Finally, flow cytometry analysis revealed a significant increase in the apoptotic sub-G1 population following Lysostilbene-4 treatment (24 h) (Fig. 4I, J). Collectively, these findings establish that Lysostilbene-4 initiates cell death through a lysosome-to-mitochondria apoptotic cascade, wherein early cathepsin B release drives BID cleavage and subsequent mitochondrial dysfunction, culminating in caspase-dependent apoptosis.

### 2.5 Lysostilbene-4 impairs lysophagy and autophagosome assembly

The mode of action of Lysostilbene-4 is particularly noteworthy, as it induces near-complete depletion of cellular lysosomes with no evidence of recovery through repair and biogenesis for a long period (24 h), in contrast to the reversible lysosomal disruption typically observed with LLOMe-induced damage. To investigate this unique mode of action further deeper, we examined Lysostilbene-4’s impact on both LMP and subsequent lysophagy. For real-time monitoring the kinetics of LMP, we used MIA PaCa-2 cells expressing the tandem fluorescent reporter tf-Gal3 (RFP-GFP-Gal3) [26]. Upon LMP, tf-Gal3 translocases to the damaged lysosomal membrane, appearing as yellow puncta (RFP^+ve^, GFP^+ve^) (Figure 5A). As lysophagy proceeds, the acidic lysosomal environment quenches the GFP fluorescence, leaving red-only puncta (RFP^+ve^, GFP^− ve^) [26]. Both LLOMe and Lysostilbene-4 induced rapid accumulation of yellow Gal3 puncta within 2 h (Fig. 5B, C). However, while LLOMe-treated cells showed a time-dependent reduction in yellow puncta and a corresponding increase in red-only puncta over 6-24 h, indicating efficient lysophagic clearance, Lysostilbene-4 failed to sustain this transition. Instead, yellow puncta diminished without significant accumulation of red-only puncta, suggesting impaired lysophagy (Fig. 5B-D).

**Figure 5.**
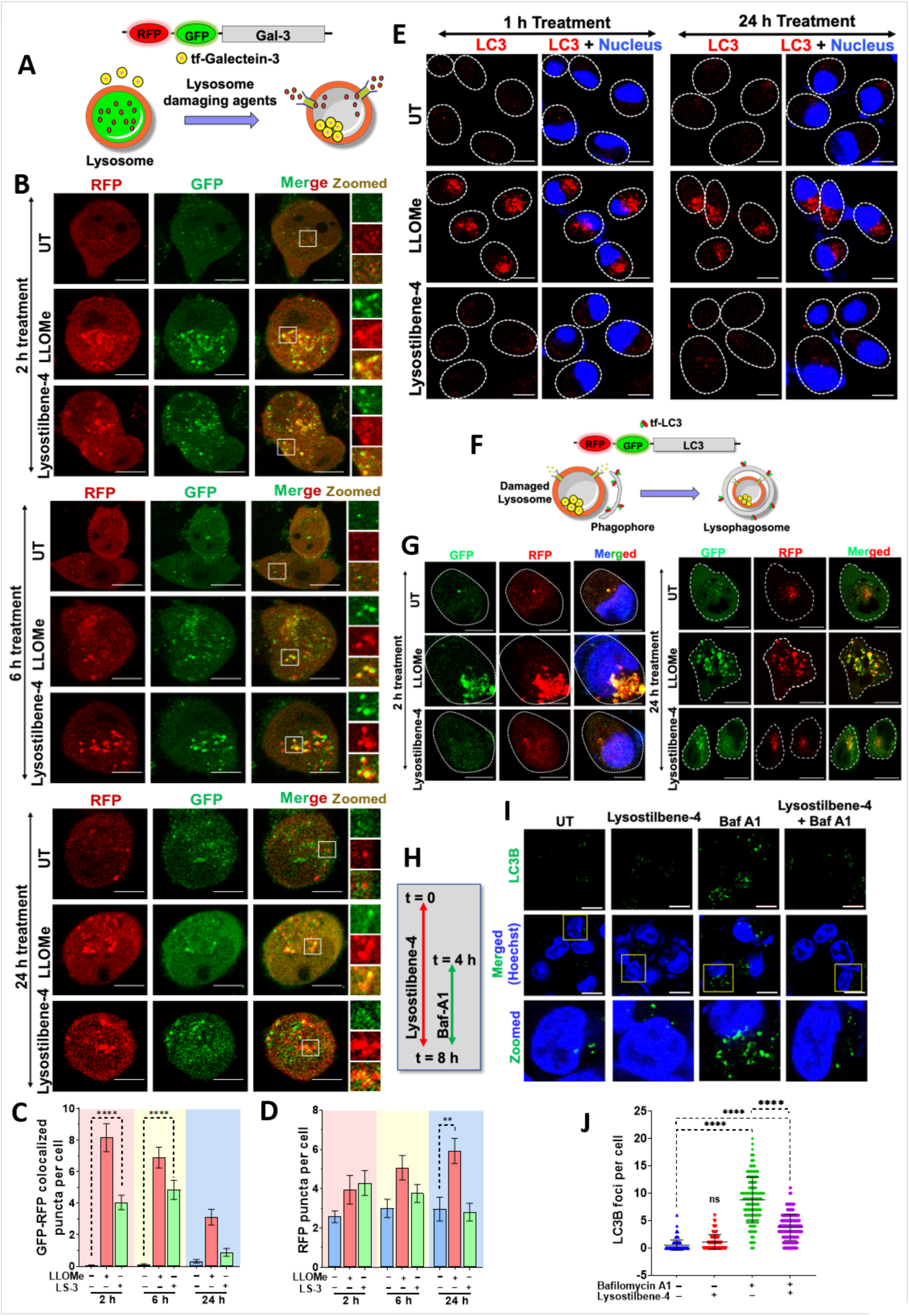
Lysostilbene-4 disrupts lysosomal integrity and impairs lysophagy for the removal of damaged lysosomes. (A) Schematic of the tandem fluorescence Galectin-3 (tf-Gal3) reporter system used to detect damaged lysosomes and its clearance dynamics by lysophagy. (B-D) Representative live cell confocal images of tf-Gal3-expressing Mia PaCa-2 cells treated with LLOMe and Lysostilbene-4 for 2 h, 6 h and 24 h, respectively. Insets show zoomed regions highlighting Gal3 puncta formation. Scale bars = 10 µm. Quantification of GFP-RFP colocalized puncta (GFP^+ve^ and RFP^+ve^) and RFP puncta (GFP^−ve^ and RFP^+ve^) per cell was quantified. Error bars represent mean ± SD, n= >100 cells per sample. (E) Immunofluorescence images of cells treated with indicated treatments for 1 h and 24 h, for the detection of endogenous LC3B foci formation. Scale bars = 10 µm. (F) Schematic illustration of lysophagy: sequestration of damaged lysosomes into phagophores to form lysophagosomes. (G) Confocal images of tf-LC3-expressing Mia PaCa-2 cells were treated with LLOMe and Lysostilbene-4 for 2 h and 24 h to assess autophagic flux. Scale bars = 10 µm. (H-J) Experimental scheme and confocal images of LC3B puncta following Lysostilbene-4 and Bafilomycin A1 treatments. Scale bars = 10 µm. Error bars represent mean ± SD, n= >180 cells per sample. (ns: not significant, \*\**p* < 0.01, \*\*\*\**p* < 0.0001).

We next assessed whether Lysostilbene-4 affects autophagosome biogenesis. Immunofluorescence analysis of LC3 revealed a robust accumulation of LC3 puncta in LLOMe-treated cells at both early (1 h) and late (24 h) time points, consistent with ongoing autophagy. In sharp contrast, Lysostilbene-4 almost completely abolished LC3 puncta formation despite unchanged total LC3 protein levels (Fig. 5E; Fig. S64A-D), indicating a block in autophagosome assembly rather than reduced LC3 expression. Further real time monitoring of kinetics of autophagsome formation was assessed using MIA PaCa-2 cells stably expressing the tandem fluorescent reporter tf-LC3 (RFP-GFP-LC3) (Figure 5F). In line with our previous results (Figure 5E), massive yellow tf-LC3 (RFP^+ve^ and GFP^+ve^) punctae formation was observed in LLOMe-treated cells (2 h and 24 h) whereas tf-LC3 (RFP^+ve^ and GFP^+ve^) punctae formation was severely impaired in Lysostilbene-4-treated cells (Figure 5F, G, Fig. S64E, F).

These findings prompted us to consider two mechanistic possibilities: (i) Lysostilbene-4 interferes with lysophagophore formation around damaged lysosomes, and/or (ii) it disrupts the general process of autophagosome assembly. To distinguish between these possibilities, MIA PaCa-2 cells were treated with Lysostilbene-4 in the presence or absence of Bafilomycin A1 (Baf-A1), a lysosomotropic inhibitor of autophagosome-lysosome fusion (Fig. 5H). As expected, Baf-A1 alone induced a robust accumulation of LC3 puncta, indicative of autophagosome build-up (Fig. 5I, J). Strikingly, Lysostilbene-4 co-treatment significantly attenuated Baf-A1-driven LC3 puncta accumulation (Fig. 5I, J), suggesting that Lysostilbene-4 impairs autophagosome formation. This effect of Lysostilbene-4 is likely attributable to extensive leakage of acidic lysosomal contents, including proteases (Fig. 4A, B), which may interfere with autophagosome assembly. Furthermore, given the pronounced LMP induced by Lysostilbene-4, we hypothesized that Lysostilbene-4 may also disrupt the turnover of pre-formed autophagosomes. To test this, cells were pre-exposed to Baf-A1 for 4 h to allow LC3 puncta (autophagosomes) accumulation, followed by a 24 h recovery period in the absence or presence of Lysostilbene-4 (Fig. S65A-C) autophagosome clearance was efficiently restored under control conditions but this clearance was almost completely abolished in Lysostilbene-4–treated cells (Fig. S65A-C). Collectively, these results demonstrate that Lysostilbene-4 disrupts lysosomal homeostasis at two levels: (i) by inducing irreversible LMP with impaired lysophagic clearance, and (ii) by inhibiting autophagosome biogenesis and turnover. This dual blockade leads to the accumulation of damaged lysosomes and defective autophagy, mechanistically positioning Lysostilbene-4 apart from classical lysosomotropic agents such as LLOMe, CQ, HCQ etc.

### 2.6 Lysostilbene-4 elicits TFEB-mediated transcriptional response for lysosome biogenesis in pancreatic cancers

Lysosomal stress is a well-known activator of transcription factor EB (TFEB), the master regulator of autophagy-lysosome gene (ALG) expression [6,27,28]. Under basal conditions, phosphorylated TFEB (S211) is sequestered in the cytoplasm, whereas lysosomal stress induces dephosphorylation and nuclear translocation, initiating transcriptional programs for lysosomal biogenesis [29,30]. To investigate whether Lysostilbene-4 triggers TFEB activation, we employed MIA PaCa-2 cells stably expressing eGFP-TFEB. Lysostilbene-4 treatment promoted robust, time- and dose-dependent nuclear accumulation of eGFP-TFEB (Fig. 6A-D; SI Fig. 66A, B), accompanied by a marked decrease in phosphorylated TFEB (S211) without altering total TFEB levels (Fig. 6E-G). Above result suggested activation of TFEB response for lysosome biogenesis in Lysostilbene-4 treatment.

**Fig 6.**
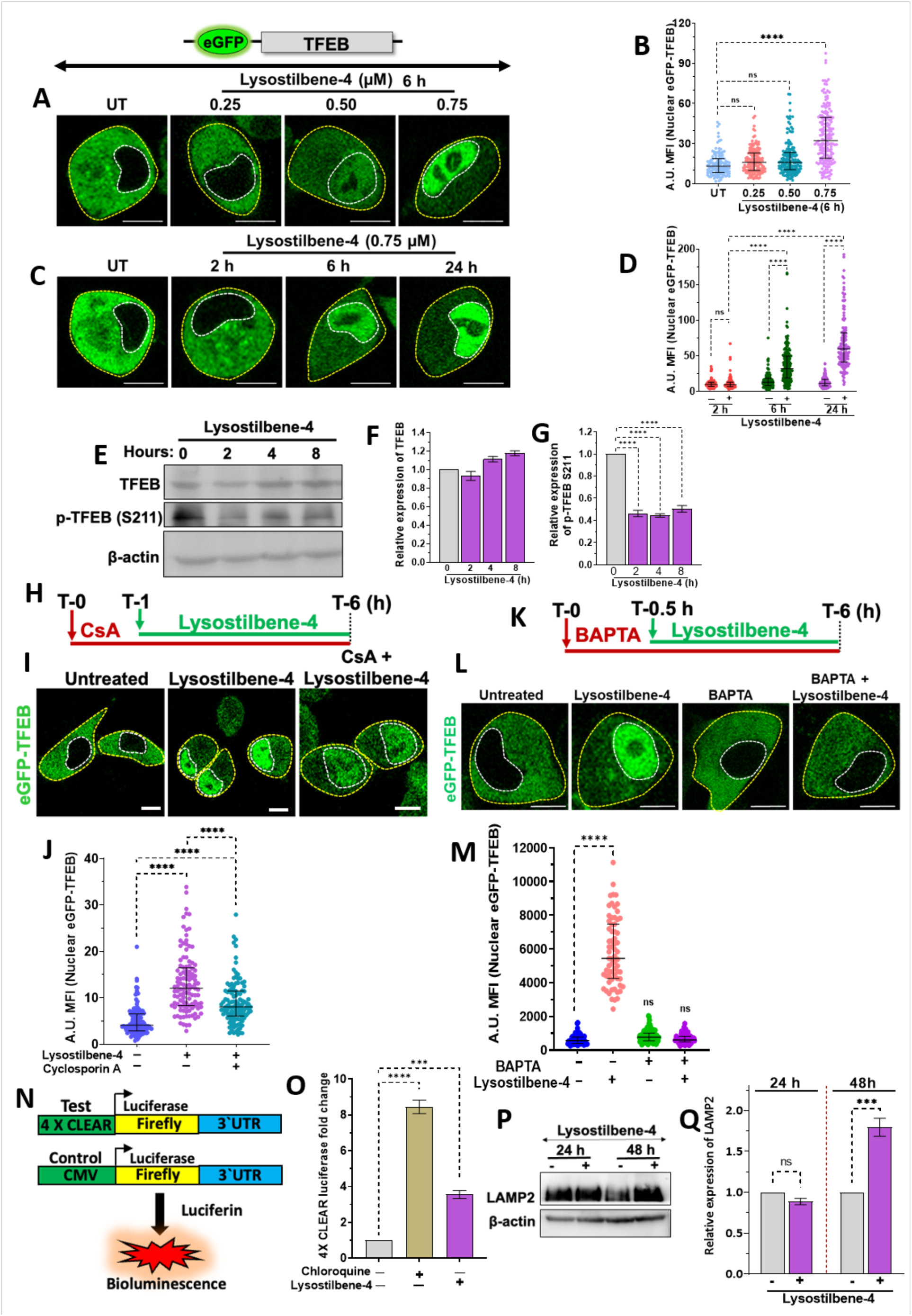
Lysostilbene-4 promotes nuclear translocation and activation of TFEB Ca²⁺-calcineurin signalling. (A-D) Representative live-cell microscopy images of MIA PaCa-2 cells, expressing eGFP-tagged TFEB, showing nuclear translocation of eGFP-TFEB upon treatment with increasing concentrations of Lysostilbene-4 (0.25-0.75 μM) for different time points. Scale bars: 10 μm. Mean fluorescence intensity (MFI) of nuclear eGFP-TFEB was quantified). Error bars represent mean ± SD, n= >150 cells per sample. (E-G) Immunoblots and quantification of TFEB and phosphorylated TFEB (Ser211) in cell extracts, prepared after treating MIA PaCa-2 cells with Lysosstillbene-4. β-actin served as a loading control. Error bars represent mean ± SEM, N= 3 biological replicates. (H) Experimental scheme for calcineurin inhibition assay using Cyclosporin A (CsA pre-treatment for 1 h, prior to Lysostilbene-4 exposure for 6 h. (I, J) Representative live cell confocal images and quantification as MFI of nuclear eGFP-TFEB for assessing TFEB’s nuclear accumulation. Error bars represent mean ± SD, n= >118 cells per sample. Scale bars: 10 μm. (K) Experimental scheme for calcium chelation using BAPTA-AM (0.5 h, pre-treatment) prior to Lysostilbene-4 exposure (6 h). (L, M) Representative live cell images and quantification as MFI of nuclear eGFP-TFEB for assessing TFEB’s nuclear accumulation under the treatment scheme mentioned above. Error bars represent mean ± SD, n= >120 cells per sample. Scale bars: 10 μm. (N) Schematic of CLEAR-luciferase reporter assay for measuring TFEB transcriptional activity. (O) Post transfection with reporter plasmid, MIA PaCa-2 cells were treated with Lysostilbene-4 (750 nM) for 24 h, the luciferase reporter activity was recorded, and the graphs show the relative luciferase activity from 4×CLEAR reporter in cells treated with chloroquine (positive control) or Lysostilbene-4. Error bars represent mean ± SEM, N= 4 biological replicates. (P,Q) Immunoblot and quantification of LAMP2, a TFEB target gene, in MIA PaCa-2 cells treated with Lysostilbene-4 for 24 h and 48 h. Error bars represent mean ± SEM, N= 3 biological replicates. (ns: not significant; \*\*\**p* < 0.001; \*\*\*\**p* < 0.0001).

Since calcium released during lysosomal damage activates calcineurin the primary phosphatase dephosphorylating TFEB at S211 [31,32] we investigated the involvement of this pathway. Pre-treatment with cyclosporin A (CsA, calcineurin inhibitor) or BAPTA (a Ca²⁺ chelator) significantly abrogated Lysostilbene-4-induced TFEB nuclear translocation (Fig. 6H-M), implicating Ca²⁺-calcineurin signaling in this process. To further confirm a role nuclear TFEB in transcriptional activation of autophagy and lysosome biogenesis genes (ALG), we employed a 4× CLEAR luciferase reporter, which harbors TFEB-binding motifs present in ALG promoters (Fig. 6N) [33]. Lysostilbene-4 treatment, as well as CQ treatment, significantly enhanced CLEAR-driven luciferase activity, validating TFEB-dependent transcriptional induction of autophagy-lysosome biogenesis genes (Fig. 6O). In corroboration with this, a substantial increase in the expression of LAMP2, a lysosomal membrane protein was observed in Lysostilbene-4 treated cells (Fig. 6P, Q), confirming a complete response of TFEB for lysosome biogenesis.

Together, these results demonstrate that although TFEB activation robustly activates TFEB through a Ca²⁺-calcineurin-dependent mechanism, driving the transcription of ALG expression, the subsequent biogenesis and assembly of functional lysosomes (Fig. 3G-K) and autophagosomes (Fig. 5E-J), which are are also profoundly impaired upon Lysostilbene-4 treatment. This represents a unique phenotype of Lysostilbene-4, not previously described for other lysosome-disrupting agents, characterized by its ability to induce sustained LMP while concurrently suppressing the coordinated “3R” response-lysosomal repair, removal *via* lysophagy, and regeneration through biogenesis.

### 2.7 Knocking out TFEB sensitizes pancreatic cancers to Lysostilbene-4

Low expression of DNA repair genes such as BRCA1 (breast/ovarian cancer), ERCC1 (lung cancer), and XRCC2 (glioma cancer) has long been recognized as a predictive biomarker for enhanced responsiveness to DNA damage-based chemotherapy. To evaluate the prognostic significance of TFEB, we analyzed TCGA (The Cancer Genome Atlas) datasets using online platform Kaplan Meier (KM) plotter. Reduced TFEB mRNA expression correlated with poor overall survival (OS) and disease-free survival (DFS) across multiple cancers (Fig. 7A), with a particularly strong association in pancreatic cancer (Fig. 7B). These clinical data suggest that TFEB expression has prognostic relevance and may influence therapeutic responses in pancreatic malignancies. Given the potent lysosome-targeting activity of Lysostilbene-4 in TFEB-high pancreatic cancers, we surmised that TFEB deficiency, by impairing functional lysosome biogenesis/assembly, may further sensitize low-TFEB tumors to Lysostilbene-4.

**Figure 7.**
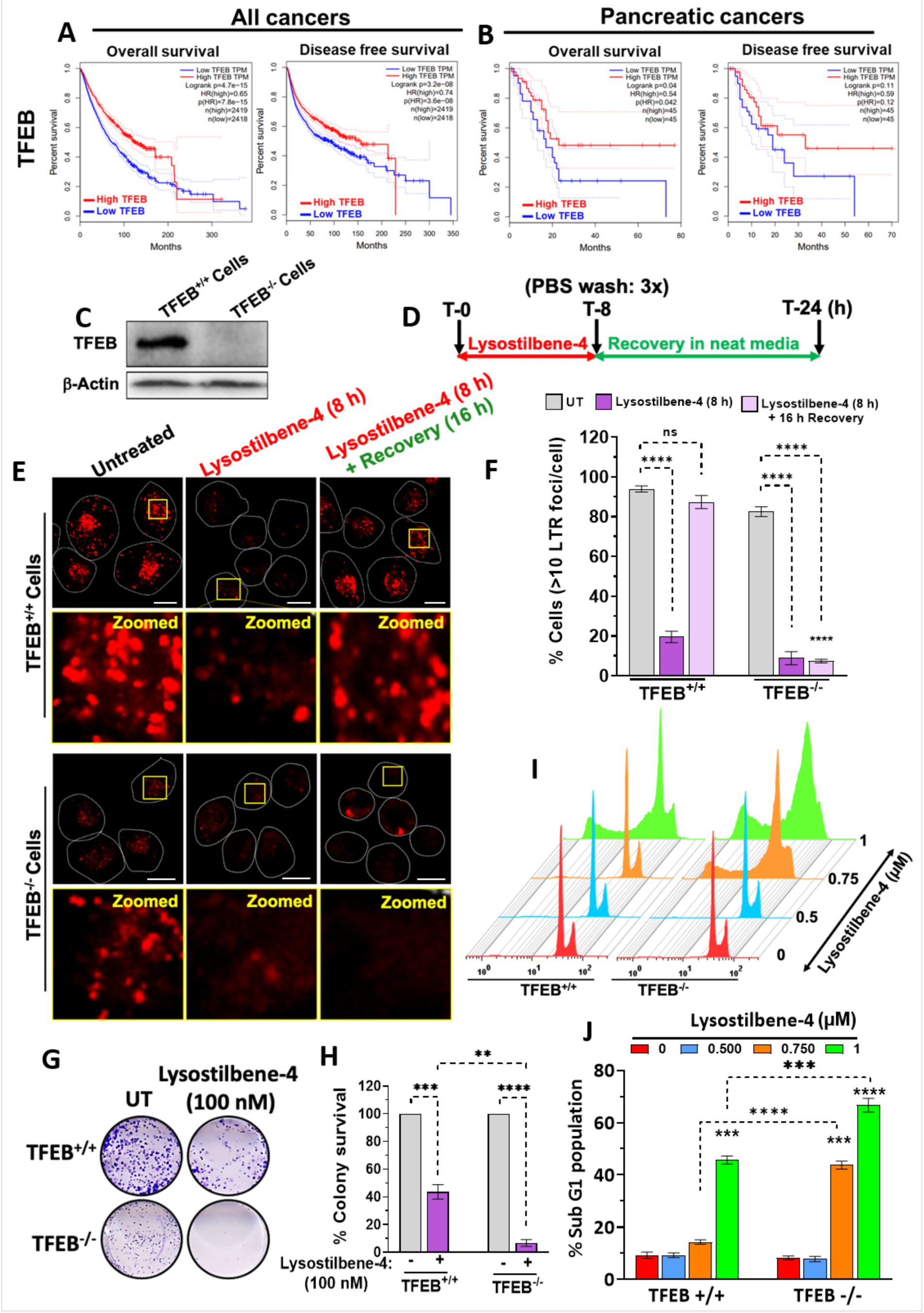
TFEB-dependent effects of Lysostilbene-4 on cancer cell survival and lysosomal function. (A, B) Kaplan-Meier survival analysis of patients with high *vs* low TFEB expression correlation with overall and disease-free survival in all types of cancers and pancreatic cancer patients. (C) Immunoblot confirmation of TFEB knockout (TFEB^−/−^) in Mia PaCa-2 cells using CRISPR-Cas9 double nickase system. β-actin served as a loading control. (D) Schematic representation of experimental design for Lysostilbene-4 treatment (8 h) followed by recovery (16 h) in Lysostilbene-4-free medium. (E) Cells were treated as per above scheme and their representative confocal images of lysosomal distribution (red, stained with LysoTracker-Red) in TFEB^+/+^ and TFEB^−/−^ cells under untreated, Lysostilbene-4 (8 h), and recovery conditions. Scale bars = 10 µm. Cells with >10 lysosomal foci per cell was quantified. Error bars represent mean ± SEM, N= 4 biological replicates. (G, H) Clonogenic survival assay for TFEB^+/+^ and TFEB^−/−^ cells treated with Lysostilbene-4 (100 nM). Representative images for colony formation and quantification of survival percentage are shown. Error bars represent mean ± SEM, N= 5 biological replicates. (I, J) Flow cytometry cell cycle distribution profiles for sub-G1 of TFEB^+/+^ and TFEB^−/−^ cells treated with increasing doses of Lysostilbene-4. Quantification of sub-G1 apoptotic cell population upon Lysostilbene-4 treatment is shown. Error bars represent mean ± SEM, N= 4 biological replicates. (ns: not significant; \*\**p* < 0.01; \*\*\**p* < 0.001; \*\*\*\**p* < 0.0001).

To test this, TFEB-knockout (TFEB⁻/⁻) MIA PaCa-2 cells were generated using CRISPR-Cas9 and validated by immunoblotting (Fig. 7C). Lysosomal integrity was assessed by Lysotracker Red in TFEB⁺/⁺ and TFEB⁻/⁻ cells following Lysostilbene-4 treatment, as per the given scheme (Fig. 7D). After 8 h of Lysostilbene-4 treatment, TFEB⁻/⁻ cells exhibited significantly greater lysosomal damage and loss of functional lysosomes compared to TFEB⁺/⁺ cells (Fig. 7E, F). Following a 16 h recovery period in Lysostilbene-4-free medium (Fig. 7E), functional lysosomal content was restored to near basal levels in TFEB⁺/⁺ cells, whereas, strikingly, recovery was almost completely abrogated in TFEB⁻/⁻ cells (Fig. 7E, F). These findings demonstrate that TFEB loss exacerbates Lysostilbene-4-induced lysosomal vulnerability by impairing compensatory lysosome biogenesis. The functional consequences of this heightened vulnerability were reflected in cell survival and apoptosis assays. TFEB⁻/⁻ pancreatic cancer cells displayed markedly reduced clonogenic survival compared with TFEB⁺/⁺ counterparts following Lysostilbene-4 treatment (Fig. 7G, H). Consistently, the IC₅₀ of Lysostilbene-4 decreased from ∼110 nM in TFEB⁺/⁺ MIA PaCa-2 cells to ∼60 nM in TFEB⁻/⁻ MIA PaCa-2 cells, reflecting heightened drug sensitivity. Flow cytometry analysis further revealed a significant increase in apoptosis (sub-G1 population) in TFEB⁻/⁻ relative to TFEB⁺/⁺ cells (Fig. 7I, J), underscoring the essential role of TFEB in supporting pancreatic cancer cell viability under lysosomal stress. Taken together, these findings demonstrate that TFEB plays a crucial role in protecting pancreatic cancer cells from lysosomal stress by promoting lysosome biogenesis and functional recovery. Lysostilbene-4 overcomes this protective mechanism by inducing sustained lysosomal damage and blocking autophagosome-lysosome assembly. Importantly, the cytotoxic effect of Lysostilbene-4 is further amplified in TFEB-deficient cells, which lack the transcriptional capacity to upregulate autophagy–lysosome (ALG) genes. This highlights a unique therapeutic vulnerability of TFEB-low pancreatic cancers, suggesting that Lysostilbene-4 may be particularly effective in patients with reduced TFEB expression.

### 2.8. Preclinical toxicity assessment of Lysostilbene-4

Although lysosome-targeting agents hold considerable promise in cancer therapy, their clinical advancement has been impeded by dose-limiting toxicities [34]. For instance, acute chloroquine and hydroxychloroquine toxicity is associated with significant mortality and morbidity *e. g.,* cardiovascular perturbations, electrolyte imbalances, and fatal dysrhythmias [35]. To investigate whether conjugating a natural stilbene pharmacophore to that of chloroquine mitigates toxicity while preserving lysosome-targeting efficacy, we assessed the safety profile of Lysostilbene-4 in vivo. First, in silico drug-likeness analysis demonstrated that Lysostilbene-4 largely falls within the threshold of Lipinski’s “rule of five,” criteria with one violation with respect to molecular weight, indicating favorable pharmacokinetic properties (Fig. 8A and methods section)

**Figure 8.**
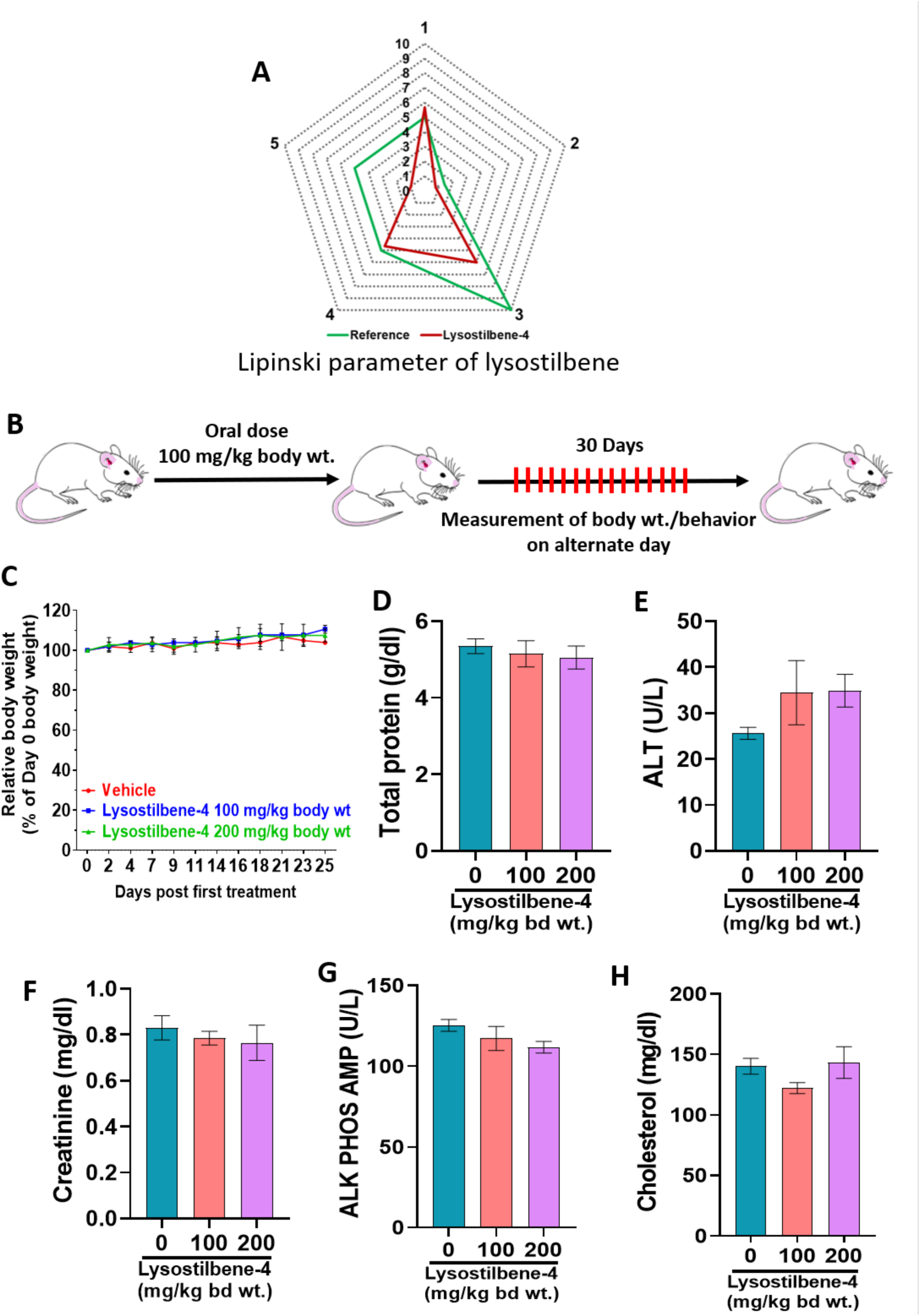
Pharmacological profiling and *in vivo* toxicity assessment of Lysostilbene-4. (A) Lipinski’s rule-based drug-likeness analysis of Lysostilbene-4. (B) Experimental scheme for *in vivo* acute toxicity assessment in mice (single oral dose: 100 or 200 mg/kg body weight) followed by monitoring of body weight and behavioural changes on alternate days for 30 days. (C) Relative body weight changes (%) over the treatment period (30 days) in vehicle and Lysostilbene-4-treated mice. Error bars represent mean ± SD, (n = 4 mice/group). (D-H) Assessment of biochemical parameters post 24 h of treatment (once an acute dose of Lysostilbene-4 of 100 or 200 mg/kg body weight) for total protein, alanine aminotransferase (ALT), creatinine, alkaline phosphatase (ALP) activity, and cholesterol levels. Error bars represent mean ± SD. (n = 4 mice/group).

To assess acute toxicity, Lysostilbene-4 was administered orally to mice at doses of 100 and 200 mg/kg body weight-doses approximately four-fold higher than the therapeutic range of its parental scaffolds, chloroquine [36] and 4,4′-dihydroxystilbene [37] (Fig. 8B). Post-treatment, animals were monitored bi-daily for one month for changes in body weight and behavior. No mortality or behavioral abnormalities were observed, and all groups exhibited a comparable gradual weight gain, indicating an absence of systemic toxicity (Fig. 8C). To further evaluate systemic safety, serum biochemistry was evaluated 24 h post-administration of the acute dose. No significant alterations were detected in total protein (Fig. 8D), alanine aminotransferase (ALT; Fig. 8E), creatinine (Fig. 8F), alkaline phosphatase (ALP; Fig. 8G), or cholesterol levels (Fig. 8H), confirming normal hepatic and renal function (Fig. 8D-H). Together, these findings demonstrate that Lysostilbene-4 is well tolerated at supra-therapeutic doses in mice, with no evidence of acute systemic or organ-specific toxicity. These results suggest that the stilbene-chloroquine hybrid design of Lysostilbene-4 confers potent lysosome-targeting activity with an improved preclinical safety profile compared to conventional lysosomotropic agents.

## 3. Discussion

Pancreatic ductal adenocarcinoma (PDAC) is an exceptionally aggressive malignancy with a dismal overall 5-year survival rate of ∼9% [38]. Despite progress in diagnostics and clinical management, therapeutic outcomes for PDAC have shown little improvement over the past five decades, underscoring the urgent need to identify vulnerable organelles and pathways as potential therapeutic entry points [39]. A defining feature of PDAC biology is its reliance on autophagy and lysosomal function to sustain growth and survival in a nutrient-deprived microenvironment[40–42]. These attributes make lysosomes an attractive therapeutic target.

Our recent review and others have emphasized diverse strategies to pharmacologically target lysosomes, employing novel chemical entities, phytochemicals, or repurposed FDA-approved drugs, either as monotherapies or in combinatorial regimens [6,43]. However, effective clinical translation of lysosome-targeting strategies has been hindered by two critical obstacles: (i) the inherent resilience of lysosomes, which activate a compensatory “3R” program (repair, removal, regeneration) in response to membrane damage [6,7], and (ii) dose-limiting toxicities associated with existing lysosomotropic drugs such as chloroquine (CQ) and hydroxychloroquine (HCQ) [23,44,45]. To overcome this, CQ is increasingly being tested in rational combination strategies with other anticancer agents [46]. Thus, there is a pressing need for next-generation lysosome-targeting agents capable of inducing sustained lysosomal damage with cancer cell selectivity and improved safety profiles. To date, no single therapeutic agent has been identified that both induces persistent lysosomal membrane permeabilization (LMP) and simultaneously suppresses the 3R compensatory programs, a prerequisite for effective lysosome-targeted sensitization of PDAC.

In this context, we hypothesized that the pharmacophore conjugation strategy may resolve above issues. No such attempts have been made to target lysosomes in cancers. The current research emphasizes the rational design of “Lysostilbenes” *via* conjugation of pharmacophores of chloroquine and natural stilbene, *trans*-4,4′-dihydroxystilbene (DHS). In this regard, one of the newly synthesized Lysostilbenes (Lysostilbene-4), showed exceptional promise in killing pancreatic cancer cells at nano molar concentration with minimal effects to non-malignant cells. Although, several CQ/HCQ variants were synthesized and shown to possess enhanced anti-cancer properties [47,48], none of these modifications are selectively sensitize malignant cells at nano molar concentration. Moreover, extensive research shows the potency of many CQ/HCQ derivatives but lacks detailed insight into the specificity for the lysosome damage and/or validated mechanism of action [49]. The differential cytotoxicity activity of lysostilbene molecules established a structure-activity relationship, emphasizing the importance of the 4-amino group, as reported earlier [18], and alkyl chain length in between CQ and DHS pharmacophore.

Mechanistically, our findings reveal a major advancement that Lysostilbene-4 is a highly selective and potent lysosomal disruptor, inducing persistent and irreversible LMP at nanomolar concentrations-unlike classical agents such as LLOMe or GPN, which act only at millimolar levels and cause transient damage [19,22]. Live-cell imaging, Galectin-3 recruitment, and cathepsin release confirmed sustained lysosomal destabilization, with minimal off-target effects on mitochondria or the ER. This lysosome-specific disruption initiates a robust apoptotic cascade [50], including cathepsin B release, BID cleavage, BCL-2 suppression, BAX upregulation, and caspase-9/3 activation. These events correlate with mitochondrial depolarization and increased sub-G1 cells, demonstrating that Lysostilbene-4 triggers a powerful lysosome-to-mitochondria apoptotic axis exceeding the effects of its parent compounds or CCCP. Autophagy is a well-established cytoprotective process in PDAC, and its inhibition represents a validated therapeutic strategy [41,46]. Our data showed another unique mechanistic feature of Lysostilbene-4, that it abolished clearance of pre-formed autophagosomes and *de novo* autophagosome assembly, suggesting a disruption of autophagic turnover at multiple levels. This multifaceted inhibition of autophagy distinguishes Lysostilbene-4 from conventional lysosomotropic agents, which typically interfere only with autophagosome-lysosome fusion. The non-involvement of the repair and removal processes suggests the possibility of activation of the regeneration/lysosome biogenesis arm as the mechanism for maintain homeostasis [7,10]. TFEB nuclear-cytoplasmic shuttling and then transcriptional activation of TFEB-mediated ALG expression is essential for lysosome biogenesis [51]. We demonstrated Lysostilbene-4 treatment led to the nuclear accumulation of TFEB in a temporal fashion *via* calcineurin-mediated dephosphorylation. Further,4X CLEAR reporter system and immunoblotting confirmed TFEB-mediated lysosome biogenesis upon Lysostilbene-4 treatment. However, despite transcriptional upregulation, functional lysosome biogenesis and autophagosome assembly were profoundly impaired in lysostilbene-4 treatment. This paradoxical phenotype-TFEB activation coupled with functional failure of the “3R” lysosomal recovery response-highlights a distinctive mechanism of Lysostilbene-4. By simultaneously inducing LMP and suppressing lysosomal repair, removal, and regeneration, Lysostilbene-4 effectively overcomes the intrinsic resilience of lysosomes that typically undermines therapeutic efficacy.

Currently, no sensitive or reliable predictive biomarkers exist to stratify pancreatic cancer patients for lysosome-targeted therapies. Fei et al. retrospectively evaluated SMAD4 expression in pancreatic adenocarcinoma specimens from patients enrolled in two clinical trials (NCT01128296, NCT01978184) testing the addition of pre-operative HCQ to neoadjuvant chemotherapy [52]. HCQ treatment was associated with improved histopathologic response and prolonged median OS in patients with SMAD4 loss, with a non-significant trend toward further OS benefit [52]. Additionally, increased peripheral blood levels of LC3-II (>51%), a surrogate marker of autophagy, correlated with extended disease-free and overall survival in pancreatic adenocarcinoma patients [53]. The functional relevance of TFEB was further clarified in our genetic studies. Analysis of TCGA datasets using KM plotter revealed that low TFEB expression correlates with poor overall and disease-free survival in pancreatic cancer, suggesting its prognostic significance. Experimentally, TFEB^−/-^ knockout PDAC cells were hypersensitive to Lysostilbene-4, exhibiting exacerbated lysosomal damage, impaired recovery, and enhanced apoptosis compared to wild-type cells. These findings identify TFEB as a key determinant of cellular resistance to lysosomal stress and establish TFEB expression status as a potential predictive biomarker for responsiveness to Lysostilbene-4. Beyond mechanistic insights, our preclinical toxicity studies provide encouraging evidence for translational potential. Lysostilbene-4 was well tolerated in mice at doses far exceeding those required for anticancer efficacy, with no observed mortality, behavioral abnormalities, or significant alterations in hepatic or renal function markers. This contrasts with the dose-limiting cardiotoxicity and retinopathy reported for CQ and HCQ [8,35]. The improved safety profile likely reflects the hybrid design, which retains lysosomotropic activity while mitigating systemic toxicity, a critical consideration for advancing lysosome-targeted agents into clinical settings.

In summary, our study establishes several key insights into lysosome-targeted therapy for pancreatic cancers: (1) Rational pharmacophore conjugation of chloroquine and stilbene yields Lysostilbene-4, which selectively sensitizes pancreatic cancers relative to non-malignant cells at nanomolar concentrations. (2) Lysostilbene-4 exhibits high lysosomal specificity with minimal off-target effects on mitochondria and endoplasmic reticulum in pancreatic cancers. (3) Lysostilbenes-4 acts uniquely through inducing LMP and abrogating lysosome repair, autophagosome (lysosphagy) and lysosome assembly (biogenesis), overcoming the inherent adaptive resilience of lysosomes, (4) TFEB-low pancreatic cancers are uniquely hypersensitized Lysostilbenes-4, highlighting TFEB as a potential predictive biomarker. (5) Importantly, Lysostilbene-4 is well tolerated at supra-therapeutic doses in mice, with no evidence of acute systemic or organ-specific toxicity. Together, these findings support Lysostilbene-4 as a next-generation lysosomotropic agent that couple potency, selectivity, and safety - key attributes for clinical translation in PDAC therapy.

## 4. Materials and Methods

Details of all materials and methods used in the current investigation are given in Supplementary Information Section. Immunoblotting assay [54, 55], Immunofluorescence assay [56, 57], Luciferase reporter assay [58], Cathepsin release assay [59], Cellular staining and Live cell imaging study [60], Kaplan Meier plotter [61], Cheminformatic analysis using Lipnski criteria [62] Preclinical toxicity study [37], were carried out as per the reported protocols with minor modifications (Supplementary Information Section).

## 5. Author contributions

B.S.P, P.D and N.C developed concepts behind the investigation. N.C and B.S.P. designed the biology experiments. B.S.P and P.D. designed the hybrid lysostilbenes while P.D. synthesized and characterized the compounds. N.C performed the biology experiments. A.G.M contributed to some of the biology experiments. N.C, P.D, AGM, M.S and B.S.P. contributed to data analysis, writing, and editing of the manuscript.

## 7. Acknowledgments

The authors express gratitude to Andrea Ballabio’s lab at TIGEM, Italy, for the generous gift of the expression plasmid EGFP-TFEB. Additionally, the authors extend their appreciation to Arnim Pause’s lab at the Goodman Cancer Research Center, Canada, for providing the 4X CLEAR luciferase reporter system.

