## Supplementary Information for "Targeting a novel chloroquine derivative to lysosomes induces massive and irreversible damage to lysosomes and suppresses autophagosomes and lysosomes assembly in cancer"

#### Materials and Methods

##### I. Chemical synthesis

All the chemicals were available commercially (Aldrich, SRL) and used without further purification. All the solvents were dried and distilled prior to use. DMF and DCM were distilled over $CaH_2$ whereas acetone and THF was dried over $K_2CO_3$ and sodium respectively. Thin-layer chromatography was done using precoated silica gel 60$F_{254}$ plates (Merck). Purifications of synthesized compounds were carried out by column chromatography using silica gel (230-400 mesh) with ethyl acetate-petroleum ether or MeOH/$CHCl_3$ mixture as the eluent. Elemental analyses were performed on Elementar Vario micro cube. Melting points were determined using Buchi M-560 melting point apparatus and are uncorrected. Infrared spectra were recorded on a Bruker Tensor II FTIR spectrophotometer and reported in wavelength numbers ($cm^{-1}$). The $^1H$ and $^{13}C$ NMR spectra were recorded with a 500 MHz Varian and 800 MHz Bruker spectrometers in $CDCl_3$, acetone-*d6* or DMSO-*d6*. Bruker TOPSPIN software was used to process the spectra. Chemical shifts are reported in ppm and calibrated relative to $CDCl_3$ ($\delta_H = 7.26$, $\delta_C = 77.00$), acetone--*d6* ($\delta_H = 2.05$, $\delta_C = 29.92$), and DMSO-*d6* ($\delta_H = 2.50$, $\delta_C = 39.51$). Peak multiplicities are abbreviated as s, singlet; bs, broad singlet; d, doublet; t, triplet; q, quartet; m, multiplet. The coupling constant *J* is reported in Hertz (Hz).

**4-azido-7-chloroquinoline (1):** To a solution of 4,7-Dichloroquinoline (1.0 g, 5.05 mmol) in 5 mL anhydrous DMF, $NaN_3$ (0.66 g, 10.1 mmol) was added, and the resulting mixture stirred under nitrogen at 100°C for 6 h. After completion (*cf.* TLC), the mixture was cooled to room temperature, DMF was removed under reduced pressure, residue was diluted with 50 mL $CH_2Cl_2$, washed with water (3 x 20 mL), dried over anhydrous $Na_2SO_4$, and evaporated to dryness. The resulting product residue was crystallized from ethyl acetate to yield the final pure product as colorless, needle-like crystals (0.91 g, 88%). Yield: 85%. m.p: 110-113°C (Lit.[1] 110-112°C). IR (neat) $\upsilon_{max}$, $cm^{-1}$: 2119, 1604, 1559, 1487, 1413, 1297, 1194, 1067, 880, 814. $^1H$ NMR (500 MHz, $CDCl_3$): δ (ppm) 8.84 (d, *J* = 5.0 Hz, 1H), 8.06-8.00 (m, 2H), 7.49 (dd, *J* = 2.0, 9.0 Hz, 1H), 7.14 (d, *J* = 5.0 Hz, 1H). $^{13}C$ NMR (125 MHz, acetone-$d_6$): δ (ppm) 152.8, 150.5, 147.1, 136.6, 128.8, 128.1, 125.0, 120.7, 110.6 (Fig. S1, S2).

**General Procedure for the synthesis of 2a-2e:** All the hydroxyquinolines (**2a**-**2e**) were synthesized as follows: A mixture of 4,7-dichloroquinoline (1.0 g, 5.05 mmol) and appropriate amino alcohol (25.25 mmol, 5 eq.) was heated at 120°C for 5h. After completion, the reaction mixture was brought to room temperature, ice was added and resulting precipitate was filtered off, washed with cold water and dried under vacuum to get the product as solid.

**2-(4-(7-chloroquinolin-4-yl)piperazin-1-yl)ethanol (2a):** Yield: 89%. White solid. m.p: 129.4-129.8°C. IR (neat) $\upsilon_{max}$, $cm^{-1}$: 3159, 1567, 1423, 1374, 1132, 1046, 866, 827. $^1H$ NMR (500 MHz, $CDCl_3$) : δ (ppm) 8.72 (d, *J* = 5.0 Hz, 1H), 8.04 (d, *J* = 1.5 Hz, 1H), 7.94 (d, *J* = 9.0 Hz, 1H), 7.42 (dd, *J* = 2.0, 9.0 Hz, 1H), 6.84 (d, *J* = 5.0 Hz, 1H), 3.70 (t, *J* = 5.5 Hz, 2H), 3.26 (s, 3H), 2.82 (bs, 4H), 2.70 (t, *J* = 5.5 Hz, 2H), 1.74 (bs, 1H). $^{13}C$ NMR (200 MHz, $CDCl_3$): δ (ppm) 156.7, 151.5, 149.6, 134.6, 128.3, 125.9, 125.0, 121.6, 108.7, 59.5, 57.9, 52.7, 51.8 (Fig. S3, S4).

**2-[(7-chloroquinolin-4-yl)amino]ethan-1-ol (2b):** Yield: 96%. Pale yellow solid. m.p.: 212-214°C (Lit.[2] 213-215°C). IR (neat) $\upsilon_{max}$, $cm^{-1}$: 3305, 1576, 1535, 1436, 1334, 1217, 1134, 1066, 798. $^1H$ NMR (500 MHz, DMSO-$d_6$) : δ (ppm) 8.37 (d, *J* = 5.0 Hz, 1H), 8.23 (d, *J* = 9.0 Hz, 1H), 7.77 (s, 1H), 7.43 (d, *J* = 9.0 Hz, 1H), 7.27-7.25 (m, 1H), 6.50 (d, *J* = 5.5 Hz, 1H), 4.92 (s, 1H), 3.64 (t, *J* = 5.0 Hz, 2H), 3.35 (t, *J* = 5.5 Hz, 2H). $^{13}C$ NMR (125 MHz, DMSO-$d_6$): δ (ppm) 151.9, 150.2, 149.1, 133.4, 127.5, 124.1, 124.0, 117.4, 98.7, 58.6, 45.0 (Fig. S5, S6).

**4-[(7-chloroquinolin-4-yl)amino]butan-1-ol (2c):** Yield: 95%. Pale yellow solid. m.p: 158-162°C. IR (neat) $\upsilon_{max}$, $cm^{-1}$: 3305, 1579, 1426, 1331, 1249, 1138, 1053, 866, 808. $^1H$ NMR (500 MHz, DMSO-$d_6$) : δ (ppm) 8.35 (d, *J* = 5.6 Hz, 1H), 8.26 (d, *J* = 8.8 Hz, 1H), 7.77 (d, *J* = 2.4 Hz, 1H), 7.40 (dd, *J* = 1.6, 8.8 Hz, 1H), 7.31 (s, 1H), 6.40 (d, *J* = 4.8 Hz, 1H), 4.91 (s, 1H), 3.45 (t, *J* = 6.4 Hz, 2H), 3.36 (t, *J* = 5.6 Hz, 2H), 1.69-1.66 (m, 2H), 1.54-1.51 (m, 2H). $^{13}C$ NMR (200 MHz, DMSO-$d_6$): δ (ppm) 151.9, 150.2, 149.1, 133.5, 127.5, 124.1, 124.0, 117.5, 98.6, 60.5, 42.4, 30.1, 24.6 (Fig. S7, S8).

**6-[(7-chloroquinolin-4-yl)amino]hexan-1-ol (2d):** Yield: 94%. Off-white solid. m.p.: 130.4-132.6°C. IR (neat) $\upsilon_{max}$, $cm^{-1}$: 3308, 1575, 1538, 1424, 1330, 1128, 1073, 896, 849. $^1H$ NMR (600 MHz, DMSO-$d_6$) : δ (ppm) 8.37 (d, *J* = 5.4 Hz, 1H), 8.26 (d, *J* = 9.0 Hz, 1H), 7.77 (s, 1H), 7.42 (d, *J* = 9.0 Hz, 1H), 7.28 (t, *J* = 4.8 Hz, 1H), 6.45-6.44 (m, 1H), 4.38 (bs, 1H), 3.39 (t merged with solvent peak, 2H), 3.24 (t, *J* = 5.4 Hz, 2H), 1.66-1.62 (m, 2H), 1.45-1.32 (m, 6H). $^{13}C$ NMR (200 MHz, DMSO-$d_6$): δ (ppm) 151.9, 150.2, 150.1, 149.1, 133.4, 127.4, 124.1, 124.0, 117.5, 98.5, 60.7, 42.5, 32.6, 27.9, 26.6, 25.4 (Fig. S9, S10).

**8-((7-chloroquinolin-4-yl)amino)octan-1-ol (2e):** Yield: 71%. White solid. m.p.: 97-101°C. IR (neat) $\upsilon_{max}$, cm$^{-1}$: 3308, 1576, 1540, 1429, 1366, 1327, 1136, 1075, 877, 801. $^1$H NMR (500 MHz, DMSO-*d$_6$*) : δ (ppm) 8.34 (d, *J* = 5.5 Hz, 1H), 8.22-8.21 (m, 1H), 7.76 (d, *J* = 2.0 Hz, 1H), 7.40 (dd, *J* = 9.0, 2.0 Hz, 1H), 7.30 (t, *J* = 5.0 Hz, 1H), 6.45 (t, *J* = 5.0 Hz 1H), 4.54 (bs, 1H), 3.36-3.33 (m, 2H), 3.25-3.22 (m, 2H), 1.65-1.59 (m, 2H), 1.37-1.24 (m, 10H). $^{13}$C NMR (125 MHz, DMSO-*d$_6$*): δ (ppm) 152.1, 150.6, 149.0, 133.9, 127.4, 124.4, 124.3, 117.7, 98.9, 61.1, 42.7, 32.7, 29.2, 29.1, 28.0, 26.9, 25.7 (Fig. S11, S12).

**General Procedure for the synthesis of 3a-3e:** All the Chloro-derivatives (**3a-3e**) from the corresponding hydroxyquinolines (**2a-2e**) were synthesized as follows: To a solution of **2a-2e** (2.0 mmol) in dry DCM and catalytic DMF, thionyl chloride (4.0 mmol) was added dropwise at 0°C and the reaction mixture was allowed to stir at room temperature for 24 h. After completion, the reaction was neutralized with a saturated solution of $NaHCO_3$ and extracted with ethyl acetate, water and brine. The combined organic phases were dried over anhydrous $Na_2SO_4$, and evaporated to dryness. The residue i.e. chlorides (**3a-3e**) were used directly for the next step without further purification.

**General Procedure for the synthesis of 4a-4e:** All the azides (**4a-4e**) were synthesized from the respective chlorides (**3a-3e**) following the procedure described for the synthesis of compound **1.**

**4-(4-(2-azidoethyl)piperazin-1-yl)-7-chloroquinoline (4a):**

Yield: 89%. Yellow viscous liquid; IR (neat) $\upsilon_{max}$, cm$^{-1:}$ 2097, 1606, 1573, 1497, 1455, 1423, 1378, 1352, 1298, 1250, 1232, 1193, 1138, 1072, 1017. $^1$H NMR (500 MHz, $CDCl_3$): δ (ppm) 8.64 (d, *J* = 4.5 Hz, 1H), 7.98 (s, 1H), 7.86 (d, *J* = 9.0 Hz, 1H), 7.35 (d, *J* = 9.0 Hz, 1H), 6.76(d, *J* = 5.0 Hz, 1H), 3.35 (q, *J* = 6.0 Hz, 2H), 3.18 (bs, 4H), 2.73 (bs, 4H), 2.67 (t, *J* = 6.0 Hz, 2H). $^{13}$C NMR (125 MHz, $CDCl_3$): δ (ppm) 156.8, 151.8, 149.9, 134.7, 128.7, 126.0, 125.1, 121.8, 108.9, 57.0, 52.9, 51.9, 48.0 (Fig. S13, S14).

**N-(2-azidoethyl)-7-chloroquinolin-4-amine (4b):**

Yield: 89%. Light yellow solid; m.p.: 139-140°C (Lit.[1] 145-147°C); IR (neat) $\upsilon_{max}$, cm$^{-1}$: 3216, 3065, 2099, 1611, 1576, 1442, 1365, 1327, 1301, 1280, 1251, 1209, 1141, 1083. $^1$H NMR (800 MHz, $CDCl_3$) : δ (ppm) 8.50 (d, *J* = 5.6 Hz, 1H), 7.93-7.92 (m, 1H), 7.78 (d, *J* = 8.8 Hz, 1H), 7.31 (dd, *J* = 8.8, 1.6 Hz, 1H), 6.40 (d, *J* = 4.8 Hz, 1H), 5.90 (s, 1H), 4.14 (s, 1H), 3.64 (t, *J* = 5.6 Hz,

2H), 3.51 (t, $J$ = 5.6 Hz, 2H). $^{13}$C NMR (200 MHz, $CDCl_3$): δ (ppm) 151.6, 149.7, 148.8, 135.3, 128.3, 125.7, 121.5, 117.3, 99.1, 49.7, 42.2 (Fig. S15, S16).

**N-(4-azidobutyl)-7-chloroquinolin-4-amine (4c):**

Yield: 89%. Pale yellow solid; m.p.: 120-122°C; IR (neat) $\upsilon_{max}$, cm$^{-1}$: 3217, 2083, 1574, 1427, 1359, 1326, 1275, 1227, 1136, 1076. $^{1}$H NMR (500 MHz, $CDCl_3$): δ (ppm) 8.47 (d, $J$ = 5.5 Hz, 1H), 7.92-7.91 (m, 1H), 7.72 (d, $J$ = 9.0 Hz, 1H), 7.31-7.29 (m, 1H), 6.36 (d, $J$ = 5.5 Hz, 1H), 5.53 (bs, 1H), 3.37-3.32 (m, 4H), 1.85-1.81 (m, 2H), 1.76-1.72 (m, 2H). $^{13}$C NMR (125 MHz, $CDCl_3$): δ (ppm) 151.7, 149.7, 148.8, 134.9, 128.4, 125.3, 121.1, 117.1, 98.9, 51.0, 42.7, 26.5, 25.9 (Fig. S17, S18).

**N-(6-azidohexyl)-7-chloroquinolin-4-amine (4d):**

Yield: 89%. White solid; m.p.: 82.4-83.1°C, IR (neat) $\upsilon_{max}$, cm$^{-1}$: 3253, 3064, 2091, 1610, 1574, 1450, 1367, 1330, 1279, 1247, 1163, 1135, 1079. $^{1}$H NMR (800 MHz, $CDCl_3$): δ (ppm) 8.48 (d, $J$ = 4.8 Hz, 1H), 7.92-7.91 (m, 1H), 7.68 (d, $J$ = 8.8 Hz, 1H), 7.29 (d, $J$ = 8.8 Hz, 1H), 6.36 (d, $J$ = 5.6 Hz, 1H), 5.28 (bs, 1H), 3.28-3.24 (m, 4H), 1.74-1.71 (m, 2H), 1.61-1.57 (m, 2H), 1.47-1.42 (m, 4H). $^{13}$C NMR (200 MHz, $CDCl_3$): δ (ppm) 151.9, 149.7, 149.0, 134.7, 128.5, 125.1, 121.0, 117.1, 98.9, 51.2, 43.0, 28.6 (x 2), 26.6, 26.4 (Fig. S19, S20).

**N-(8-azidooctyl)-7-chloroquinolin-4-amine (4e):**

Yield: 89%. Off-white solid; m.p.: 81.2-82.8°C, IR (neat) $\upsilon_{max}$, cm$^{-1}$: 2092, 1610, 1576, 1535, 1451, 1367, 1331, 1249, 1157, 1135, 1079. $^{1}$H NMR (500 MHz, $CDCl_3$): δ (ppm) 8.52 (d, $J$ = 5.0 Hz, 1H), 7.96-7.95 (m, 1H), 7.65(d, $J$ = 9.0 Hz, 1H), 7.36 (dd, $J$ = 9.0, 2.0 Hz, 1H), 6.41 (d, $J$ = 5.5 Hz, 1H), 4.99 (bs, 1H), 3.30 (q, $J$ = 7.0 Hz, 2H), 3.26 (t, $J$ = 7.0 Hz, 2H), 1.79-1.73 (m, 2H), 1.62-1.57 (m, 2H), 1.50-1.40 (m, 8H). $^{13}$C NMR (125 MHz, $CDCl_3$): δ (ppm) 151.8, 149.7, 149.0, 134.8, 128.6, 125.1, 120.9, 117.1, 99.0, 51.3, 43.2, 29.1, 28.9, 28.7 (2), 26.9, 26.5 (Fig. S21, S22).

**4-formylphenyl benzoate (5):**

To a solution of 4-hydroxybenzaldehyde **1** (6 g, 49.13 mmol) in DCM (50 ml) was added triethylamine (13.7 ml, 98.26 mmol) and stirred for half an hour. Then benzoyl chloride (6.8 ml, 58.96 mmol) was added to it slowly and stirred for 2.5 h. After completion (*cf.* by TLC) the reaction mixture was diluted with DCM (50 ml), washed with water (2 x 30 ml), brine (2 x 20 ml) and the combined organic extract was dried over anhydrous $Na_2SO_4$. The organic layer was concentrated in vacuum and the residue was purified by column chromatography (silica gel 100-200 mesh, 0-25% ethyl acetate in petroleum ether) to yield compound **2** as a white crystalline solid (9.0 g, 85%). m.p.: 92.1-92.3°C; IR (neat) $\upsilon_{max}$, $cm^{-1}$: 2722, 1736, 1693, 1590, 1258, 1203, 1054, 704. $^1H$ NMR (500 MHz, $CDCl_3$) δ 10.02 (s, 1H), 8.21 (d, *J* = 7.5 Hz, 2H), 7.98 (d, *J* = 8.0 Hz, 2H), 7.68-7.65 (m, 1H), 7.54 (t, *J* = 7.5 Hz, 2H), 7.42 (d, *J* = 8.0 Hz, 2H); $^{13}C$ NMR (125 MHz, $CDCl_3$) δ 190.9, 164.5, 155.6, 134.0, 131.2, 130.2, 128.8, 128.7, 122.5. Anal. calculated for $C_{14}H_{10}O_3$ (226.23): C, 74.33; H, 4.46; Found: C, 74.78; H, 4.49.

**(E)-4,4'-(ethene-1,2-diyl)diphenol (6):**

To a suspension of Zn (20.1 g, 309.47 mmol) in dry THF, $TiCl_4$ (16.9 ml, 154.7 mmol) was added slowly at 0°C under Argon and refluxed for 2 h. After that it was cooled to 0°C, the aldehyde **2** (5.0 g, 22.10 mmol) was added to it slowly and refluxed for 3 h. After completion (*cf.* by TLC) the reaction was diluted with ethyl acetate, 10% $K_2CO_3$ solution was added to it slowly at 0°C. Then the reaction mixture was filtered through a celite pad, the filtrate extracted with brine. Then organic layer was dried over anhydrous sodium sulphate. It was then concentrated under vacuum, the residue was purified by column chromatography (0-5% MeOH/$CHCl_3$) to get (2.15 g, 92%). m.p.: 286.5-286.6°C; IR (neat) $\upsilon_{max}$, $cm^{-1}$: 3352, 1599, 1509, 1446, 1370, 1247, 1103, 959, 829. $^1H$ NMR (500 MHz, $CDCl_3$) δ 8.55 (s, 1H), 7.39 (d, *J* = 8.5 Hz, 4H), 6.95 (s, 2H), 6.82 (d, *J* = 8.5 Hz, 4H); $^{13}C$ NMR (125 MHz, $CDCl_3$) δ 157.9, 130.5, 128.3, 126.5, 116.4. Anal. calculated for $C_{14}H_{12}O_2$ (212.25): C, 79.23; H, 5.70; Found: C, 79.49; H, 5.49.

**(E)-4-(4-(prop-2-yn-1-yloxy)styryl)phenol (7):**

To a stirred solution of 4-(4-hydroxystyryl)phenol (1.5 g, 7.08 mmol) in dry acetone, anhydrous potassium carbonate (1.47 g, 10 mmol) and propargyl bromide (1.17 g, 8.50 mmol) were added. The resultant mixture was stirred at 45°C for 18h, then the mixture was cooled and the solvent was removed under reduced pressure. The residue was diluted with ethyl acetate (50 mL) and extracted with water (3 x 10 mL). The combined organic phases were dried over anhydrous $Na_2SO_4$ and evaporated under reduced pressure. The crude product was purified by column chromatography (silica gel, 5-25% ethyl acetate/petroleum ether) to yield off-white solid product (1.06 g, 60%).

m.p.: 160-168°C; IR (neat) $\upsilon_{max}$, $cm^{-1}$: 3278, 3020, 1597, 1505, 1450, 1374, 1233, 1018, 962, 827. $^1H$ NMR (600 MHz, $CDCl_3$) δ 8.57 (s, 1H), 7.50 (d, $J$ = 9.0 Hz, 2H), 7.41 (d, $J$ = 8.4 Hz, 2H), 7.02-6.98 (m, 4H), 6.83 (d, $J$ = 8.4 Hz, 2H), 4.79 (d, $J$ = 2.4 Hz, 2H), 3.08 (t, $J$ = 2.4 Hz, 1H); $^{13}C$ NMR (200 MHz, $CDCl_3$) δ 157.9, 132.3, 130.2, 128.5, 128.3, 128.1, 127.7, 125.9, 116.4, 115.9, 79.8, 77.1, 56.3. Anal. calculated for $C_{17}H_{14}O_2$ (250.30): C, 81.58; H, 5.64; Found: C, 81.21; H, 5.86 (Fig. S23, S24).

**General procedure for the synthesis of LS (1-6):** To a stirred solution of (**7**) (161 mg, 0.64 mmol) and CQ azides **4a-e** (0.6 mmol) in 3 mL ethanol: water (9:1) were added 1M $CuSO_4$ (40 mol%) and 1M sodium ascorbate (20 mol%) and sequentially and stirred at room temperature for 16-18h. The crude product was precipitated out, it was filtered, washed with water, dried under vacuum and purified by column chromatography (silica gel, 5% methanol/chloroform-20% methanol/chloroform) to get an off-white solid product.

**(E)-4-(4-((1-(7-chloroquinolin-4-yl)-1H-1,2,3-triazol-4-yl)methoxy)styryl)phenol (LS1):**

Yield: 62%. Off-white solid; m.p.: 151-153°C; IR (neat) $\upsilon_{max}$, $cm^{-1}$: 3143, 3063, 1605, 1511, 1447, 1372, 1244, 1025, 962, 826. $^1H$ NMR (500 MHz, DMSO-$d_6$) δ 9.61 (bs, 1H), 9.15 (s, 1H), 8.96 (s, 1H), 8.29 (s, 1H), 7.99 (d, $J$ = 8.5 Hz, 1H), 7.86 (s, 1H) 7.80 (d, $J$ = 8.5 Hz, 1H), 7.51 (d, $J$ = 7.0 Hz, 2H), 7.38 (d, $J$ = 7.0 Hz, 2H), 7.09 (d, $J$ = 7.0 Hz, 2H), 7.02-6.99 (m, 2H), 6.98-6.95 (m, 2H), 6.75 (d, $J$ = 7.0 Hz, 2H), 5.32 (s, 2H); $^{13}C$ NMR (125 MHz, DMSO-$d_6$) δ 157.3, 157.1, 152.5, 149.5, 143.8, 140.5, 135.6, 130.9, 129.2, 128.5, 128.2, 127.7, 127.5, 127.0, 126.7, 125.5, 124.8, 120.5, 117.3, 115.7, 115.2, 61.0. Anal. calculated for $C_{26}H_{19}ClN_4O_2$ (454.91): C, 68.65; H, 4.21; N, 12.32; Found: C, 68.51; H, 4.40, N, 12.52 (Fig. S25, S26).

**(E)-4-(4-((1-(2-(4-(7-chloroquinolin-4-yl)piperazin-1-yl)ethyl)-1H-1,2,3-triazol-4-yl)methoxy)styryl)phenol (LS2):**

Yield: 56%. Off-white solid; m.p.: 156-158°C; IR (neat) $\upsilon_{max}$, $cm^{-1}$: 3247, 3138, 3075, 1607, 1574, 1512, 1456, 1380, 1247, 1012, 968, 831. $^1H$ NMR (500 MHz, DMSO-$d_6$) δ 9.63 (bs, 1H), 8.65-8.64 (m, 1H), 8.22 (s, 1H), 7.98 (d, $J$ = 9.0 Hz, 1H), 7.95 (s, 1H), 7.53 (d, $J$ = 9.0 Hz, 1H), 7.44 (d, $J$ = 8.0 Hz, 2H), 7.32 (d, $J$ = 8.5 Hz, 2H), 6.98 (d, $J$ = 8.0 Hz, 2H), 6.96-6.95 (m, 1H), 6.91 (d, $J$ = 8.0 Hz, 2H), 6.73 (d, $J$ = 7.5 Hz, 2H), 5.17 (s, 2H), 4.54 (t, $J$ = 6.0 Hz, 1H), 3.09 (bs, 4H), 2.86 (t, $J$ = 6.0 Hz, 2H), 2.68 (bs, 4H); $^{13}C$ NMR (125 MHz, DMSO-$d_6$) δ 157.3, 157.0, 156.4, 152.3,

149.7, 142.6, 133.8, 130.7, 128.5, 128.1, 127.7, 127.4, 126.5, 126.2, 125.9, 125.1, 124.8, 121.5, 115.7, 115.2, 109.5, 61.2, 56.9, 52.3, 51.8, 47.0. Anal. calculated for $C_{32}H_{31}ClN_6O_2$ (567.09): C, 67.78; H, 5.51; Cl, 6.25; N, 14.82. Found: C, 67.45; H, 5.86, N, 14.68 (Fig. S27, S28).

**(E)-4-(4-((1-(2-((7-chloroquinolin-4-yl)amino)ethyl)-1H-1,2,3-triazol-4-yl)methoxy)styryl)phenol (LS3):**

Yield: 68%. Off-white solid; m.p.: 174-176°C; IR (neat) $\upsilon_{max}$, $cm^{-1}$: 3268, 3068, 1610, 1515, 1453, 1372, 1247, 1052, 963, 830. $^1H$ NMR (500 MHz, DMSO-$d_6$) δ 9.63 (bs, 1H), 8.38-8.37 (m, 1H), 8.13 (d, $J$ = 9.0 Hz, 1H), 7.80-7.79 (m, 1H), 7.50-7.37 (m, 6H), 6.97-6.94 (m, 4H), 6.75 (d, $J$ = 8.5 Hz, 2H), 6.54-6.53 (m, 1H), 5.10 (s, 2H), 4.66 (t, $J$ = 6.0 Hz, 2H), 3.81-3.78 (m, 2H); $^{13}C$ NMR (125 MHz, DMSO-$d_6$) δ 157.7, 157.3, 151.8, 150.8, 150.7, 148.8, 134.1, 131.0, 128.8, 128.0, 127.7, 127.3, 126.8, 125.1, 124.8, 124.6 (x 2), 115.9, 115.4, 99.1, 61.6, 49.6, 42.0, 27.8, 25.1. Anal. calculated for $C_{28}H_{24}ClN_5O_2$ (497.98): C, 67.53; H, 4.86; N, 14.06; Found: C, 67.80; H, 5.13, N, 14.15 (Fig. S29, S30).

**(E)-4-(4-((1-(4-((7-chloroquinolin-4-yl)amino)butyl)-1H-1,2,3-triazol-4-yl)methoxy)styryl)phenol (LS4):** Yield: 71%. Off-white solid, m.p.: 142-144°C; IR (neat) $\upsilon_{max}$, $cm^{-1}$: 3284, 3069, 3019, 1603, 1510, 1449, 1371, 1240, 1019, 962, 828. $^1H$ NMR (500 MHz, DMSO-$d_6$) δ 9.62 (s, 1H), 8.38-8.37 (m, 1H), 8.26 (d, $J$ = 9.5 Hz, 2H), 7.79 (s, 1H), 7.58 (s, 1H), 7.47-7.36 (m, 5H), 6.99 (d, $J$ = 8.0 Hz, 2H), 6.95 (d, $J$ = 7.0 Hz, 2H), 6.74 (d, $J$ = 8.0 Hz, 2H), 6.52-6.51 (m, 2H), 5.12 (s, 2H), 4.43 (t, $J$ = 7.0 Hz, 2H), 3.32-3.31 (m, 2H), 1.97-1.90 (m, 2H), 1.63-1.57 (m, 2H) ; $^{13}C$ NMR (125 MHz, DMSO-$d_6$) δ 157.7, 157.3, 151.8, 150.8, 150.7, 148.8, 134.1, 131.0, 128.8, 128.0, 127.7, 127.3, 126.8, 125.1, 124.8, 124.6 (x 2), 115.9, 115.4, 99.1, 61.6, 49.6, 42.0, 27.8, 25.1. Anal. calculated for $C_{30}H_{28}ClN_5O_2$ (526.04): C, 68.50; H, 5.37; N, 13.31; Found: C, 68.76; H, 5.34, N, 13.35 (Fig. S31, S32).

**(E)-4-(4-((1-(6-((7-chloroquinolin-4-yl)amino)hexyl)-1H-1,2,3-triazol-4-yl)methoxy)styryl)phenol (LS5):**

Yield: 73%. Off-white solid; m.p.: 138-139°C; IR (neat) $\upsilon_{max}$, $cm^{-1}$: 3243, 3016, 1693, 1590, 1452, 1258, 1063, 968, 832. $^1H$ NMR (500 MHz, DMSO-$d_6$) δ 9.63 (bs, 1H), 8.36-8.35 (m, 1H), 8.24 (d, $J$ = 9.0 Hz, 1H), 7.77 (s, 1H), 7.46-7.41 (m, 3H), 7.37-7.32 (m, 3H), 6.99 (d, $J$ = 8.0 Hz, 2H), 6.95 (d, $J$ = 6.0 Hz, 2H), 6.74 (d, $J$ = 8.0 Hz, 2H), 6.45-6.43 (m, 1H), 5.12 (s, 2H), 4.35 (t, $J$ = 7.0 Hz,

1H), 3.24-3.20 (m, 2H), 1.84-1.79 (m, 2H), 1.64-1.58 (m, 2H), 1.41-1.35 (m, 2H), 1.28-1.27 (m, 2H); $^{13}C$ NMR (125 MHz, DMSO-$d_6$) δ 157.4, 157.0, 151.9, 150.5, 148.9, 142.8, 133.7, 130.7, 128.6, 127.7, 127.4, 127.3, 126.6, 124.9, 124.6, 124.3, 124.2, 117.5, 115.7, 115.1, 98.8, 61.3, 49.5, 42.4, 29.8, 27.7, 26.1, 25.8. Anal. calculated for $C_{32}H_{32}ClN_5O_2$ (554.09): C, 69.37; H, 5.82; Cl, 6.40; N, 12.64; Found: C, 69.46; H, 5.46, N, 12.87 (Fig. S33, S34).

**(E)-4-(4-((1-(6-((7-chloroquinolin-4-yl)amino)octyl)-1H-1,2,3-triazol-4-yl)methoxy)styryl)phenol (LS6):**

Yield: 64%. Off-white solid; m.p.: 156-158°C; IR (neat) $\upsilon_{max}$, $cm^{-1}$: 3263, 3115, 1611, 1512, 1456, 1363, 1244, 1091, 968, 834. $^{1}H$ NMR (500 MHz, DMSO-$d_6$) δ 9.62 (bs, 1H), 8.36-8.35 (m, 1H), 8.26 (d, $J$ = 6.5 Hz, 1H), 7.77 (s, 1H), 7.46-7.32 (m, 6H), 7.00-6.94 (m, 4H), 6.74 (d, $J$ = 8.5 Hz, 2H), 6.45-6.44 (m, 1H), 5.12 (s, 2H), 4.33 (t, $J$ = 7.0 Hz, 1H), 3.24-3.20 (m, 2H), 1.81-1.75 (m, 2H), 1.64-1.58 (m, 2H), 1.32-1.18 (m, 8H); $^{13}C$ NMR (125 MHz, DMSO-$d_6$) δ 157.4, 157.1, 151.9, 150.4, 142.8, 133.7, 130.7, 128.6, 127.7, 127.4, 127.3, 126.6, 124.9, 124.5, 124.2, 115.7, 115.1, 98.8, 61.3, 49.5, 42.5, 29.8, 28.7, 28.4, 27.8, 26.6, 25.9. Anal. calculated for $C_{34}H_{36}ClN_5O_2$ (582.15): C, 70.15; H, 6.23; Cl, 6.09; N, 12.03; Found: C, 70.49; H, 6.63, N, 12.35 (Fig. S35, S36).

**General Procedure for the synthesis of compounds (9-10):**

A mixture of Meldrum's acid (1.5 eqv.) and triethyl orthoformate (4 eqv.) was heated to reflux for 1 h. After cooling to room temperature, a solution of appropriate aniline **8a-8b** (1 eqv.) in DMF (5 ml) was added to it slowly and again refluxed for 2 h. Then the reaction was cooled to room temperature and poured into ice-cold water when the solid product got precipitated out. The solid was filtered and washed with ice-water and air dried to get the compounds **9-10** which were sufficiently pure and used directly for the next step without further purification.

**5-(((3-methoxyphenyl)amino)methylene)-2,2-dimethyl-1,3-dioxane-4,6-dione (9):**

Starting from m-anisidine **8a** (5.0 g, 40.60 mmol), Meldrum's acid (8.77 g, 60.90 mmol) and triethyl orthoformate (27.0 ml, 162.4 mmol), the product **9** was isolated as a yellow crystalline solid (11.05 g, 98%). m.p.: 107-109°C; IR (neat) $\upsilon_{max}$, $cm^{-1}$: 1719, 1674, 1629, 1590, 1489, 1444, 1415, 1329, 1270, 1201, 1160, 1089, 1020, 997. $^{1}H$ NMR (500 MHz, $CDCl_3$) δ 11.19 (d, $J$ = 13.5

Hz, 1H), 8.61 (d, $J$ = 14.5 Hz, 1H), 7.31 (t, $J$ = 8.0 Hz, 1H), 6.83-6.78 (m, 2H), 6.74 (d, $J$ = 7.0 Hz, 1H), 3.82 (s, 3H), 1.74 (s, 6H); $^{13}$C NMR (125 MHz, $CDCl_3$) δ 165.5, 163.4, 161.0, 152.5, 138.9, 130.9, 112.5, 110.1, 105.1, 103.8, 87.2, 55.5, 27.0 (Fig. S37, S38).

**2,2-dimethyl-5-(((3-(trifluoromethyl)phenyl)amino)methylene)-1,3-dioxane-4,6-dione (10):**

Starting from m-trifluoromethyl aniline **8b** (5.0 g, 31.03 mmol), Meldrum's acid (6.70 g, 46.54 mmol) and triethyl orthoformate (20.6 ml, 124.12 mmol), the product **10** was isolated as a yellow crystalline solid (8.67 g, 93%). m.p.: 154-156°C; IR (neat) $\upsilon_{max}$, cm$^{-1}$: 1735, 1679, 1620, 1583, 1415, 1303, 1265, 1121, 1011. $^{1}$H NMR (500 MHz, $CDCl_3$) δ 11.32 (d, $J$ = 13.5 Hz, 2H), 8.66 (d, $J$ = 14.0 Hz, 2H), 7.60-7.57 (m, 1H), 7.54-7.51 (m, 2H), 7.45 (d, $J$ = 8.0 Hz, 1H), 1.76 (s, 6H); $^{13}$C NMR (125 MHz, $CDCl_3$) δ 165.4, 163.2, 152.4, 138.4, 132.7 (q, $^{2}J_{C\text{-}F}$ = 33.0 Hz), 130.8, 123.3 (q, $^{1}J_{C\text{-}F}$ = 271.0 Hz), 123.2 (q, $^{3}J_{C\text{-}F}$ = 3.7 Hz), 121.1 (d, $^{4}J_{C\text{-}F}$ = 1.0 Hz), 114.9 (d, $^{3}J_{C\text{-}F}$ = 3.8 Hz), 105.5, 88.4, 27.1 (Fig. S39, S40).

**General Procedure for the synthesis of compound (11-12):**

A solution of compounds **9-10** in diphenyl ether (2 ml/mmol of **9-10**) was heated to 250°C under argon atmosphere for 45 min-1 h. Then it was allowed to cool and diluted with equal amount of n-hexane when the solid product got precipitated. The residue was separated by filtration followed by washing twice with n-hexane. Finally dried in vacuum to get compounds **11-12**.

**7-methoxyquinolin-4-ol (11):** Starting from **5-(((3-methoxyphenyl)amino)methylene)-2,2-dimethyl-1,3-dioxane-4,6-dione (9)** (8.0 g, 28.88 mmol) in diphenylether (58 ml), the compound **11** as a mixture of 2-regioisomers **11a** and **11b**, was isolated as brown solid (3.5 g, 69%). IR (neat) $\upsilon_{max}$, cm$^{-1}$: 1615, 1554, 1469, 1258, 1218, 1165, 1020. $^{1}$H NMR (500 MHz, $CDCl_3$) δ 7.99 (d, $J$ = 9.0 Hz, 1H), 7.87 (d, $J$ = 9.0 Hz, 1H), 6.96-6.93 (m, 2H), 6.03 (d, $J$ = 6.0 Hz, 1H), 3.85 (s, 3H); $^{13}$C NMR (125 MHz, $CDCl_3$) δ 176.9, 176.8, 162.3, 162.2, 142.1, 142.0, 139.8, 139.7, 127.1, 127.0, 120.1, 120.0, 114.1, 114.0, 108.6, 99.4, 99.2, 55.8, 55.7 (Fig. S41, S42).

**7-(trifluoromethyl)quinolin-4-ol (12):** Starting from **2,2-dimethyl-5-(((3-(trifluoromethyl)phenyl)amino)methylene)-1,3-dioxane-4,6-dione (10)** (3.3 g, 10.46 mmol) in diphenyl ether (20.9 ml), compound **12** as a mixture of **12a** and **12b** was isolated as a brown solid (1.76 g, 79 %). IR (neat) $\upsilon_{max}$, cm$^{-1}$: 1643, 1600, 1568, 1474, 1429, 1191, 1116, 1053. $^{1}$H NMR (500 MHz, $CDCl_3$) δ 7.99 (d, $J$ = 9.0 Hz, 1H), 7.87 (d, $J$ = 9.0 Hz, 1H), 6.96-6.93 (m, 2H), 6.03

(d, $J$ = 6.0 Hz, 1H), 3.85 (s, 3H); $^{13}$C NMR (125 MHz, $CDCl_3$) δ 17.8, 141.0, 139.8, 131.8 (q, $^{2}J_{C-F}$ = 32.0 Hz), 127.8, 127.1, 123.9 (q, $^{1}J_{C-F}$ = 271.2 Hz), 119.1 (q, $^{3}J_{C-F}$ = 3.0 Hz), 116.2, 110.1 (Fig. S43, S44).

**General Procedure for the synthesis of 13-14:**

To a solution of compound **11** (2.75 g, 15.69 mmol) or compound **12** (2.05 g, 9.62 mmol) in DMF (2 ml), $POCl_3$ (2.9 ml, 31.38 mmol) for **11** or $POCl_3$ (1.8 ml, 19.24 mmol) for **12** was added dropwise at 0°C and then the reaction mixture was stirred at 100°C for 3 h. After completion, crushed ice was added and stirred for half an hour. The precipitate formed was collected by filtration, washed with saturated $NaHCO_3$ solution, dried under vacuum. The solid mixture was purified by column chromatography using 0-25% EA/PE to get **13** (66%) or compound **14** (74%) with **13a** and **14a** as the major product.

**4-chloro-7-methoxyquinoline (13a):**

Yield: 1.28 g, 64%; White crystalline solid; m.p.: 80-86°C; IR (neat) $\upsilon_{max}$, cm$^{-1}$: 3091, 3065, 1560, 1495, 1465, 1436, 1342, 1304, 1256, 1225, 1194, 1159, 1123, 1018. $^{1}$H NMR (500 MHz, $CDCl_3$) δ 8.69 (d, $J$ = 4.5 Hz, 1H), 8.44 (s, 1H), 8.38 (d, $J$ = 9.5 Hz, 1H), 8.11 (d, $J$ = 9.5 Hz, 1H), 7.43 (d, $J$ = 2.5 Hz, 1H), 7.35 (d, $J$ = 5.0 Hz, 1H), 7.29 (dd, $J$ = 9.5, 2.5 Hz, 1H), 3.97 (s, 3H); $^{13}$C NMR (125 MHz, $CDCl_3$) δ 161.3, 150.9, 150.2, 142.4, 125.3, 121.6, 120.8, 119.2, 107.5, 55.6 (Fig. S45, S46).

**4-chloro-7-(trifluoromethyl)quinoline (14a):**

Yield: 1.36 g, 83%. White crystalline solid. m.p.: 69-71°C. IR (neat) $\upsilon_{max}$, cm$^{-1}$: 3092, 1587, 1559, 1451, 1376, 1323, 1284, 1194, 1148, 1112, 1059. $^{1}$H NMR (500 MHz, $CDCl_3$): δ (ppm) 8.89 (d, $J$ = 4.5 Hz, 1H), 8.40 (d, $J$ = 9.0 Hz, 1H), 8.38 (d, $J$= 9.0 Hz, 1H), 7.84-7.82 (m, 1H), 7.62 (d, $J$ = 5.0 Hz, 1H). $^{13}$C NMR (125 MHz, $CDCl_3$): δ (ppm) 151.2, 148.2, 142.6, 138.2, 132.1 ($^{2}J_{C-F}$ = 32.7 Hz), 128.0, 127.7 (q, $^{3}J_{C-F}$ = 4.2 Hz), 125.6, 123.6 (q, $^{1}J_{C-F}$ = 271.3 Hz), 123.2 ($^{3}J_{C-F}$ = 2.5 Hz), 122.9 (Fig. S47, S48).

**General Procedure for the synthesis of 15-16:** Both the Chloro derivatives (**13a-14a;** 1 eq.) were treated with 4-amino-1-butanol (25.25 mmol, 5 eq.) and heated at 120°C for 5h. After completion, the reaction mixture was brought to room temperature, ice was added and resulting precipitate was filtered off, washed with cold water and dried under vacuum to get the product **15/ 16** as solid.

**4-((7-methoxyquinolin-4-yl)amino)butan-1-ol (15) :**

Yield: 90%. Off-white solid. m.p.: 128-130°C. IR (neat) $\upsilon_{max}$, $cm^{-1}$: 3306, 3078, 1579, 1540, 1434, 1339, 1296, 1233, 1136, 1074. $^{1}$H NMR (500 MHz, acetone-$d_6$): δ (ppm) 8.34 (d, $J$ = 5.0 Hz, 1H), 8.04 (d, $J$ = 9.5 Hz, 1H), 7.25 (d, $J$ = 1.5 Hz, 1H), 7.00-6.98 (m, 1H), 6.64 (bs, 1H), 6.38 (d, $J$ = 5.5 Hz, 1H), 3.88 (s, 3H), 3.62 (t, $J$ = 6.5 Hz, 2H), 3.37-3.33 (m, 2H), 1.86-1.80 (m, 2H), 1.70-1.64 (m, 2H). $^{13}$C NMR (125 MHz, acetone-$d_6$): δ (ppm) 161.1, 151.9, 151.5, 151.4, 123.1, 116.8, 114.5, 108.7, 98.2, 62.1, 55.6, 43.7, 31.2, 26.0 (Fig. S49, S50).

**4-((7-(trifluoromethyl)quinolin-4-yl)amino)butan-1-ol (16):**

Yield: 93%. Pale-yellow solid. m.p.: 143-144°C. IR (neat) $\upsilon_{max}$, $cm^{-1}$: 3321, 3081, 1590, 1549, 1430, 1371, 1326, 1284, 1268, 1156, 1069. $^{1}$H NMR (500 MHz, DMSO-$d_6$): δ (ppm) 8.46-8.40 (m, 2H), 8.06 (s, 1H), 7.66-7.64 (m, 1H), 7.48-7.47 (m, 1H), 6.59-6.58 (m, 1H), 4.70 (t, $J$ = 4.5 Hz, 1H), 3.46-3.42 (m, 2H), 3.31-3.27 (m, 2H), 1.71-1.65 (m, 2H), 1.55-1.50 (m, 2H). $^{13}$C NMR (125 MHz, acetone-$d_6$): δ (ppm) 153.2, 151.4, 151.3, 148.7, 130.8 ($^{2}J_{C\text{-}F}$ = 32.0 Hz), 127.5 ($^{3}J_{C\text{-}F}$ = 4.3 Hz), 126.4, 124.0, 123.1 ($^{1}J_{C\text{-}F}$ = 280.4 Hz), 120.0 ($^{3}J_{C\text{-}F}$ = 3.2 Hz), 100.7, 62.1, 43.8, 31.0, 25.7 (Fig. S51, S52).

**General Procedure for the synthesis of 17-18:**

All the amino alcohols (**15-16**) were converted to their respective chlorides following the procedure described for the synthesis of compounds **3a-3e**. These chlorides were finally converted to their respective azides following the procedure described for the synthesis of compound **1.**

**N-(4-azidobutyl)-7-methoxyquinolin-4-amine (17):**

Yield: 64% (yield of two steps). Off-white solid. m.p.: 119-120°C. IR (neat) $\upsilon_{max}$, $cm^{-1}$: 3020, 2099, 1553, 1466, 1368, 1298, 1242, 1124, 1085, 1035. $^{1}$H NMR (500 MHz, $CDCl_3$): δ (ppm) 8.29-8.28 (m, 1H), 7.79 (d, $J$ = 9.5 Hz, 1H), 7.23 (s, 1H), 6.98 (d, $J$ = 8.5 Hz, 1H), 6.27 (bs, 1H), 6.24-6.23 (m, 1H), 3.85 (s, 3H), 3.34-3.32 (m, 4H), 1.83-1.78 (m, 2H), 1.73-1.68 (m, 2H). $^{13}$C NMR (125 MHz, $CDCl_3$): δ (ppm) 160.9, 151.2, 148.3, 147.5, 121.7, 117.2, 112.6, 105.8, 97.2, 55.4, 51.0, 42.7, 26.4, 25.9 (Fig. S53, S54).

**N-(4-azidobutyl)-7-(trifluoromethyl)quinolin-4-amine (18):**

Yield: 72% (yield of two steps). Pale-yellow solid. m.p.: 110.7-111°C. IR (neat) $\upsilon_{max}$, $cm^{-1}$: 3074, 2096, 1590, 1552, 1431, 1370, 1323, 1275, 1162, 1111, 1066. $^{1}H$ NMR (500 MHz, $CDCl_3$): δ (ppm) 8.56-8.55 (m, 1H), 8.23 (s, 1H), 7.89-7.88 (m, 1H), 7.54-7.52 (m, 1H), 6.47-6.46 (m, 1H), 5.48 (bs, 1H), 5.74-5.70 (m, 1H), 3.36-3.33 (m, 4H), 1.87-1.81 (m, 2H), 1.76-1.71 (m, 2H). $^{13}C$ NMR (125 MHz, $CDCl_3$): δ (ppm) 152.1, 149.5, 147.5, 130.8 (q, $^{2}J_{CF}$ = 32.5 Hz), 127.3, 127.1, 123.9 (q, $^{1}J_{CF}$ = 270.8 Hz), 120.9, 120.4, 120.0 (q, $^{2}J_{CF}$ = 2.7 Hz), 116.8, 114.5, 108.7, 100.0, 51.0, 42.7, 26.4, 25.9 (Fig. S55, S56).

**General Procedure for the synthesis of compounds LS-(7-8):**

Compounds **LS-7** and **LS-8** were synthesized following the same procedure as compounds **LS-(1-6).**

**(E)-4-(4-((1-(((7-methoxyquinolin-4-yl)amino)methyl)-1H-1,2,3-triazol-4-yl)methoxy)styryl)phenol (LS7):**

Yield: 56%. Off-white solid. m.p.: 249-251°C. IR (neat) $\upsilon_{max}$, $cm^{-1}$: 3266, 3072, 1738, 1682, 1619, 1544, 1517 1464, 1372, 1073, 970. $^{1}H$ NMR (500 MHz, DMSO-$d_6$): δ (ppm) 9.67 (bs, 1H), 8.51 (bs, 1H), 8.33-8.31 (m, 1H), 8.29-8.27 (m ,1H), 7.45-7.44 (m, 2H), 7.37-7.35 (m, 2H), 7.22-7.19 (m, 2H), 6.99-6.97 (m, 2H), 6.94 (d, $J$ = 6.5 Hz, 2H), 6.74 (d, $J$ = 6.5 Hz, 2H), 6.61 (d, $J$ = 8.0 Hz, 1H), 5.11 (s, 2H), 4.42 (t, $J$ = 7.0 Hz, 2H), 3.88 (s, 3H), 3.44-3.40 (m, 2H), 1.96-1.90 (m, 2H), 1.64-1.58 (m, 2H). $^{13}C$ NMR (125 MHz, DMSO-$d_6$): δ (ppm) 160.5, 157.4, 157.0, 152.6, 151.3, 149.3, 142.9, 130.7, 128.6, 127.7, 127.5, 126.6, 124.9, 124.6, 123.6, 116.1, 115.7, 115.2, 106.3, 61.3, 55.5, 49.3, 41.9, 27.5, 25.0. Anal. calculated for $C_{31}H_{31}N_5O_3$ (521.62): C, 71.38; H, 5.99; N, 13.43; Found: C, 72.69; H, 6.06, N, 13.23 (Fig. S57, S58).

**(E)-4-(4-((1-(((7-(trifluoromethyl)quinolin-4-yl)amino)methyl)-1H-1,2,3-triazol-4-yl)methoxy)styryl)phenol (LS8):**

Yield: 76%. Off-white solid. m.p.: 236-238°C. IR (neat) $\upsilon_{max}$, $cm^{-1}$: 3261, 1585, 1544, 1469, 1441, 1375, 1329, 1241, 1161, 1118, 1069, 965. $^{1}H$ NMR (500 MHz, DMSO-$d_6$): δ (ppm) 9.64 (s, 1H), 8.47-8.42 (m, 2H), 8.07 (s, 1H), 7.66 (d, $J$ = 8.5 Hz, 1H), 7.49 (t, $J$ = 4.0 Hz, 1H), 7.45 (d, $J$ = 8.5 Hz, 2H), 7.36 (d, $J$ = 8.5 Hz, 2H), 7.00-6.94 (m, 4H), 6.74 (d, $J$ = 8.5 Hz, 2H), 6.57 (d, $J$ = 5.5 Hz,

1H), 5.11 (s, 2H), 4.43 (t, $J$ = 6.5 Hz, 2H), 3.35-3.30 (m, 2H), 1.99-1.93 (m, 2H), 1.66-1.60(m, 2H). $^{13}$C NMR (125 MHz, DMSO-$d_6$): δ (ppm) 157.4, 157.0, 152.3, 150.2, 147.3, 142.9, 130.7, 129.4 (q, $^2J_{CF}$ = 31.8 Hz), 127.6 (q, $^1J_{CF}$ = 269.2 Hz), 126.1 (q, $^4J_{CF}$ = 4.3 Hz), 125.4, 124.9, 124.6, 124.1, 123.2, 121.0, 119.2 (q, $^3J_{CF}$ = 8.2 Hz), 115.7, 115.1, 99.9, 61.3, 49.3, 41.9, 27.5, 24.7. Anal. calculated for $C_{31}H_{28}F_3N_5O_2$ (559.59): C, 66.54; H, 5.04; N, 12.52; Found: C, 66.86; H, 5.27, N, 12.78 (Fig. S59, S60).

#### II. Biological studies

**Materials**: Unless otherwise noted, all the chemicals were purchased from Sigma-Aldrich, USA. Dulbecco's modified Eagle's medium (DMEM), Trypsin, fetal bovine serum (FBS) and antibiotics-antimycotic solution from Gibco Life Technologies, USA. Following antibodies were used in the current study: galectin-3 (# sc-32790, SCBT), LAMP2 (# ab25631, Abcam), LC3 (# 12741, CST), TFEB (# 4240, CST), Caspase 3 (# 9662, CST), Caspase-9 (# sc-81663, SCBT), β-actin (# ab8227, Abcam), BID (# 2003, CST), BAX (# sc-493, SCBT), BCL-2 (# B3170, Sigma), p-TFEB S211 (# 37681, CST). Secondary antibodies tagged with Alexa fluor 488/594 were from Jackson Immunoreseach, USA. Secondary antibody HRP conjugated rabbit IgG and mouse IgG for immunoblotting were procured from CST. Lipofectamine 2000, Lipofectamine 3000 and Prolong Diamond anti-fade reagent were procured from Invitrogen (Carlsbad, CA). Protease inhibitor, phosphatase inhibitor cocktails (PhosStop) and western blotting substrate and were from Roche, Switzerland. Cellular organelles staining dyes as Lysotracker red-DND (# L7528), Lysotracker red-DND (# L7526), Mitotracker Red CM-H2ROX (# M7513) and ER-Tracker green (# E34251) were from Invitrogen (Carlsbad, CA), while acridine orange (# A6014) was from Sigma-Aldrich, USA. The chemicals MTT reagent (# M5655, Sigma), digitonin (# D141, Sigma), LLOMe (# L7393, Sigma), Chloroquine (# C6628), Bafilomycin A1 (# 11038, Sigma), G418 sulfate (# 345810, Merck), puromycin (# P8833), Cyclosporin A (# C3662, Sigma) and Propidium Iodide (# P4170) were procured from Sigma-Aldrich, USA. Chambered Coverglass for confocal microscopy (# 155382 and 155409) was procured from Thermo Fisher Scientific, USA. Plasmids mRFP-EGFP-Gal3 (tf-Gal3) (# 64149) and mRFP-EGFP-LC3 (tf-LC3) (#21074) were from Addgene. The expression plasmid EGFP-TFEB was a generous gift from Andrea Ballabio`s lab*, TIGEM, Italy*. The pRL Renilla Luciferase Control Reporter Vector (# E2261) and Dual-Luciferase Reporter Assay System (# E1960) were purchased from Promega, Madison, Wisconsin,

United States. The 4X CLEAR luciferase reporter system was a generous gift from Arnim Pause`s lab, Goodman Cancer Research Center, Canada.

**Cell culture**: Pancreatic cancer cell lines such as MIA PaCa-2 and PANC-1 were procured from the European Collection of Authenticated Cell Cultures (ECACC). Non-cancerous cell lines (Vero and MCF10A) were from the American Type Culture Collection (ATCC). The cell lines were cultured as per the instructions mentioned in ECACC and ATCC. Cancer cells were maintained routinely in the suggested medium supplemented with 10% FBS and 1% of antibiotics solutions (penicillin-streptomycin and amphotericin) in an incubator (37°C; 95% relative humidity; 5% $CO_2$).

**Generation of the knockout cell line.** The CRISPR-CAS9 double nickase plasmid for TFEB (# sc-401388) was purchased from Santa Cruz Biotechnology (Dallas, TX, USA). Control double nickase was used for transfection control. Exponentially grown MIA PaCa-2 cells at around 80% confluency were transfected using Lipofectamine 2000 and CRISPR-Cas9 plasmid as per the manufacturer`s protocol. Further, cells were selected in puromycin and the efficiency of stable knockout was assessed by western blotting.

**Generation of EGFP-TFEB, mRFP-EGFP-Gal3 (tf-Gal3) and mRFP-EGFP-LC3 (tf-LC3) expressing MIA PaCa-2 cells.** Exponentially growing cells at the sub-confluent stage were transfected using lipofectamine 3000 and plasmids. Expression-positive cells were selected in G418 sulfate and expression was assessed by live cell confocal microscopy.

**Clonogenic survival assay.** The cells were seeded at a density of 750 cells/well in 6-well plate and allowed to grow overnight. The cells were treated with different concentrations of test compounds and incubated for 8-10 days. The colonies were fixed with chilled methanol and stained with 0.5% crystal violet solution in 1:1 water-methanol. The stained colonies were counted and the surviving fraction in treatment was determined using 100% survival reference in nontreated control.

**Cell viability assay.** The cells ($3 \times 10^3$ cells/well) were seeded in 96-well plate and allowed to attach and grow for 24 hours. The cells were incubated with different concentrations of the compounds for 72 hours. Metabolically active and viable cells were assessed by the MTT assay. MTT solution (0.5 mg/mL) was added and incubated for 2 h. Later, the formazan crystals were solubilized in DMSO and absorbance was recorded at 570 nm in a BMG-Polestar multi-plate reader.

**Analysis of cell death by sub-G1 assay.** Briefly, overnight-grown cells (8.0 x $10^4$ cells/well) in 6-well plates, were treated at different concentrations as indicated. Post-treatment, cells were trypsinized and washed once with ice-cold PBS. The cells were resuspended in Na-citrate hypotonic buffer (0.1% Triton X-100; 0.1% sodium citrate) containing RNase A (50 µg/mL) and propidium iodide (25 µg/mL) and incubated in the dark for 15 min. Further, at least 2.5 x $10^4$ cells were acquired by the Partec CyFlo flow cytometer. For quantification of the Sub-G1 population, cells with <2n DNA amount were quantified using offline FlowJo software.

**Immunoblotting assay.** Western blotting was employed as per earlier protocol with minor modification [54, 55] to analyze the change in protein expression under the treatment of Lysostilbene-4. Briefly, MIA PaCa-2 cells (6 x $10^5$ cells/dish) were seeded in 60 mm culture dishes. After 24 h, cells were treated with indicated concentrations and time points and then cells were harvested by scraping. Later, cells were lysed using RIPA lysis buffer having protease inhibiter and phosphatase inhibiter cocktails for 30 min on ice and vortexed intermittently. The lysate was then centrifuged at 12000 rpm at 4°C for 20 min. Later, supernatant was collected, a fraction was used for protein concentration estimation via Bradford assay and lamelle buffer was added to the remaining sample and boiled at 95°C for 10 minutes. Later, 30-40 µg protein per sample was loaded and resolved in SDS-PAGE and then blotted on PVDF membrane. Afterwards, blocking was done with 2% BSA in PBST (PBS containing 0.1% Tween-20), and then the interested protein was probed with primary antibody in 1% BSA in PBST at 4°C for overnight. Blots were washed three times with PBST each for 7 minutes and HRP-conjugated secondary antibody for 3 h at room temperature. Blots were washed again as mentioned above, and then developed using Lumi-Light Plus western blotting substrate. Protein bands were detected using GeneSys software in the Syngene G-Box instrument. Further, protein level was determined by densitometric analysis using ImageJ offline software.

**Immunofluorescence assay.** Immunofluorescence assays were conducted with earlier published protocol with minor modification [56, 57]. Briefly, MIA PaCa-2 (1.5 x $10^5$) cells were seeded per well in 6-well plate and allowed to adhere and grow for 24 h. The cells were treated with different test molecules at various time points as indicated in the figures. Post-treatment, cells were washed with ice-cold PBS and fixed with 3% PFA at 4°C for 15 minutes. Later, cells were washed with PBS and permeabilized with 0.25% Triton X-100 in PBS at 4°C for 15 minutes. Further, blocking was done by incubating cells with 5% BSA in PBS for 1 h. After that, the primary antibody was added in 2.5% BSA in PBS for 3 h. After washing cells thrice with PBS each for 10 minutes, cells were incubated with secondary antibodies conjugated with Alexa fluor 488/594 for 3 h at room temperature. Cells were again washed three times; coverslips were air-dried and mounted on a glass slide using 80% glycerol containing Hoechst 33258. Samples were then acquired with laser scanning confocal microscope (LSM780, Carl Zeiss, Germany). Image analysis was done using Zeiss Zen offline software.

**Luciferase reporter assay.** Experiments were done as per earlier published protocol with few changes [58]. Cells were seeded in 24-well plates and grown overnight. Cells were co-transfected with 4X CLEAR luciferase reporter plasmid and pRL Renilla Luciferase Control Reporter plasmid in 100:1 quantity. The individual plasmid transfection was also done to rule out any plasmid cross-interaction affecting transfection efficiency. Post-24-hour transfection, cells were maintained in drug-free complete media for another 24 h to minimize the effect of the transfection reagent on the reporter activity. Cells were treated with the indicated treatment and then cell lysis was done at 4°C for 10 minutes using the provided lysis solution. The supernatant was transferred to the transparent bottom in 96-well black plate and further processing was done as per the manufacturer`s protocol. The luminescence intensity was recorded using a microplate Illuminometer feature of BMG Labtech, POLARstar Omega. Firefly luciferase readings were normalized to internal control Renilla luciferase readings for each respective well.

**Cathepsin activity assay.** Live cell confocal microscopy was performed to assess cathepsin B activity [5]. Briefly, 5 x $10^4$ cells/ well were seeded in confocal microscopy compatible chambered Coverglass and allowed to grow overnight. Cells were treated with Lysostilbene-4 and LLOMe as indicated. Then, cells were incubated with magic red cathepsin B substrate as per instructions mentioned on their website. The cells were observed and images were acquired under 40x objective lens of the confocal microscope. The data was analyzed using Zeiss Zen Blue software.

**Cathepsin release assay.** The cells were seeded in 24-well plate and allowed to attach overnight. Cells were treated with Lysostilbene-4 as mentioned in the figure. Post-treatment, media was discarded and cells were washed once with PBS and further processed with a few modifications to the published protocol for this assay [59]. The limited cell permeabilization was done by incubating cells with 20 µg/mL of digitonin in 100 mM phosphate buffer (pH 6.0) for 10 min on ice in rocking conditions. The supernatant was then collected and transferred to a transparent bottom 96-well black plate on ice. Then, a fluorogenic substrate for cathepsin B, magic red solution was added to the wells. Immediately, the fluorescence intensity was recorded using a multiwell microplate fluorescence feature of BMG Labtech, POLARstar Omega. The data were plotted using GraphPad prism after normalization with control sample data.

**Cellular staining** [60]**.** (a) For confocal microscopy, 2 x $10^5$ cells were seeded per 30 mm confocal culture dishes and grown overnight. Cells were treated for mentioned concentration and time points. Post-treatment, drug-containing media was discarded and then cells were incubated for 5 min with cellular organelle staining dyes as LysoTracker Red (100 nM) or LysoTracker Green (100 nM) or acridine orange (10 µM). The samples were acquired as per the manufacturer's suggested excitation laser and emission filter using an LSM780 confocal microscope. Fluorescence intensity quantification was done using Zeiss Zen Blue offline software. (b) For flow cytometry, 7.5 x $10^5$ cells/well were seeded in a 12-well plate and treated with test molecules as indicated after 24 h. Post-treatment, drug-containing media was discarded and cells were incubated with dyes as Lysotracker red (100 nM), Mitotracker Red CM-H2ROX (300 nM) and ER-Tracker green (300 nM) as per the manufacturer's protocol. The cells were collected by trypsinization and washed once with PBS. Then, at least 2.5 x $10^4$ events were recorded using Partec CyFlow flow cytometer. The data was then analyzed by gating population or change in MFI (Mean Fluorescence Intensity) using FlowJo software.

**Live cell imaging** [60]**.** Briefly, EGFP-TFEB or tf-Gal3 or tf-LC3 MIA PaCa-2 stable cells were seeded (5 x $10^5$ cells/well) in chambered coverglass compatible with live cell imaging. Cells were treated after 24 h of seeding for mentioned treatments. Live cell imaging acquisition was done at the indicated time points using the confocal microscope. During acquisition, live cell conditions (5% $CO_2$; 37°C and 95% humidity) were maintained using an in-built cell incubator chamber. Quantification and analysis of acquired images were done as mentioned earlier with Zeiss Zen Blue and Black software.

**KM plots.** The Kaplan Meier (KM) plots were plotted using freely accessible online platform KM plotter (https://www.kmplot.com/). The KM Plots were used for assessing the correlation of TFEB protein with survival outcome (overall and diseases free survival) in all cancer vs pancreatic cancer [61].

**Cheminformatics analysis**. Cheminformatics analysis of Lysostilbene-4, using Lipinski criteria. MR: Molar refractivity; HBA: Hydrogen Bond Acceptors; CLogP: calculated Log P; HBD: Hydrogen Bond Donors; MW: Molecular Weight. Please refer to experimental section for detailed analytical method and models employed [62].

| | **MW (1/100)** | **MR (1/100)** | **HBA** | **CLogP** | **HBD** | **Lipinski** |
|---|---|---|---|---|---|---|
| **Threshold** | <5 | 0.4-1.3 | <10 | <5 | <5 | ------------- |
| **Lysostilbene-4** | 5.67 | 0.79 | 6 | 4.64 | 1 | 1 violation (MW >500) |

**Toxicity study in mice** [37]**:** Male BALB/c mice (7-8-week-old, 23-26 g average body weight) were obtained from the BARC animal breeding facility. The approval of the use of animals for the current study was obtained from the Institutional Animal Ethics Committee (IAEC) of Bhabha Atomic Research Center (approval number BAEC/01/2023). Animals were maintained as per the institutional guidelines and randomly segregated into 3 groups (n = 4 in each group) at the start of the experiment. Mice were given a single oral gavage of vehicle or Lysostilbene-4 (100 and 200 mg/kg body wt). Post-acute treatment, body weight, behavior and stool texture were monitored and recorded every alternate day for 30 days. Then, animals were sacrificed. In another experiment, mice were treated as mentioned above for 24 h. Post-treatment, animals were sacrificed and blood was collected by cardiac puncture using a syringe needle. The blood serum was subjected to various biochemical parameters such as Alkaline phosphate AMP, Glucose PAP, Cholesterol, creatinine, Alanine aminotransferase (ALT), and total protein levels. All mentioned parameters were determined using an auto-analyzer (Rx Daytona, Randox, Antrim, UK).

**Statistical analysis.** At least three independent experiments were carried out and values were presented as the mean ± SEM or mean ± SD. All the statistical analysis was done using GraphPad Prism version 8 software. Unpaired, two-tailed, student`s *t*-test or ANOVA with Tukey post-hoc analysis was employed to test the statistical significance of the data. The number of biological replicates denoted by (N) and number of entities quantified denoted by (n), are mentioned in the respective Figure legends. For the animal toxicity experiment, four animals were used per group and the statistical significance was determined using the Student's t-test. A value of $P < 0.05$ was considered significant throughout the paper.

### Supplementary Figures

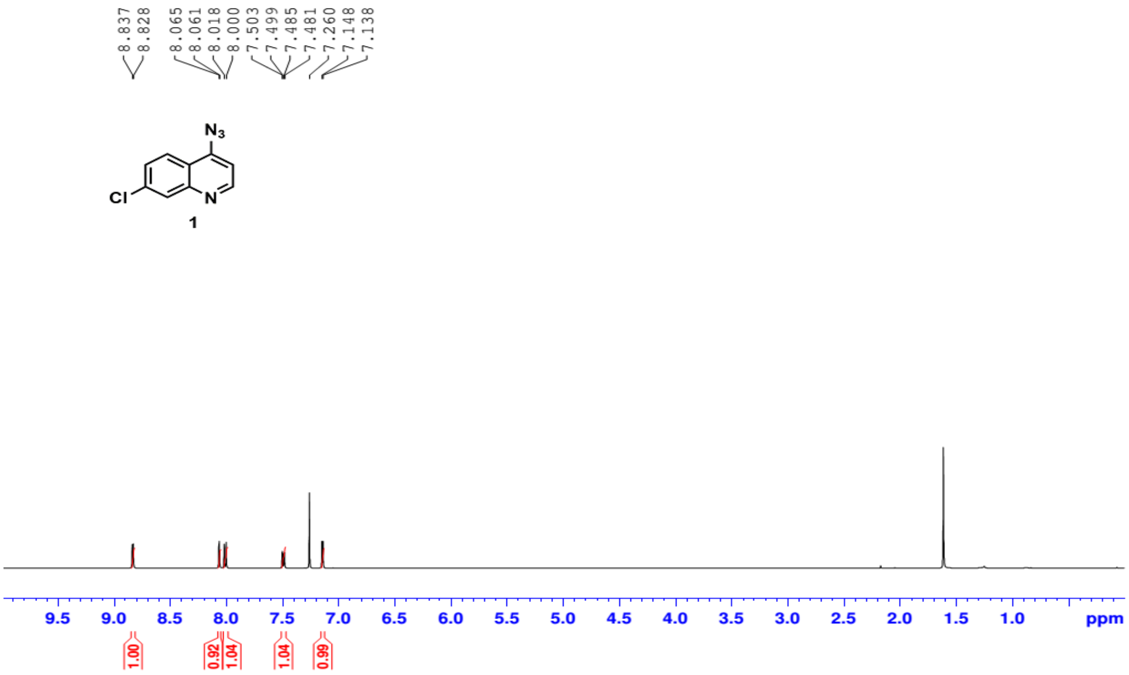


**Fig. S1: $^1$H NMR spectrum of compound 1**

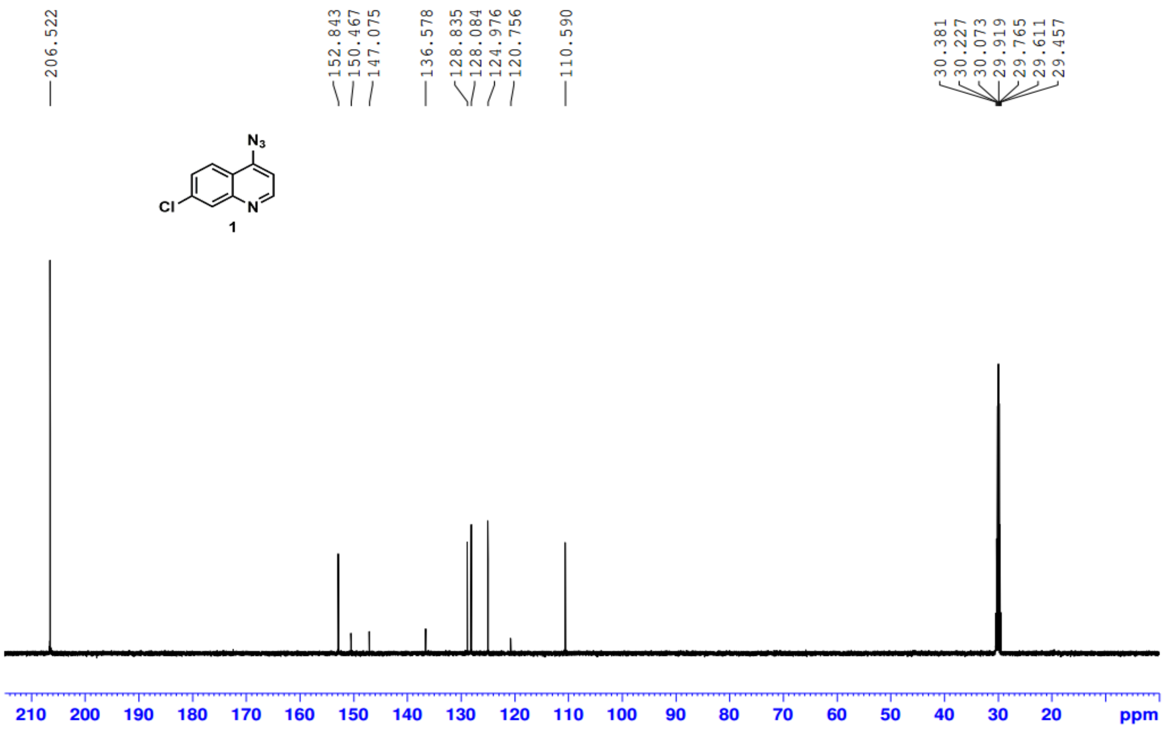


**Fig. S2: $^{13}$C NMR spectrum of compound 1**

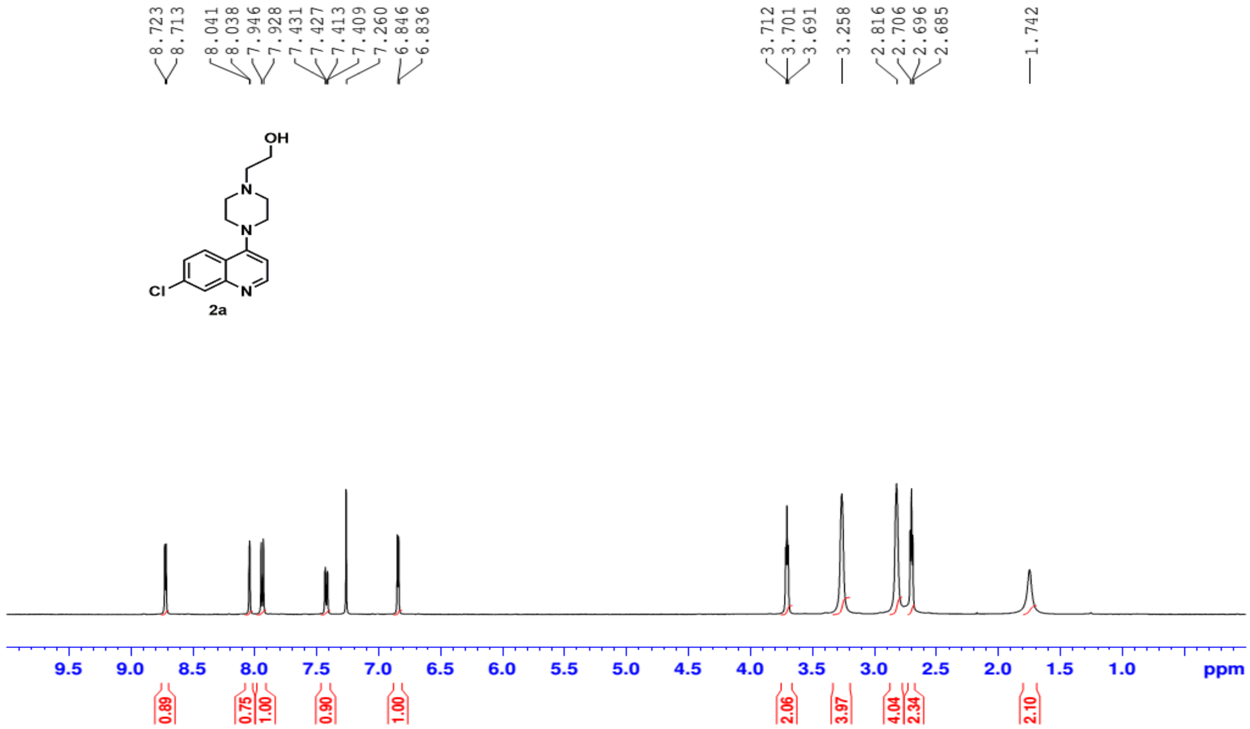

**Fig. S3: $^{1}$H NMR spectrum of compound 2a**

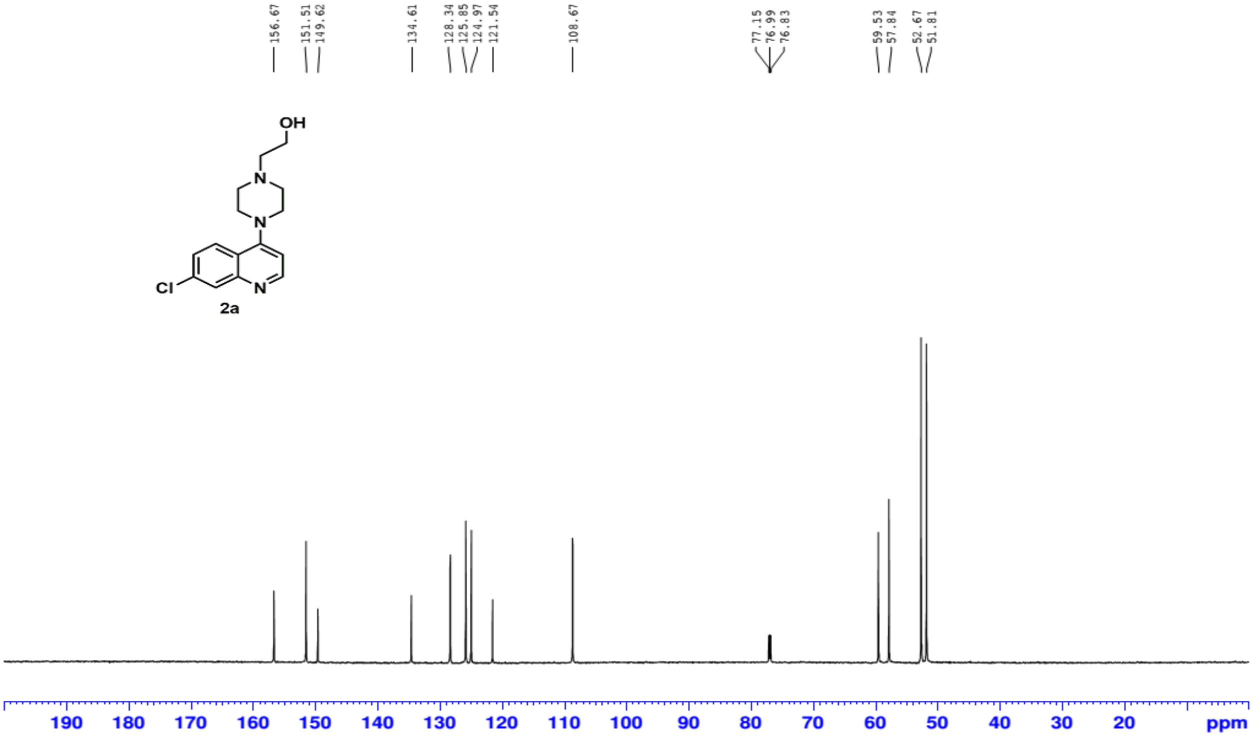

**Fig. S4: $^{13}$C NMR spectrum of compound 2a**

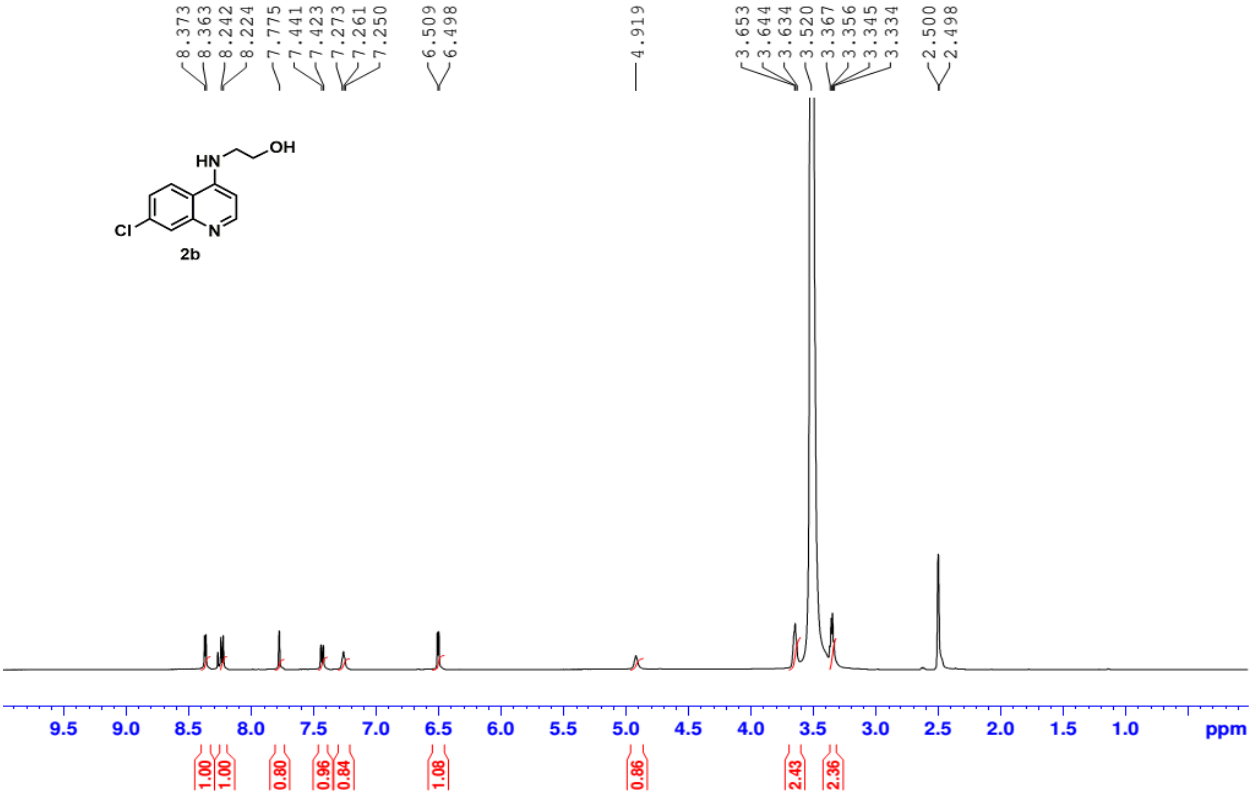


**Fig. S5: $^{1}$H NMR spectrum of compound 2b**

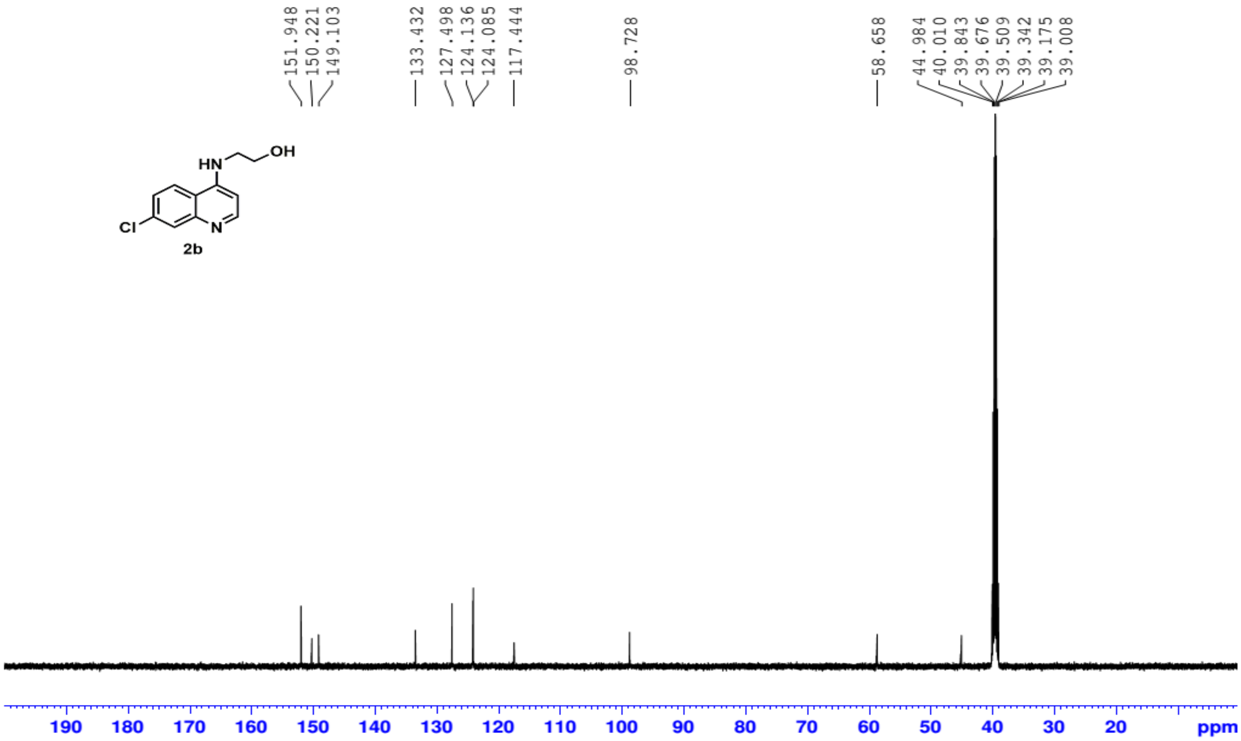


**Fig. S6: $^{13}$C NMR spectrum of compound 2b**

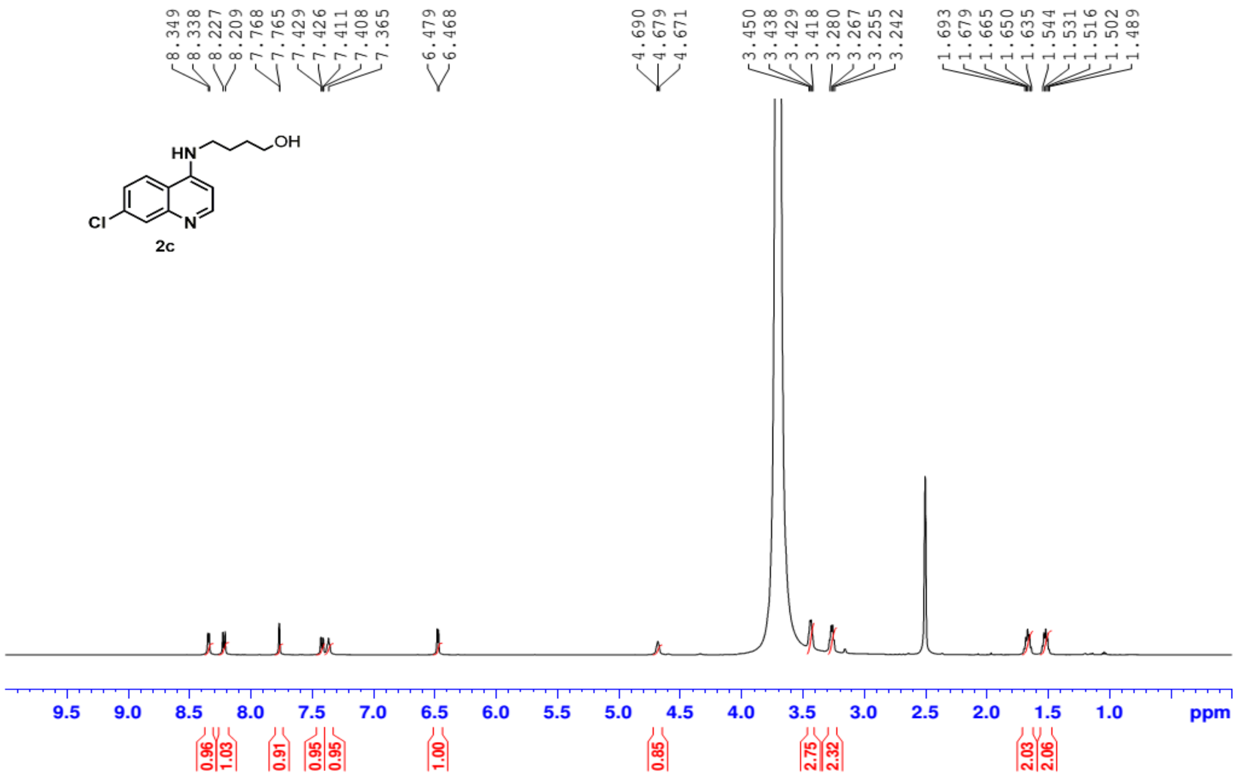


**Fig. S7: $^{1}$H NMR spectrum of compound 2c**

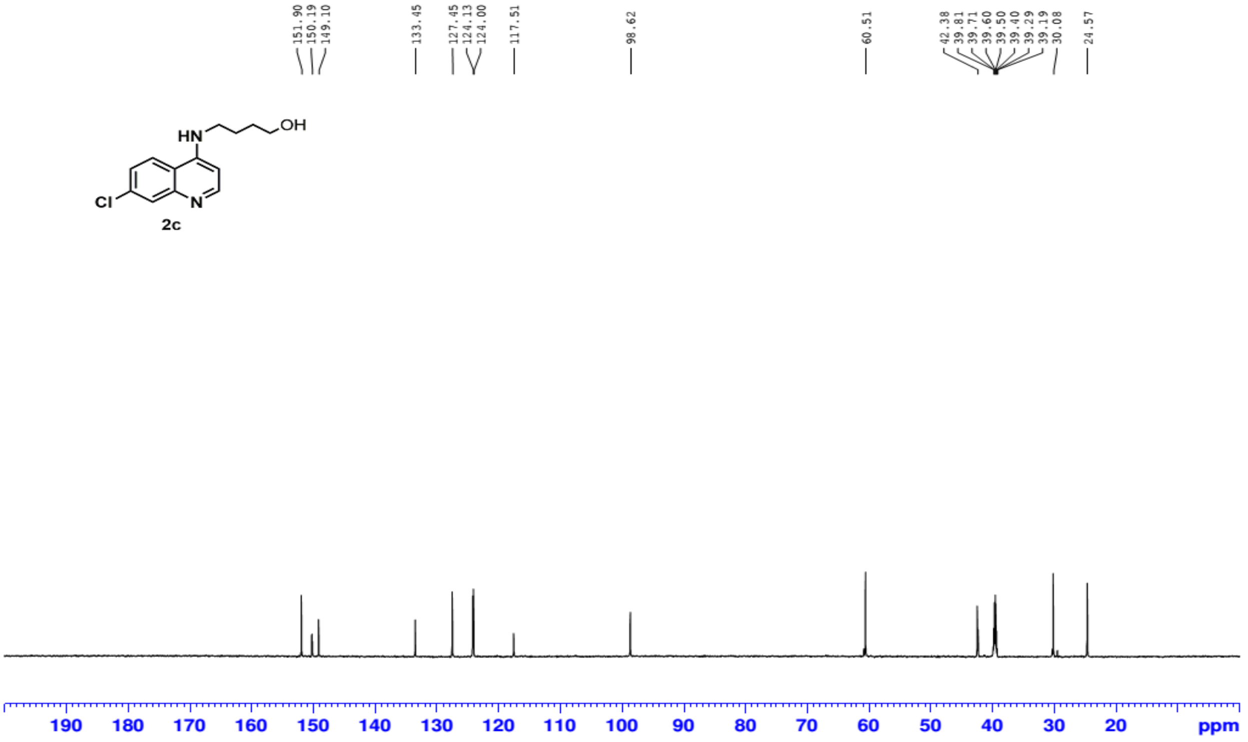


**Fig. S8: $^{13}$C NMR spectrum of compound 2c**

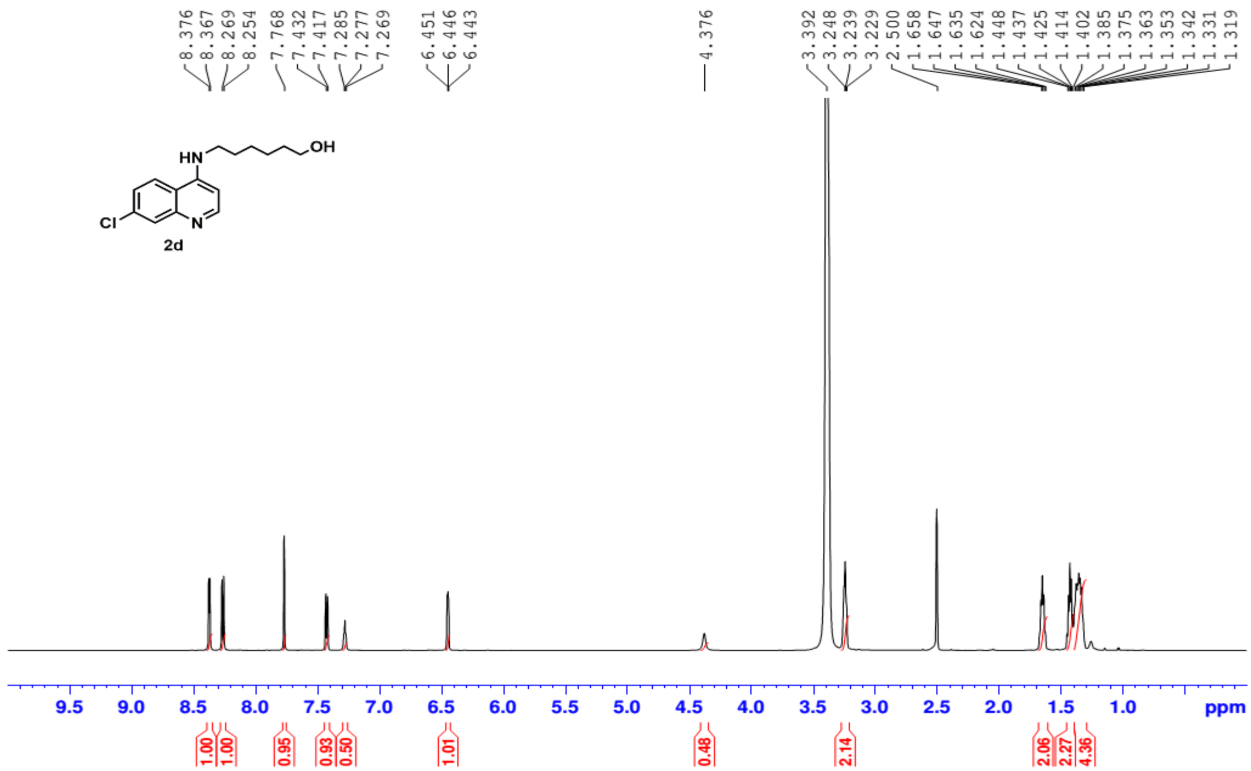


**Fig. S9: $^{1}$H NMR spectrum of compound 2d**

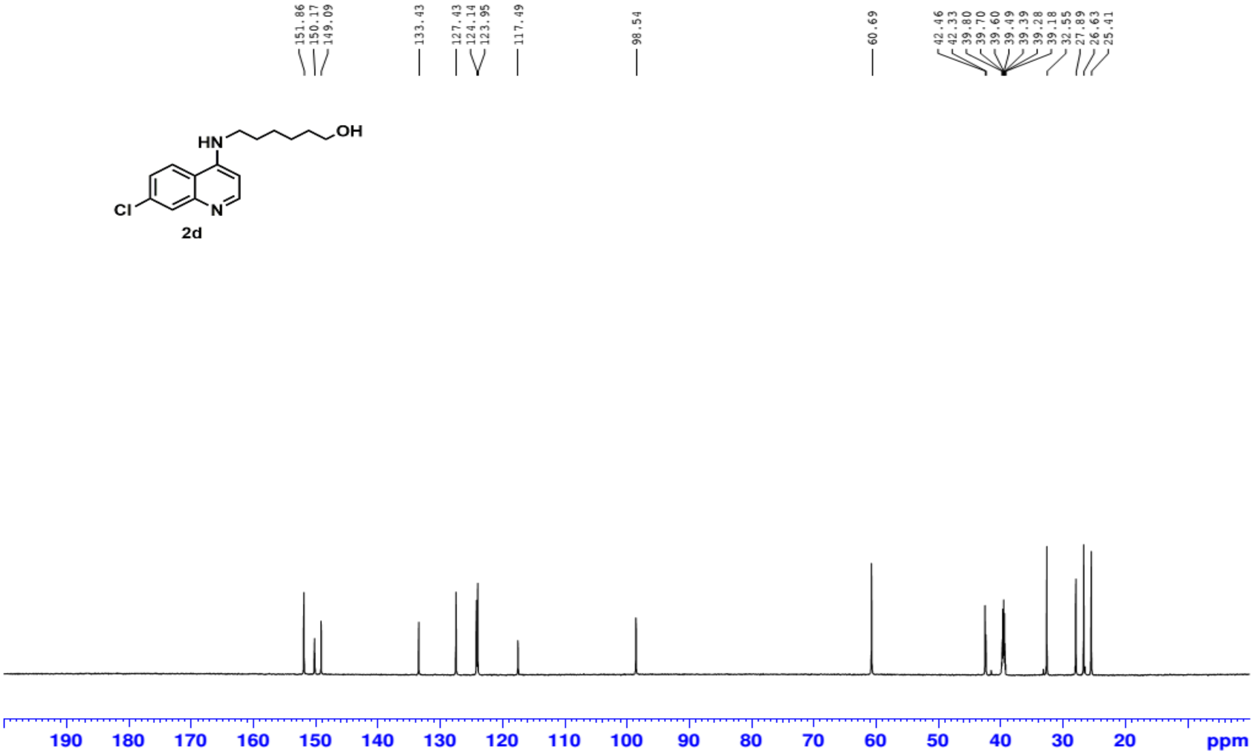


**Fig. S10: $^{13}$C NMR spectrum of compound 2d**

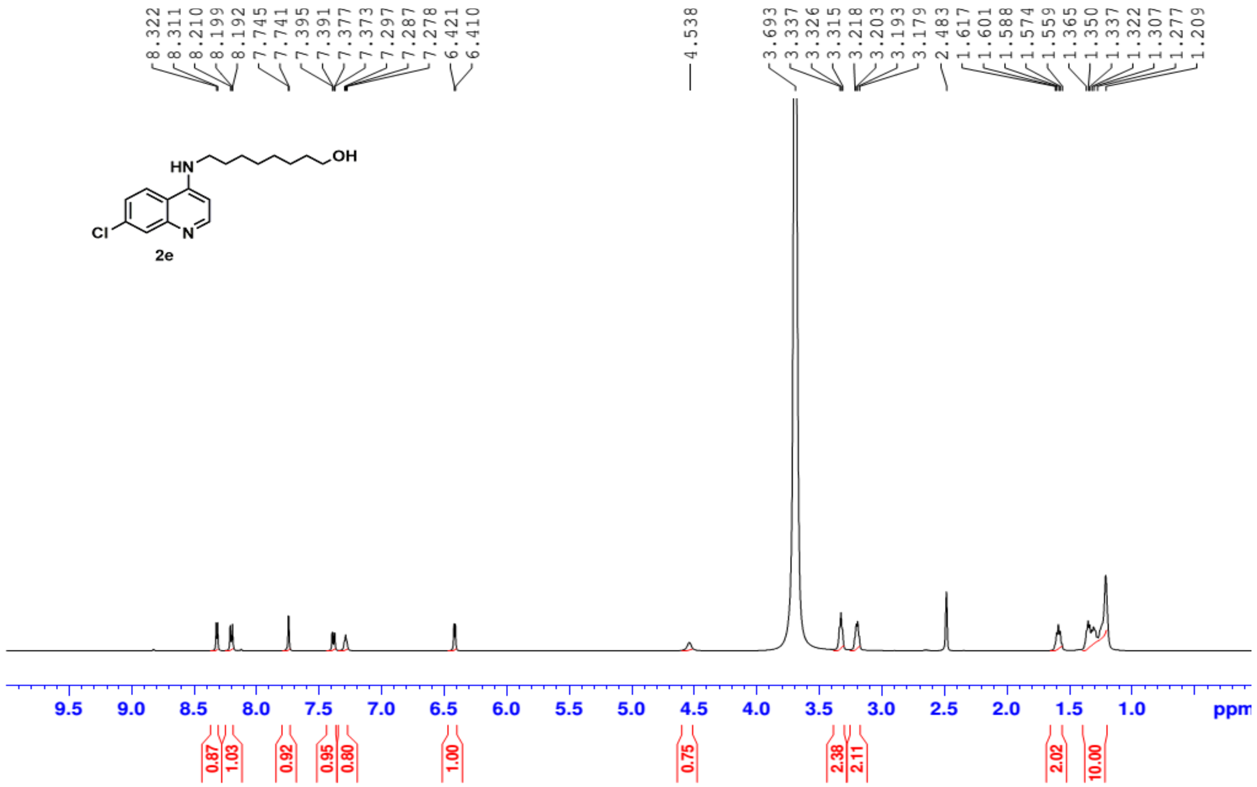


**Fig. S11: $^{1}$H NMR spectrum of compound 2e**

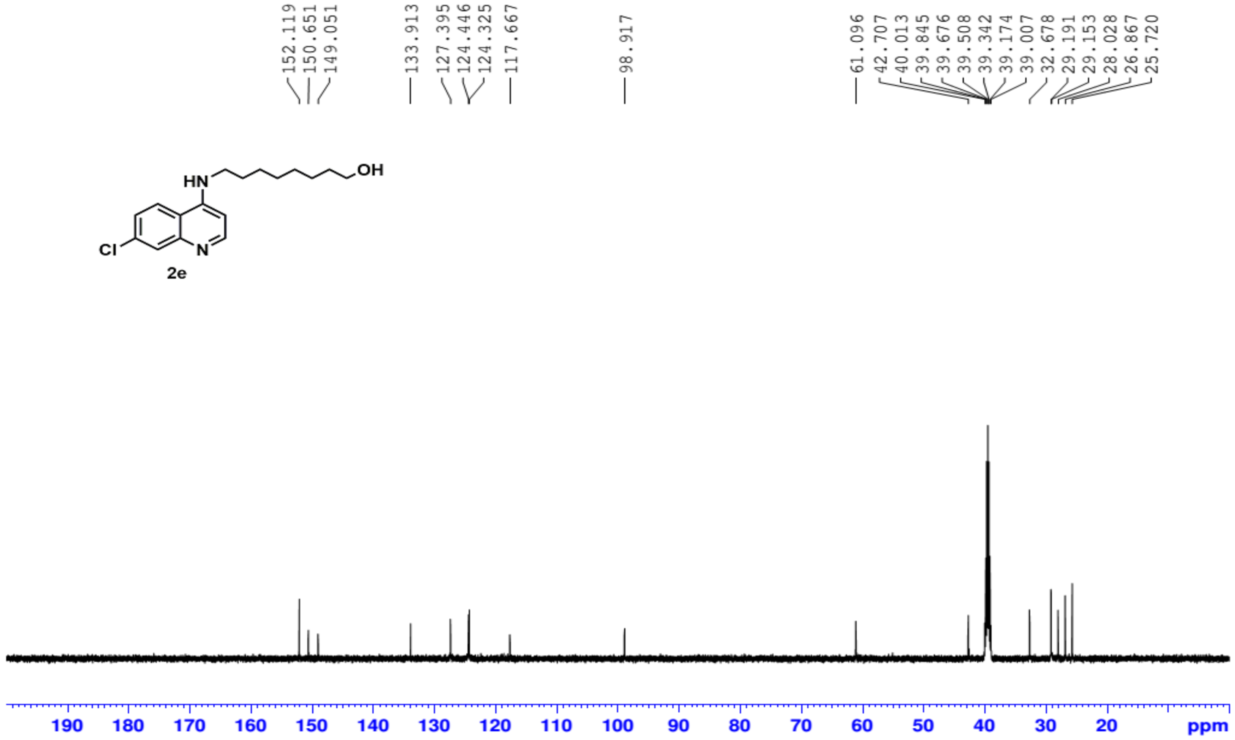


**Fig. S12: $^{13}$C NMR spectrum of compound 2e**

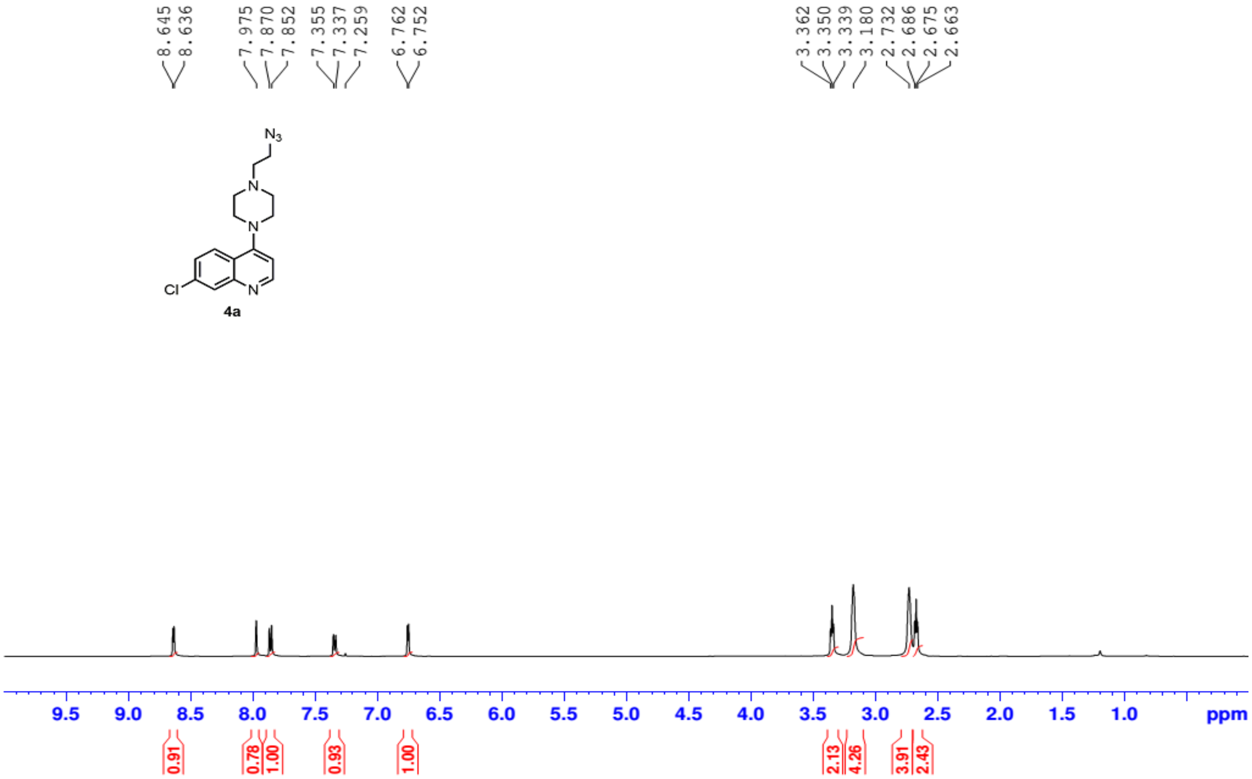


Fig. S13: $^{1}$H NMR spectrum of compound 4a

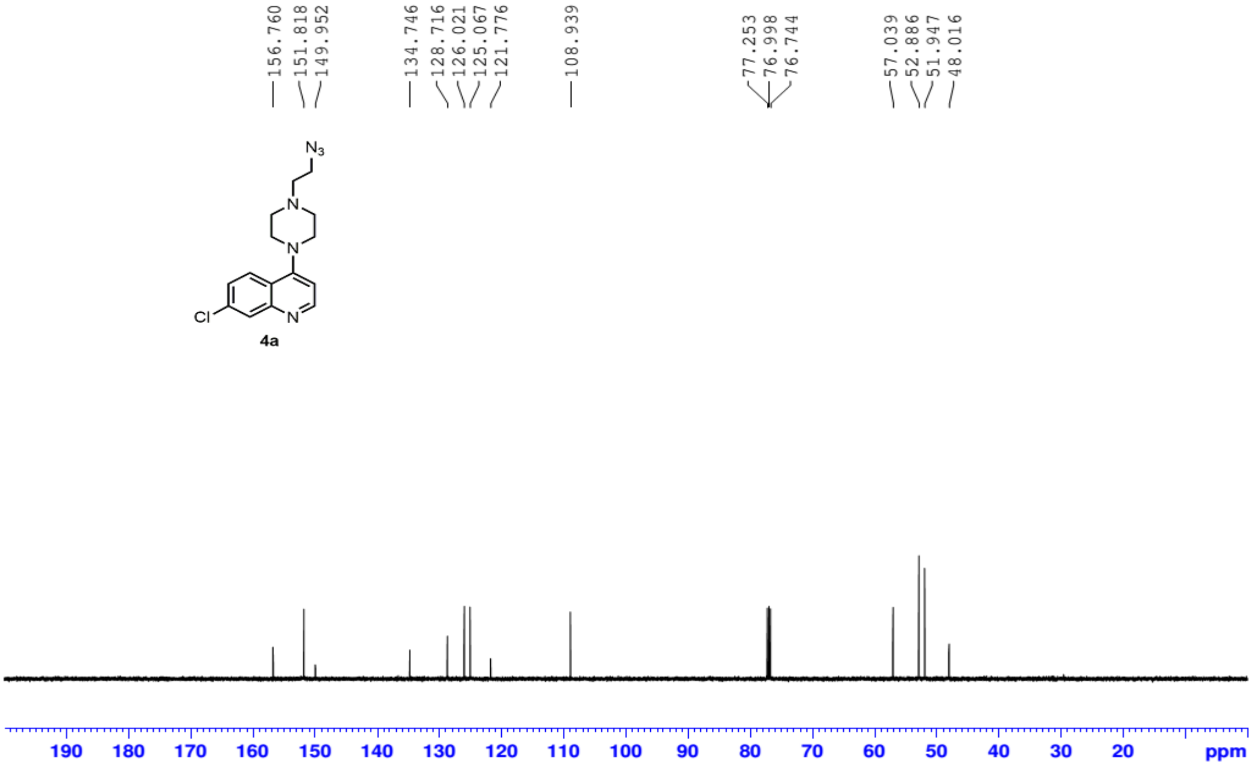


Fig. S14: $^{13}$C NMR spectrum of compound 4a

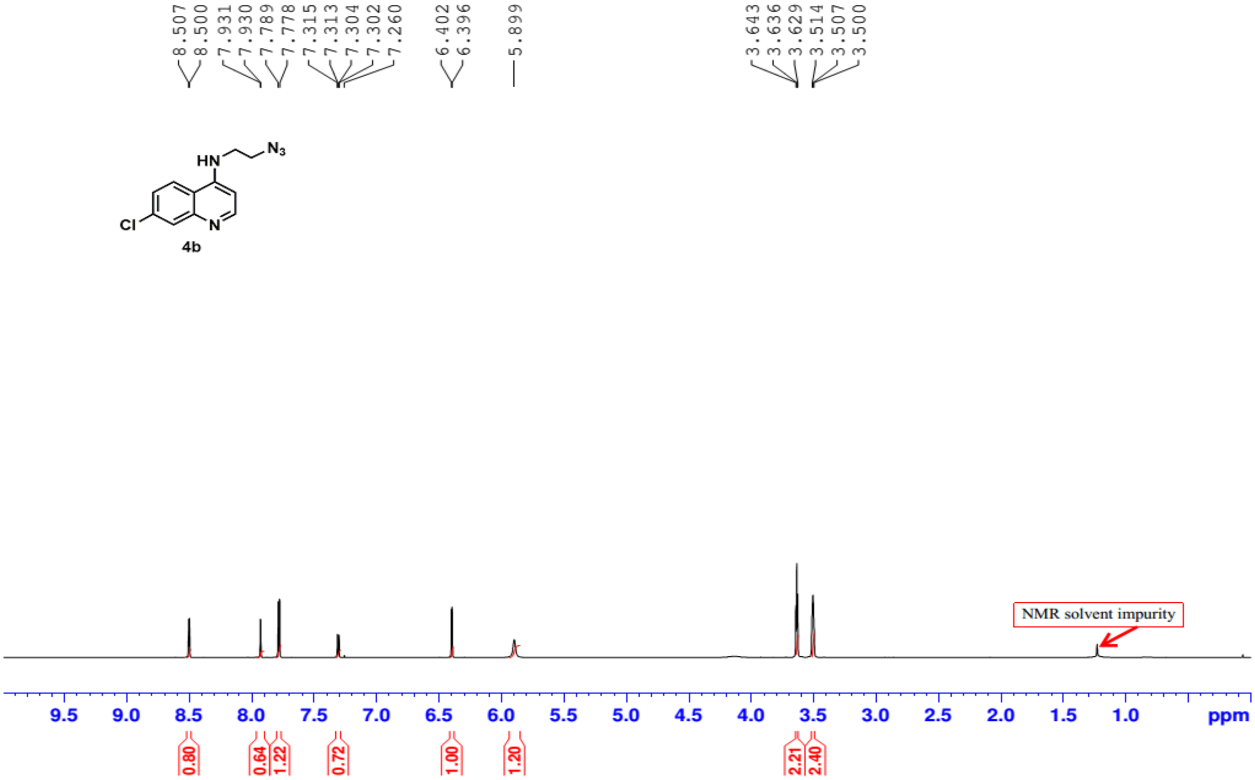


**Fig. S15: $^{1}$H NMR spectrum of compound 4b**

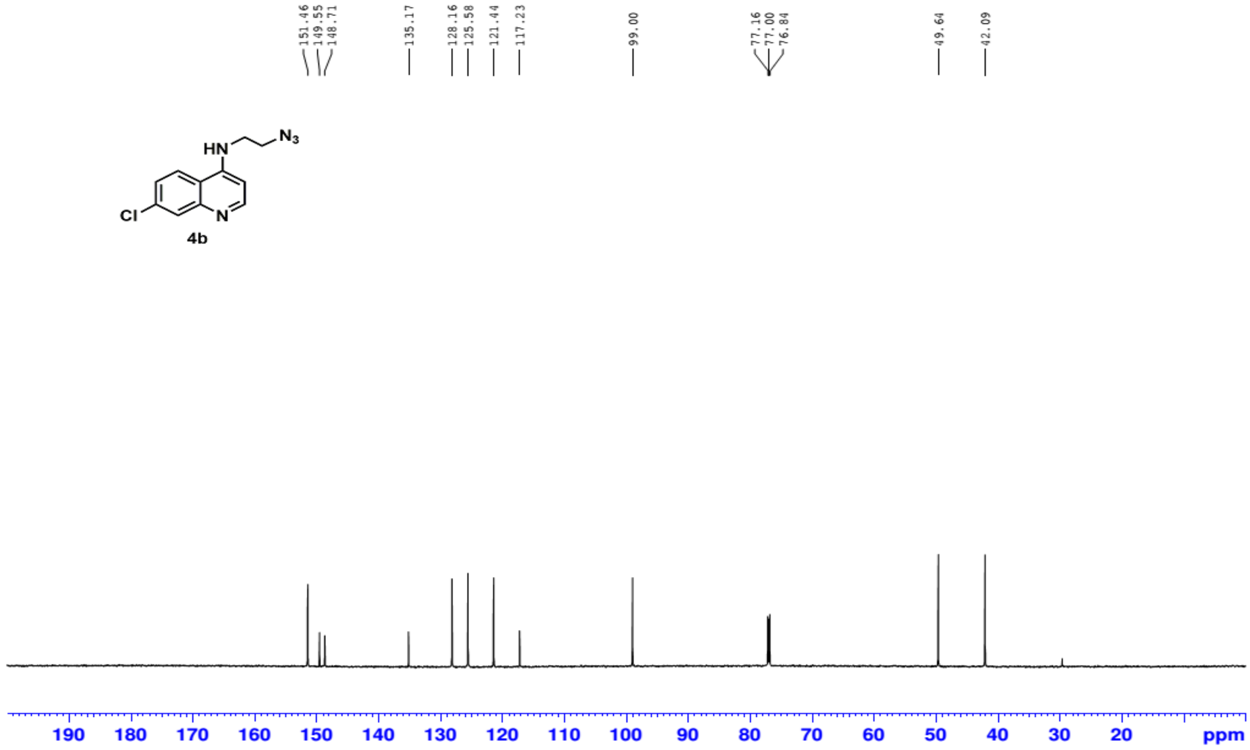


**Fig. S16: $^{13}$C NMR spectrum of compound 4b**

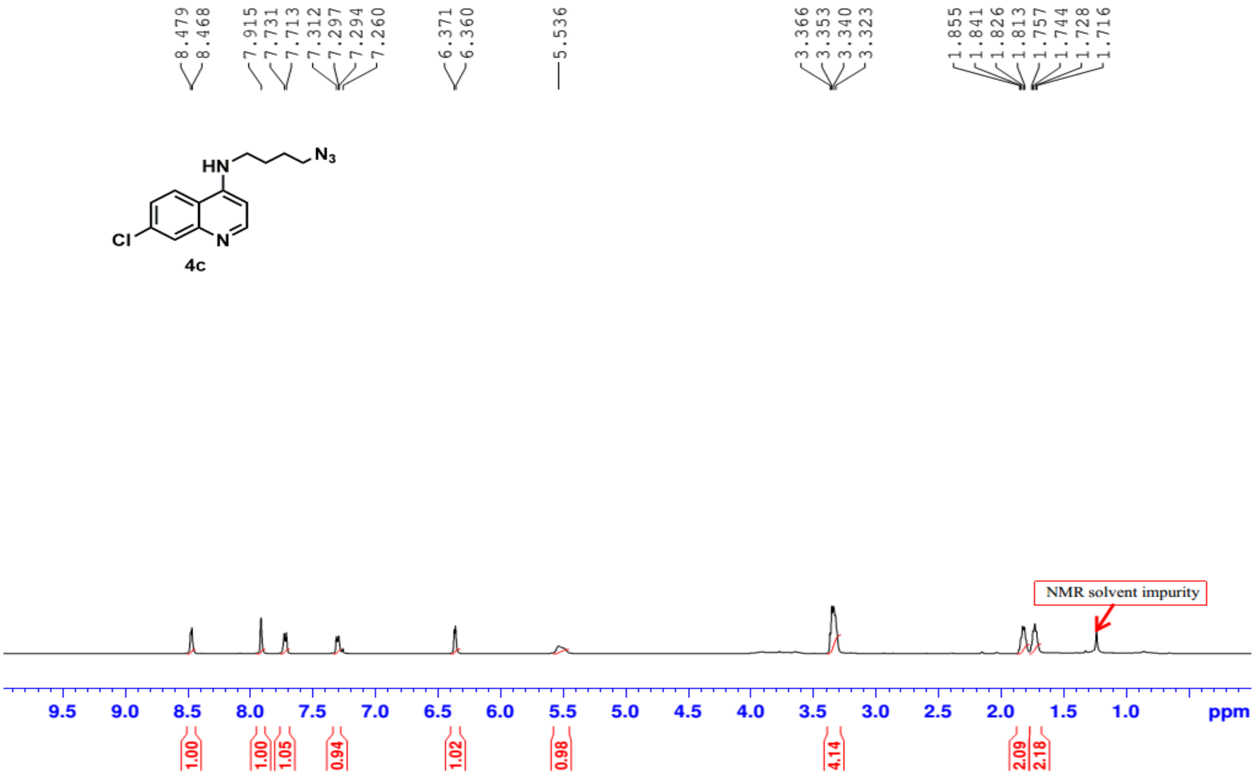


**Fig. S17: $^{1}$H NMR spectrum of compound 4c**

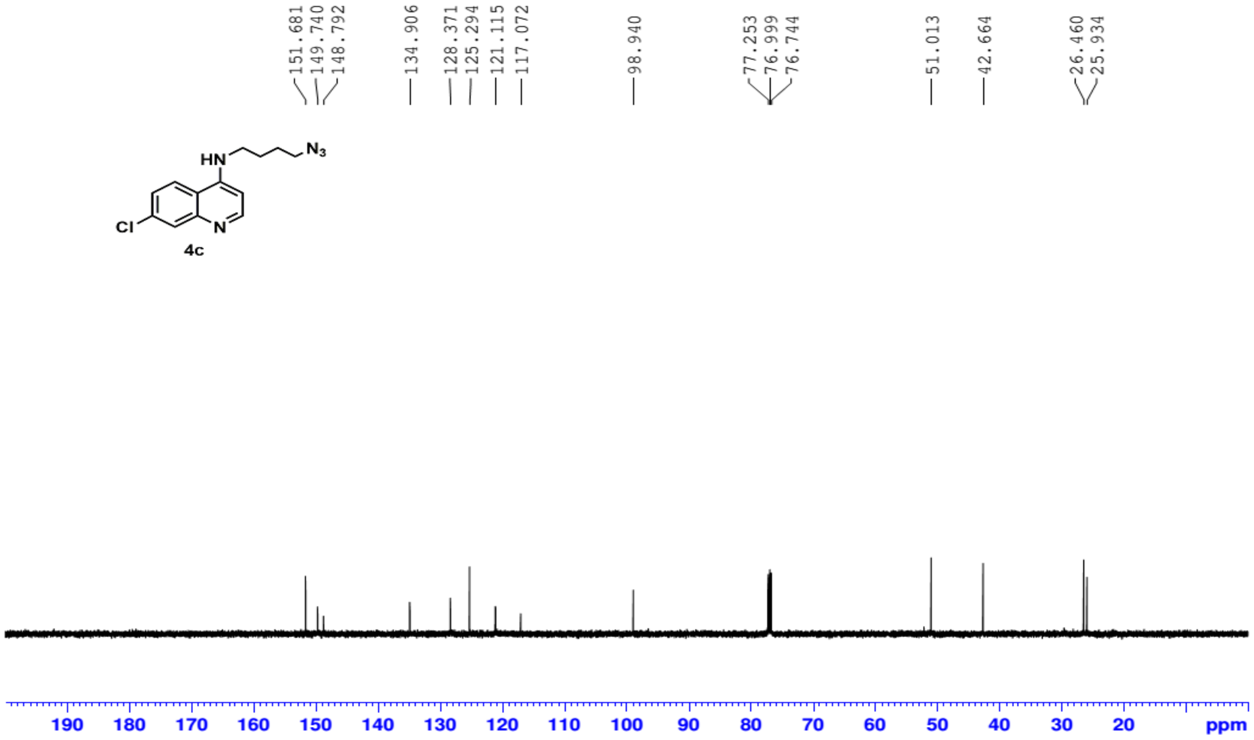


**Fig. S18: $^{13}$C NMR spectrum of compound 4c**

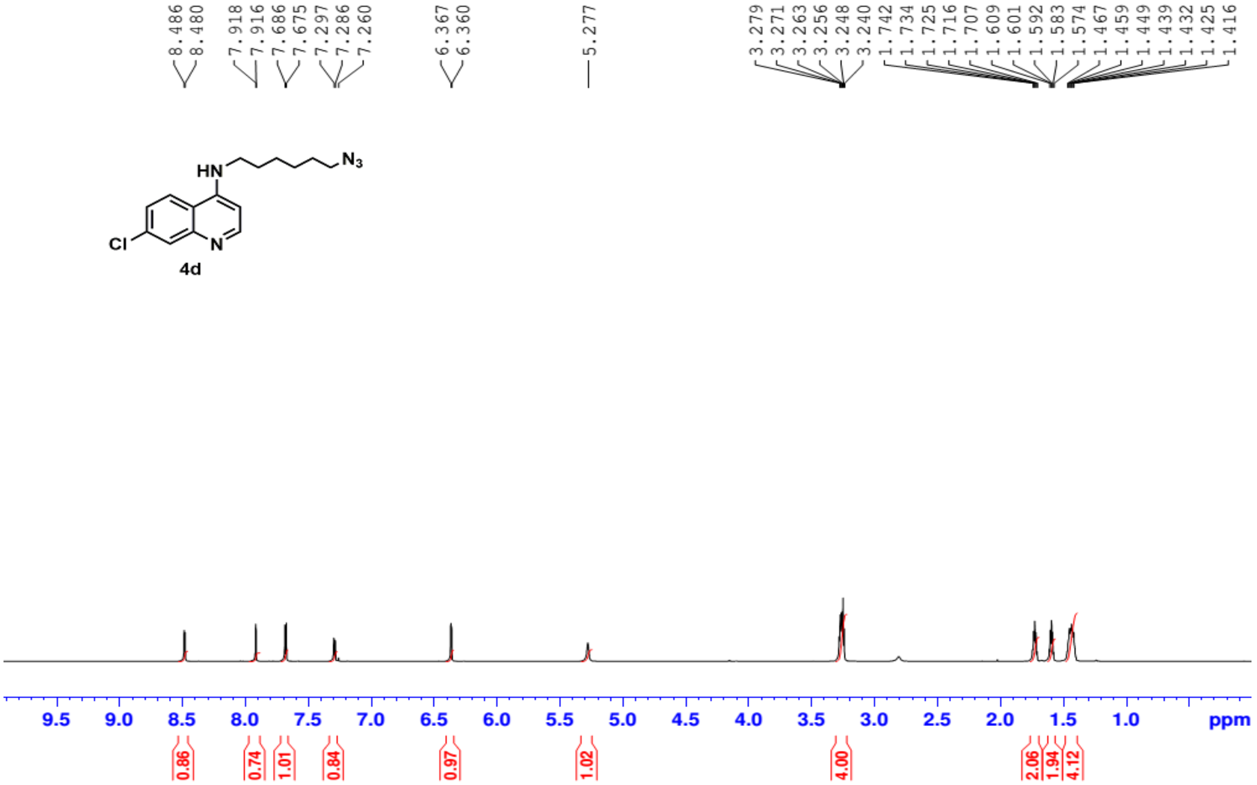


**Fig. S19: $^{1}$H NMR spectrum of compound 4d**

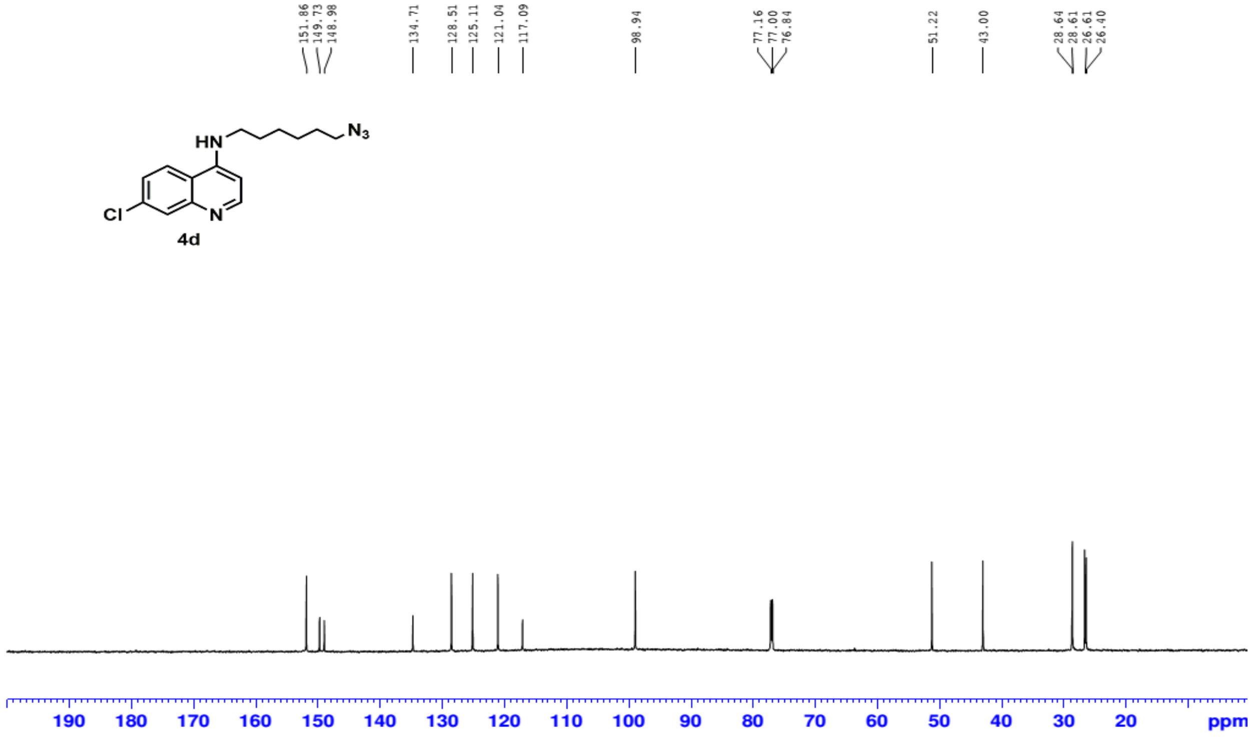


**Fig. S20: $^{13}$C NMR spectrum of compound 4d**

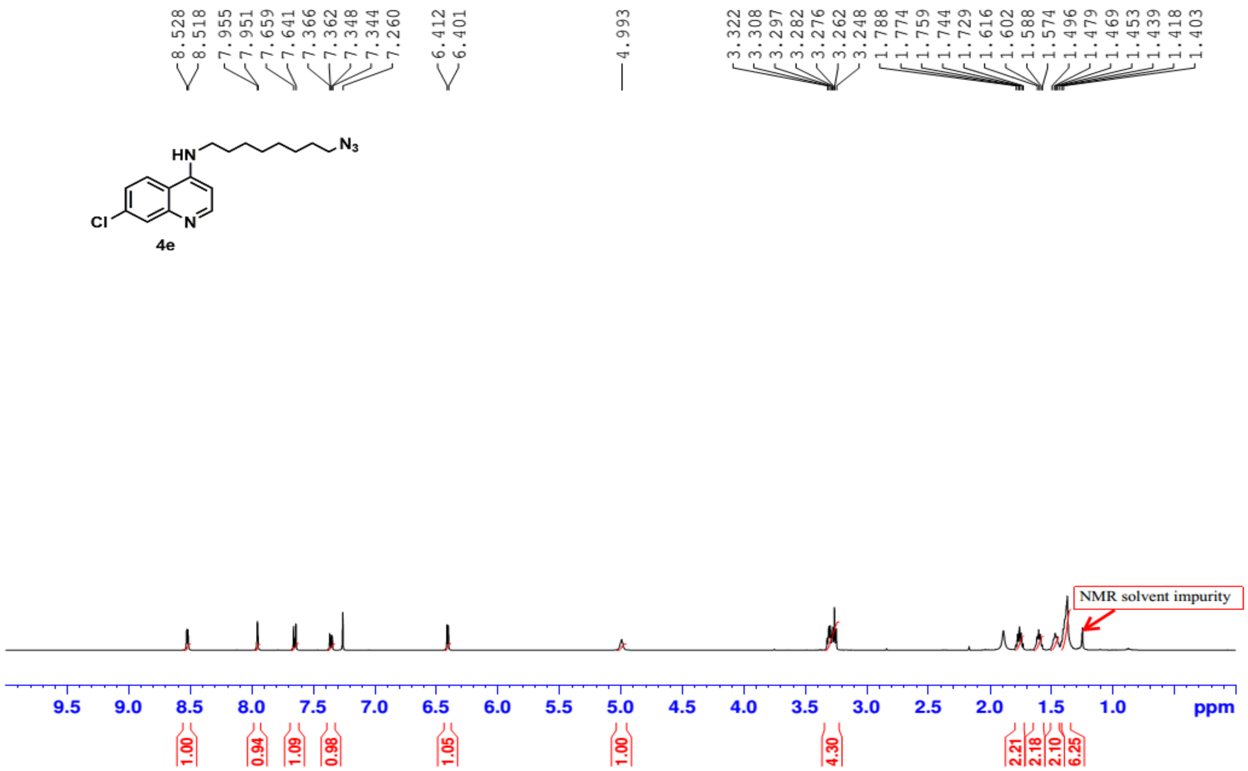


**Fig. S21: $^{1}$H NMR spectrum of compound 4e**

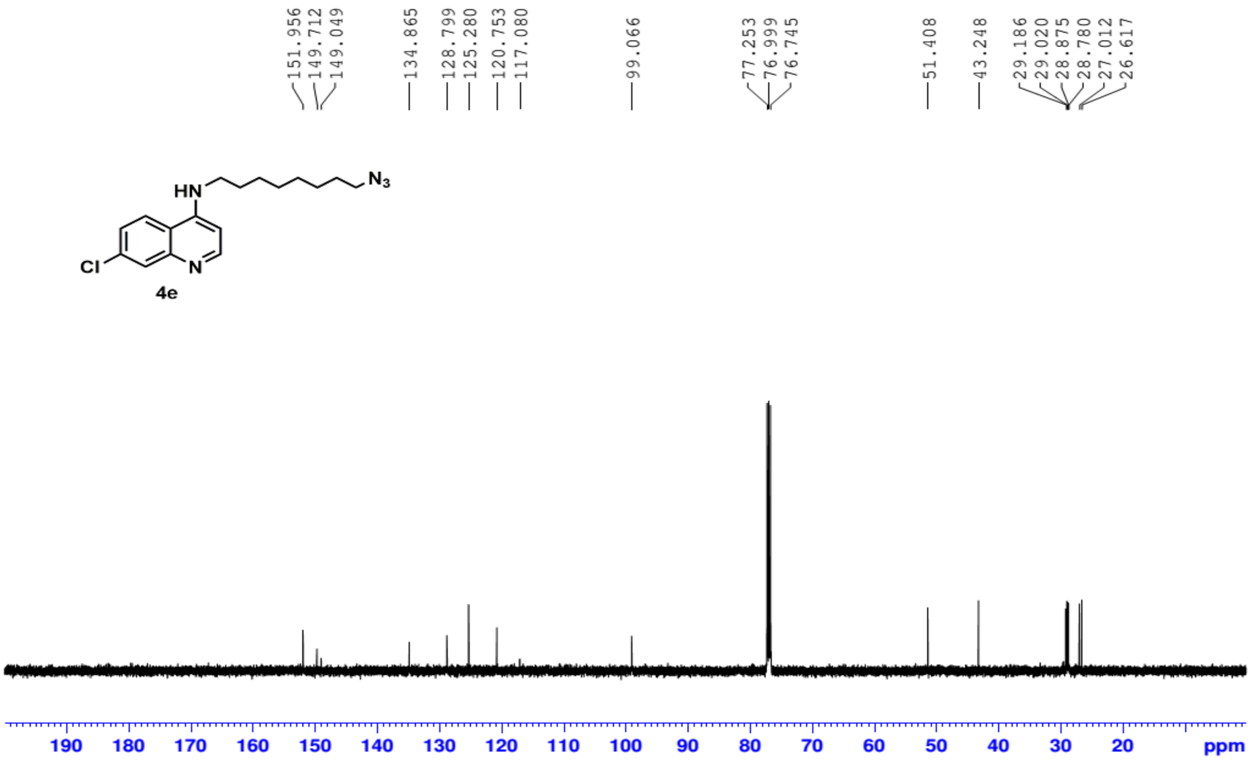


**Fig. S22: $^{13}$C NMR spectrum of compound 4e**

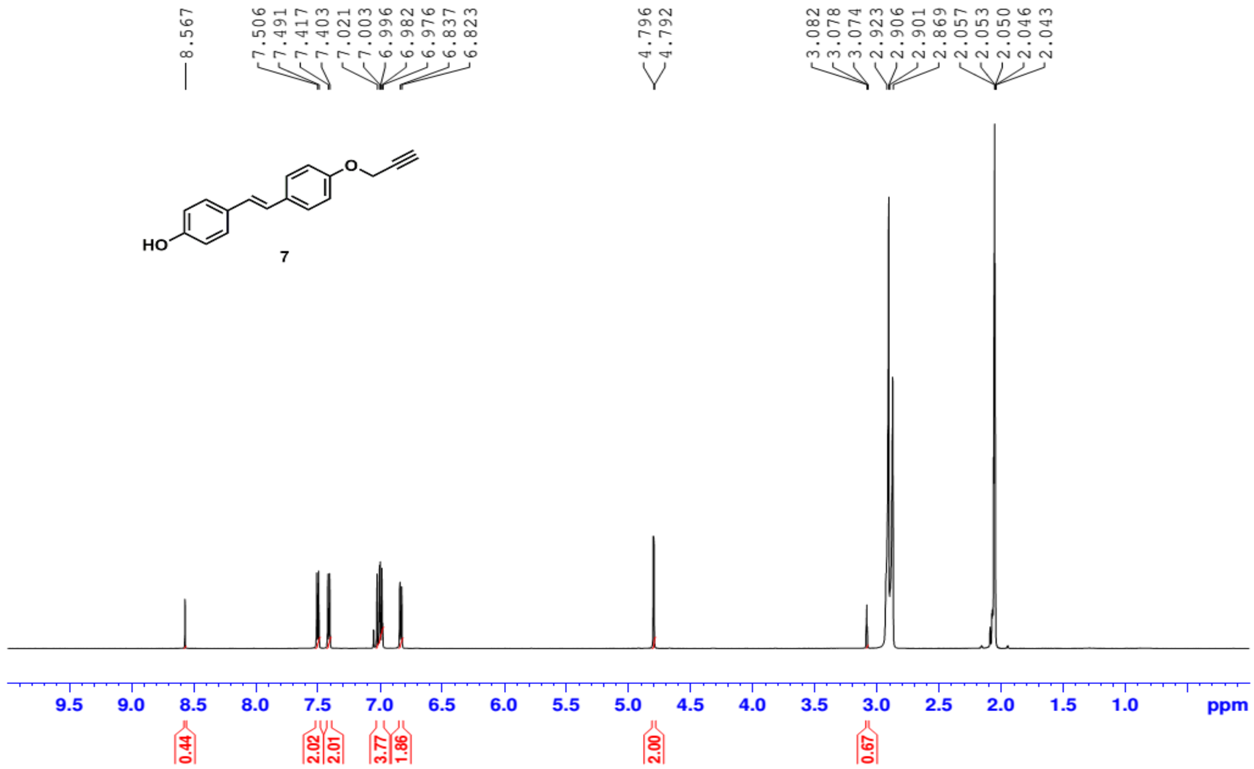


**Fig. S23: [1]H NMR spectrum of compound 7**

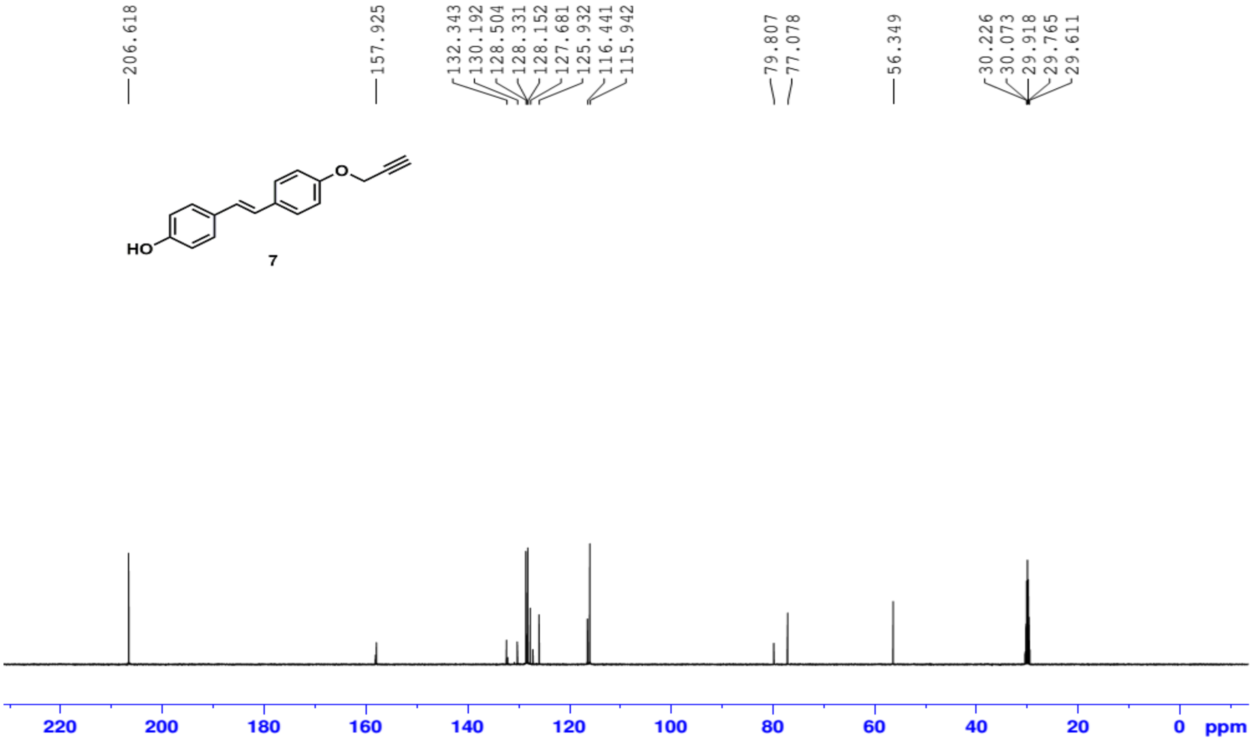


**Fig. S24: [13]C NMR spectrum of compound 7**

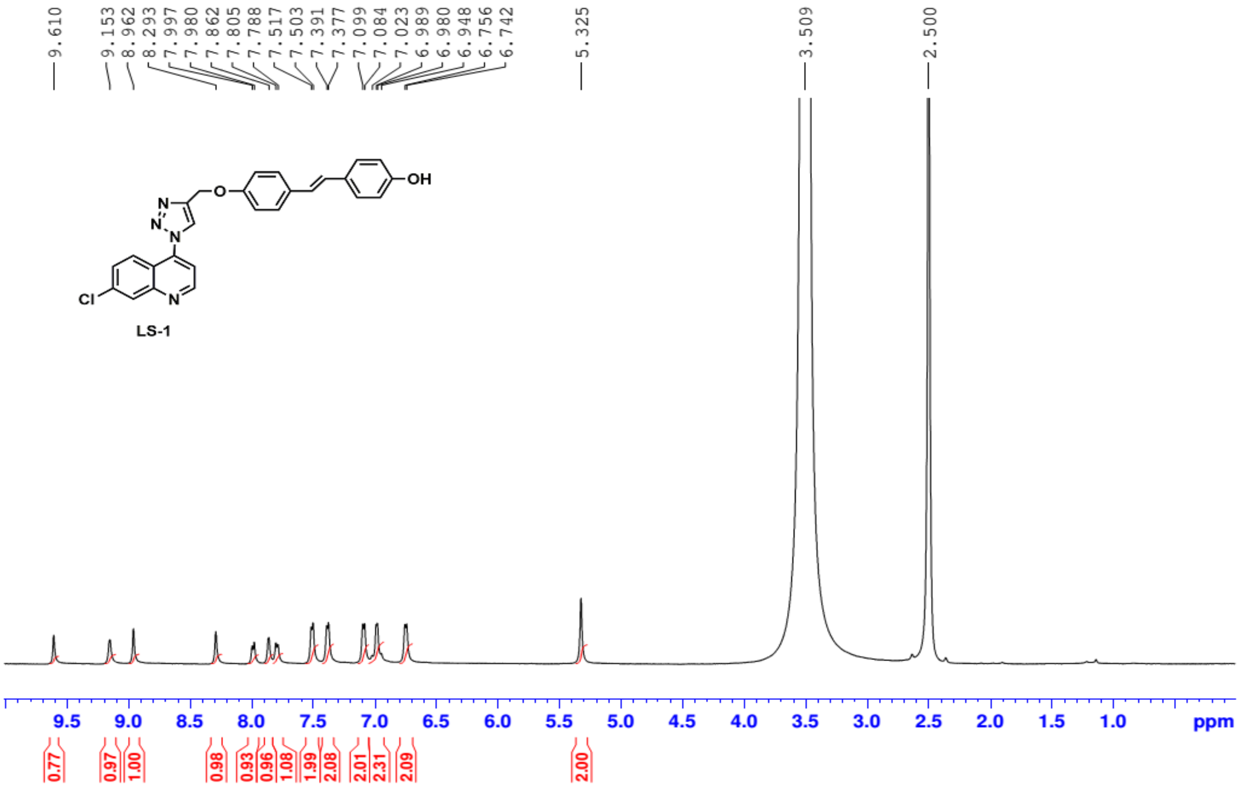


**Fig. S25: $^{1}$H NMR spectrum of compound LS1**

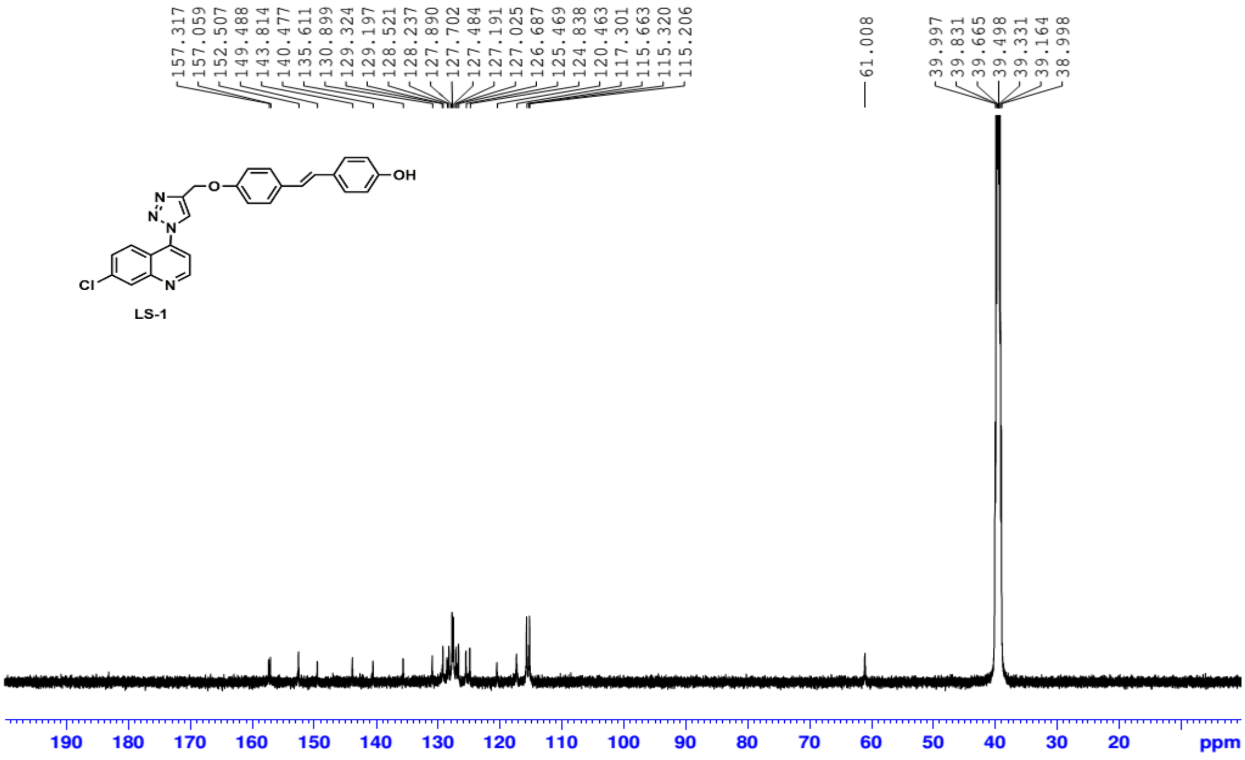


**Fig. S26: $^{13}$C NMR spectrum of compound LS1**

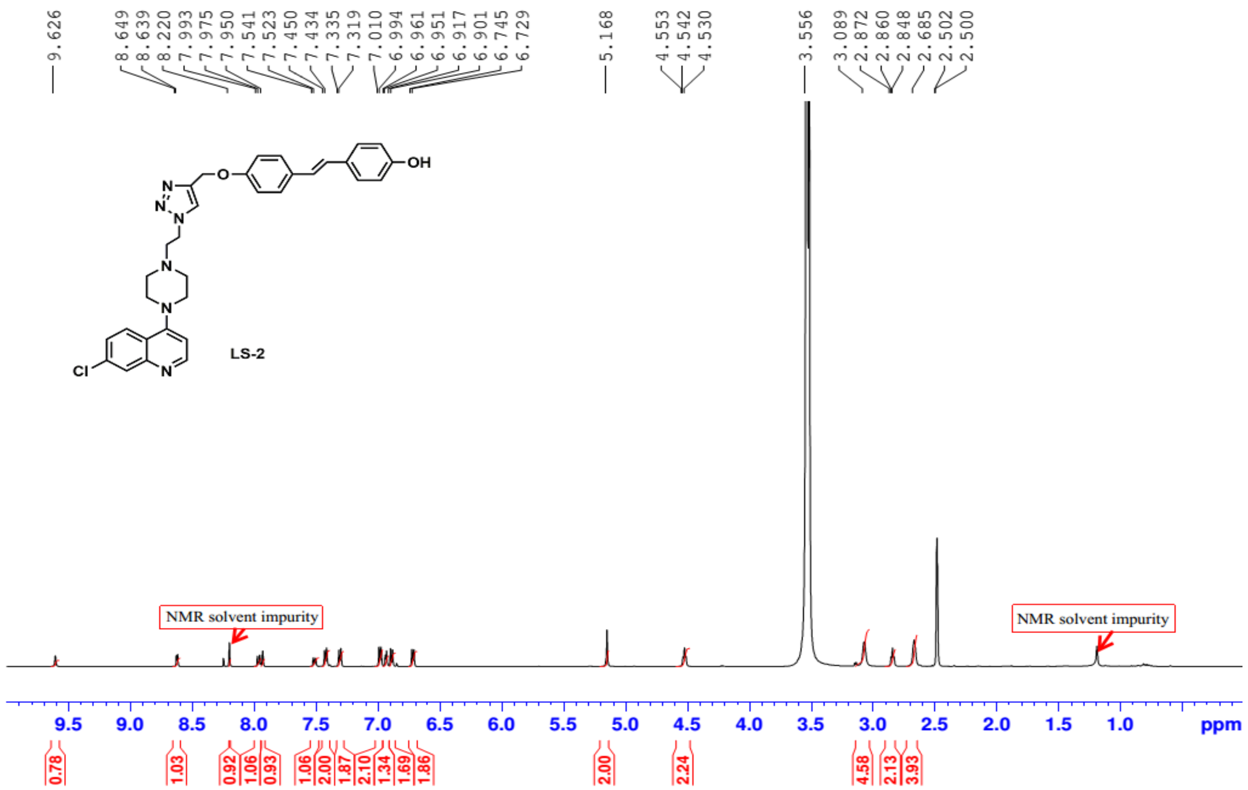


Fig. S27: $^{1}$H NMR spectrum of compound LS2

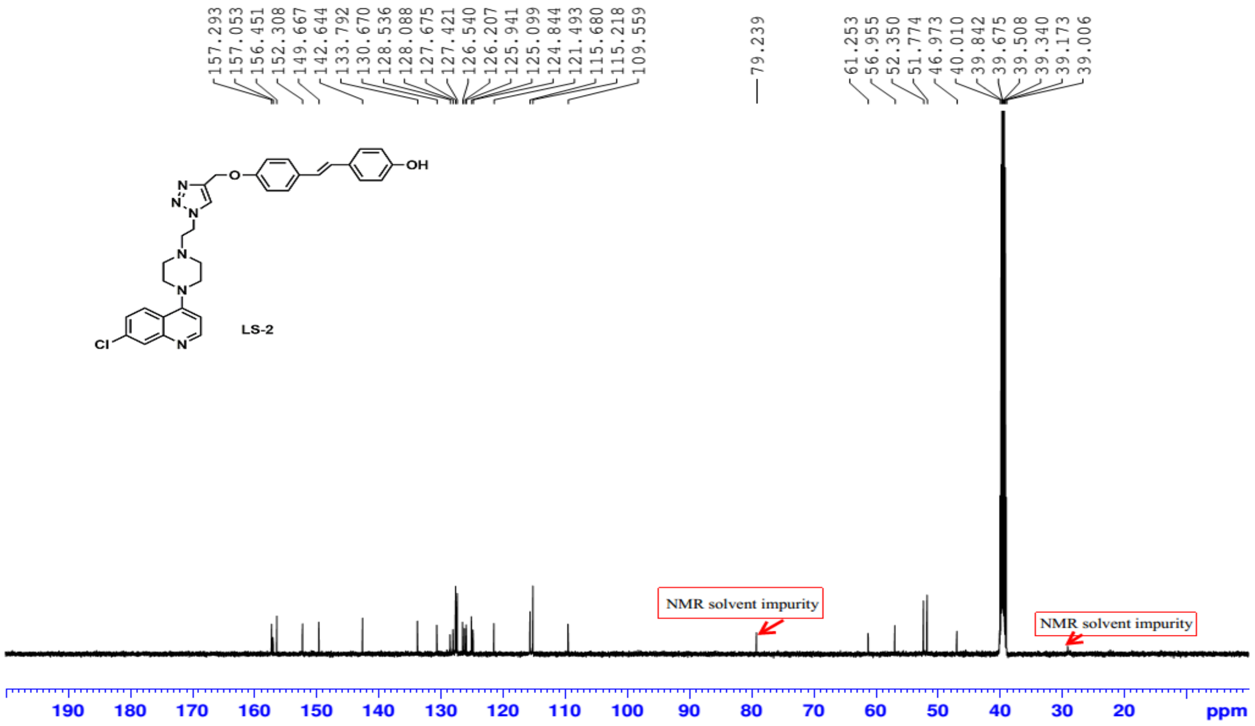


Fig. S28: $^{13}$C NMR spectrum of compound LS2

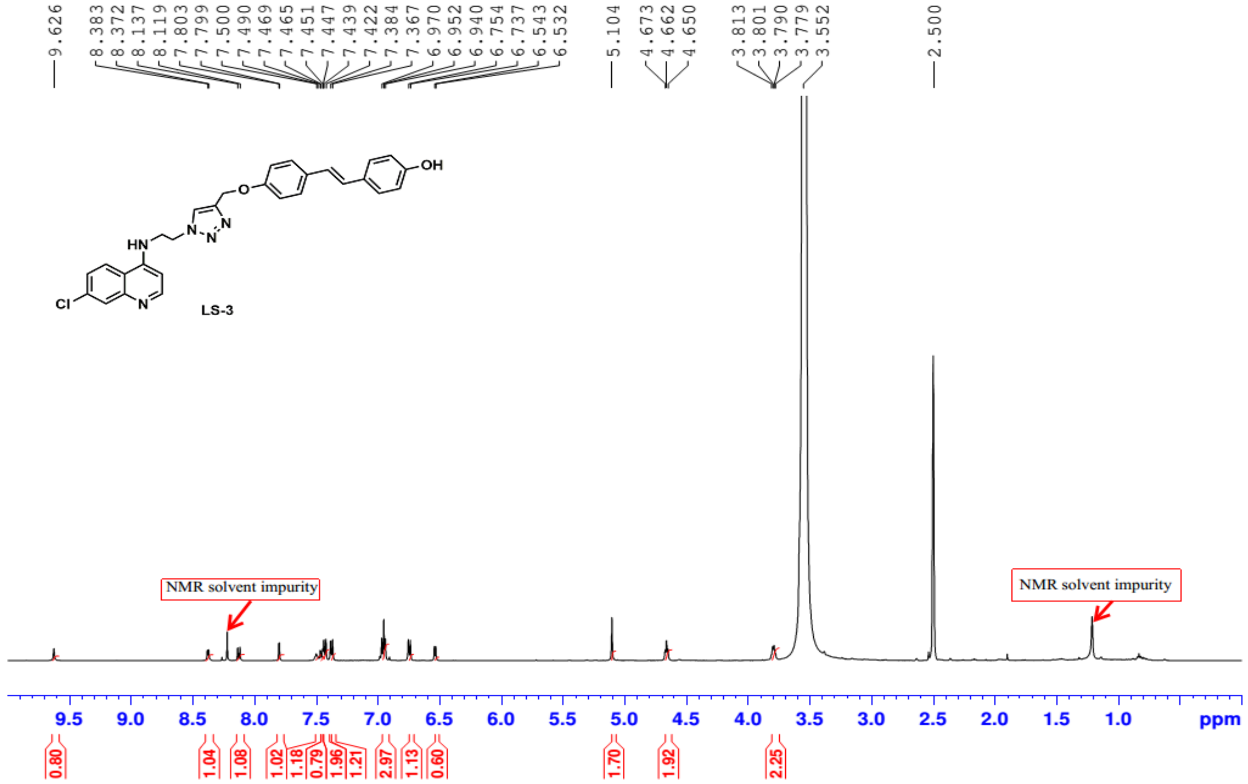


**Fig. S29: $^{1}$H NMR spectrum of compound LS3**

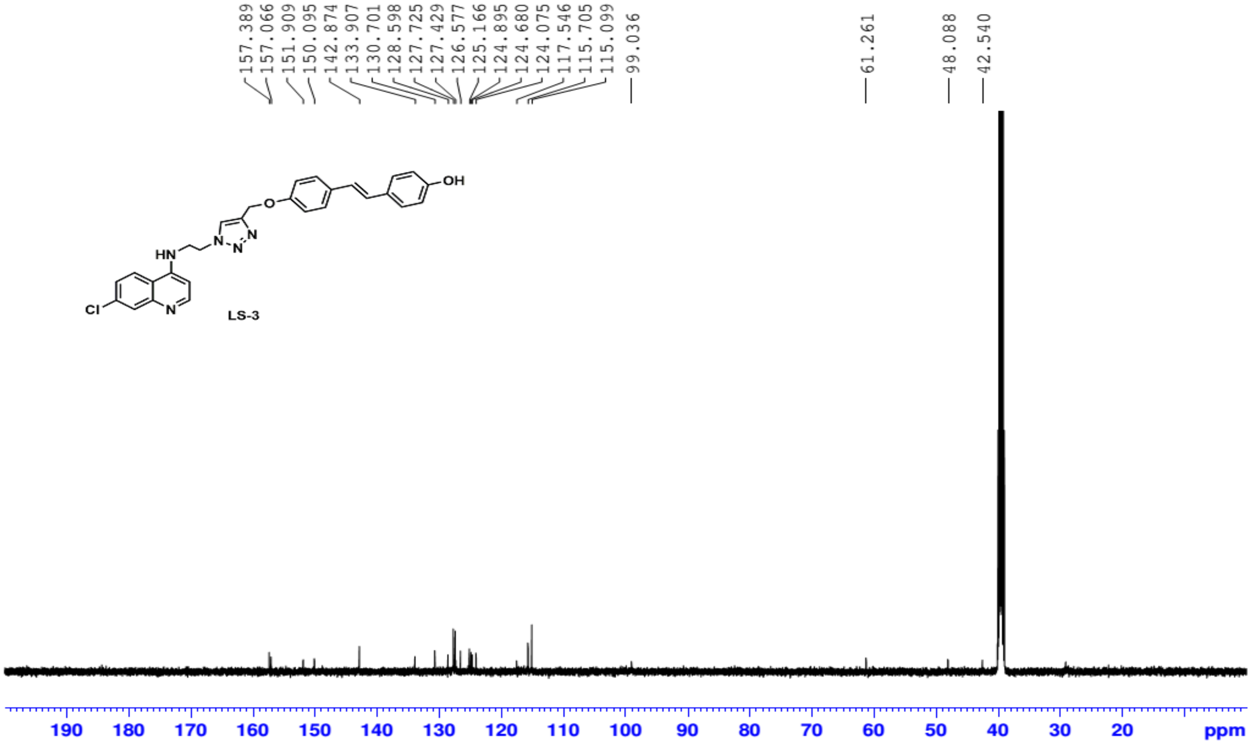


**Fig. S30: $^{13}$C NMR spectrum of compound LS3**

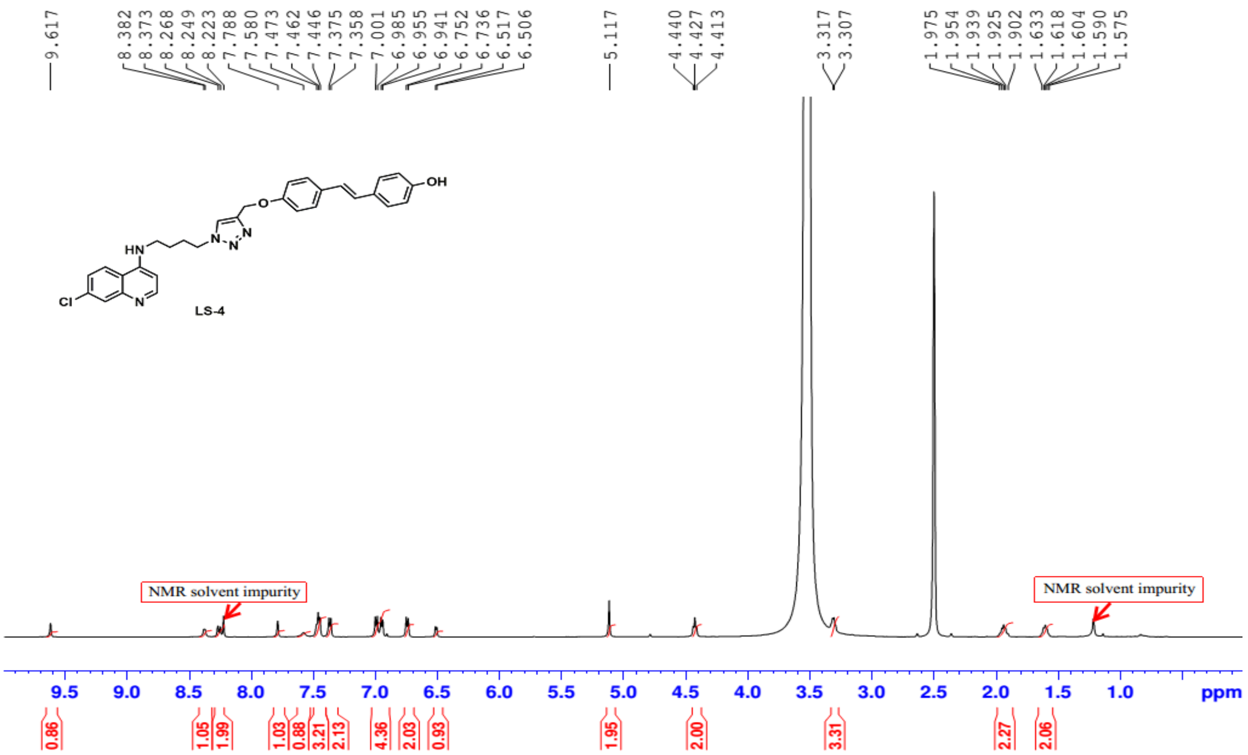


**Fig. S31: [1]H NMR spectrum of compound LS4**

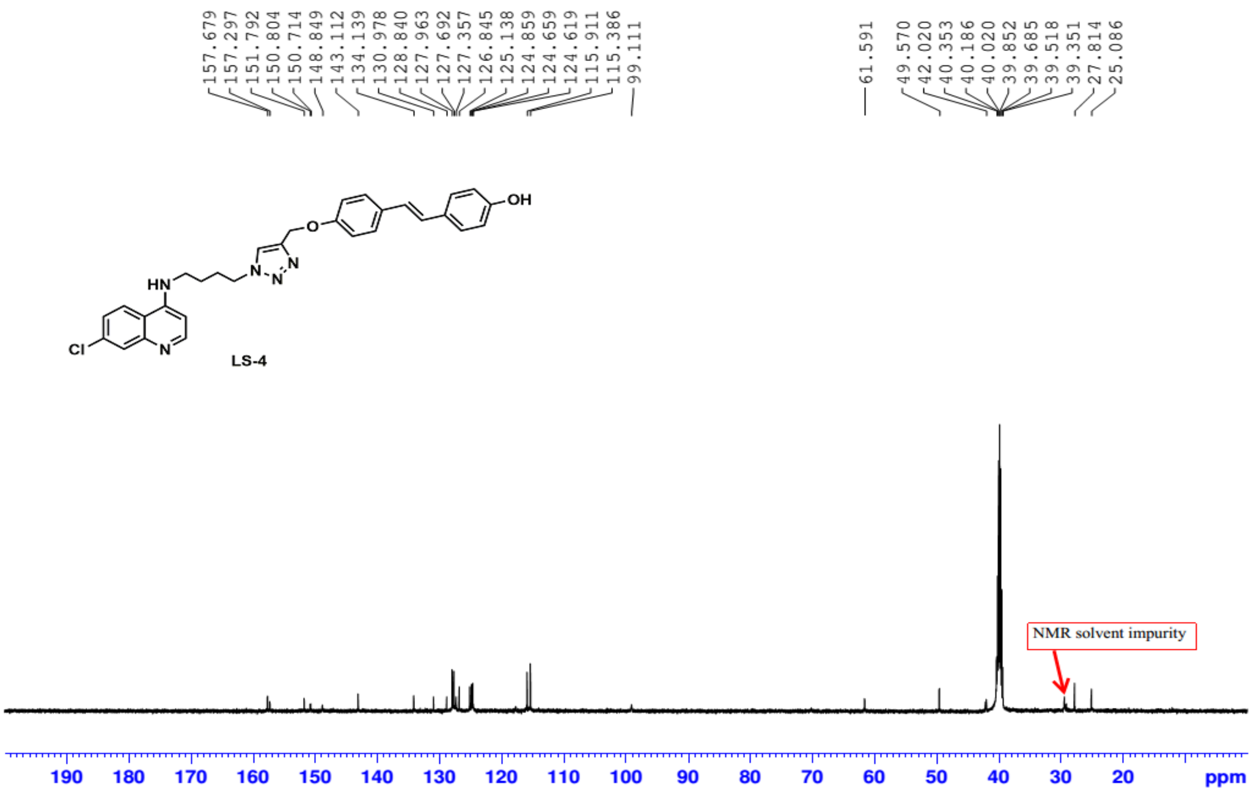


**Fig. S32: [13]C NMR spectrum of compound LS4**

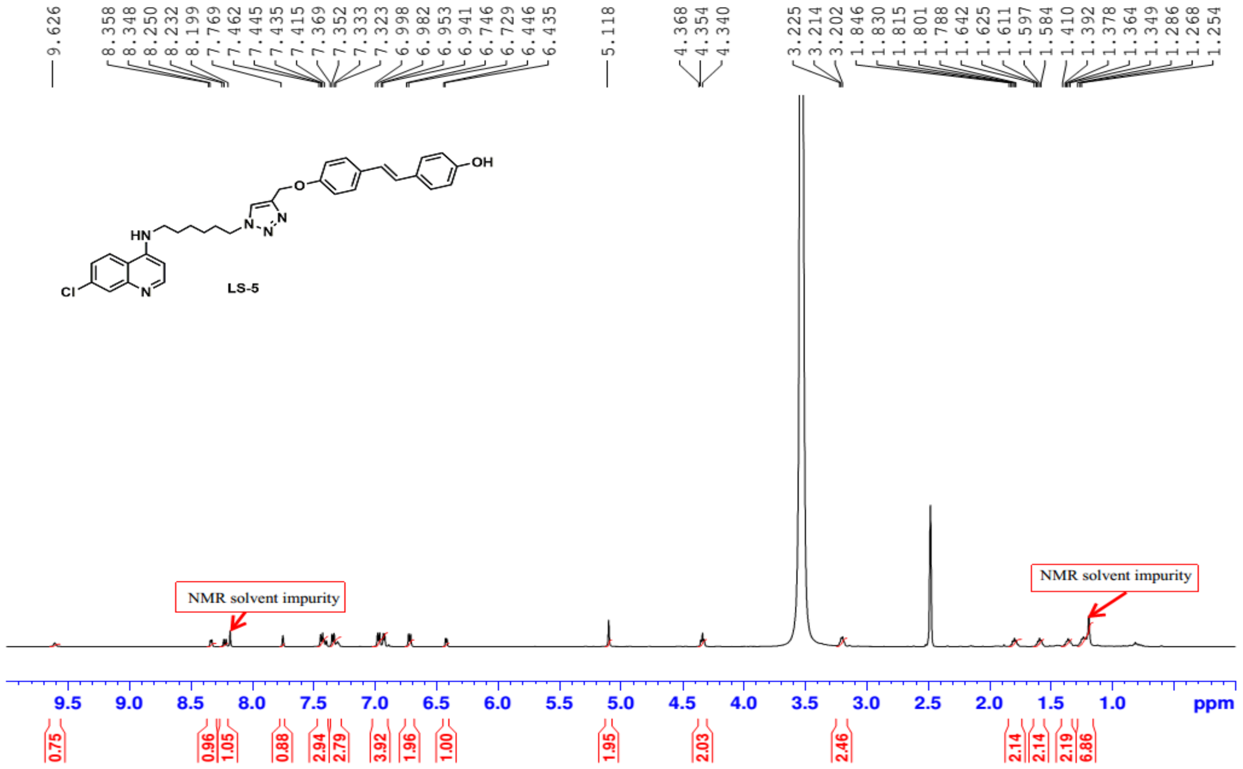

Fig. S33: $^{1}$H NMR spectrum of compound LS-5

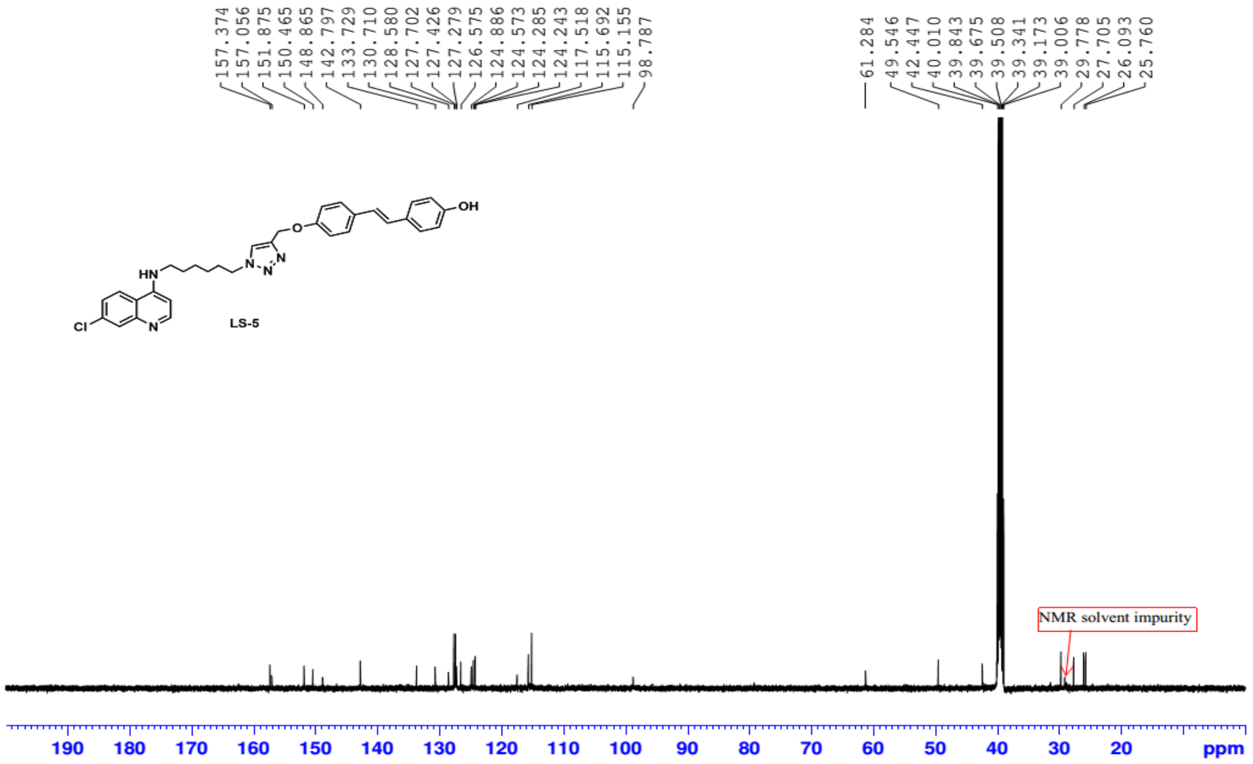

Fig. S34: $^{13}$C NMR spectrum of compound LS5

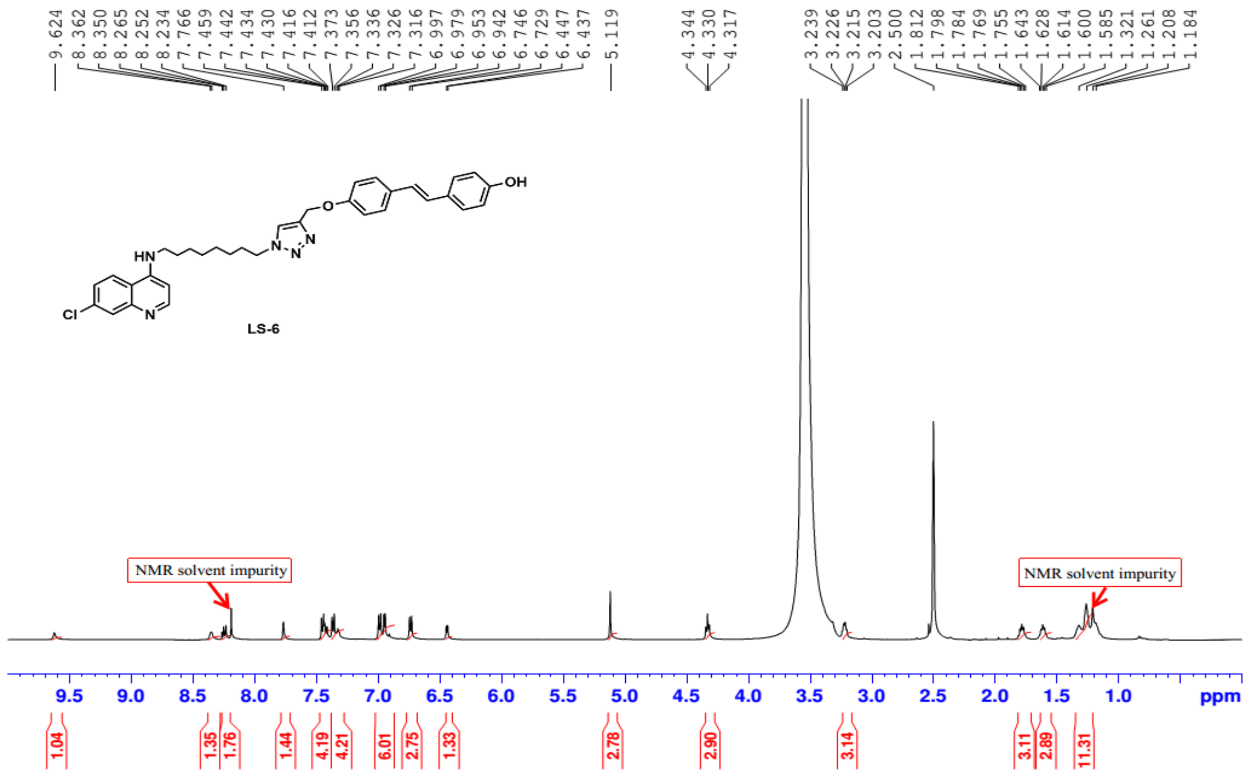


**Fig. S35: $^{1}$H NMR spectrum of compound LS6**

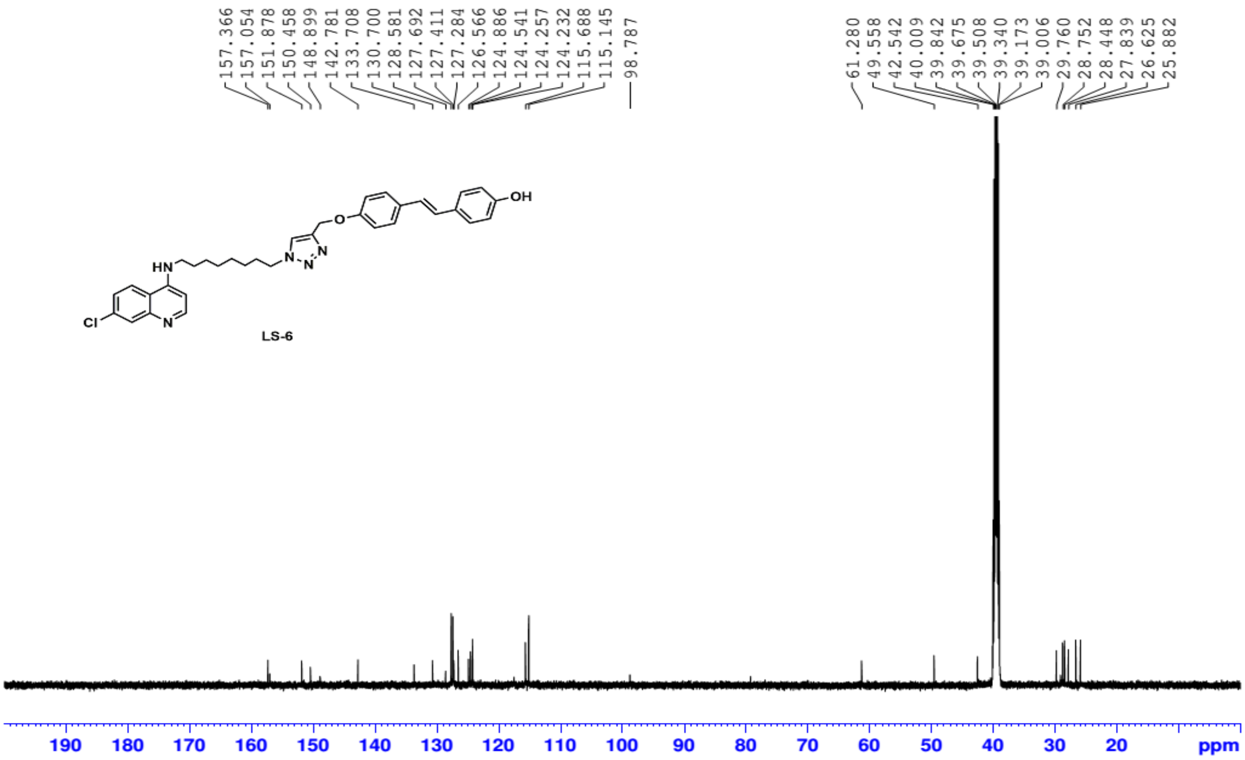


**Fig. S36: $^{13}$C NMR spectrum of compound LS6**

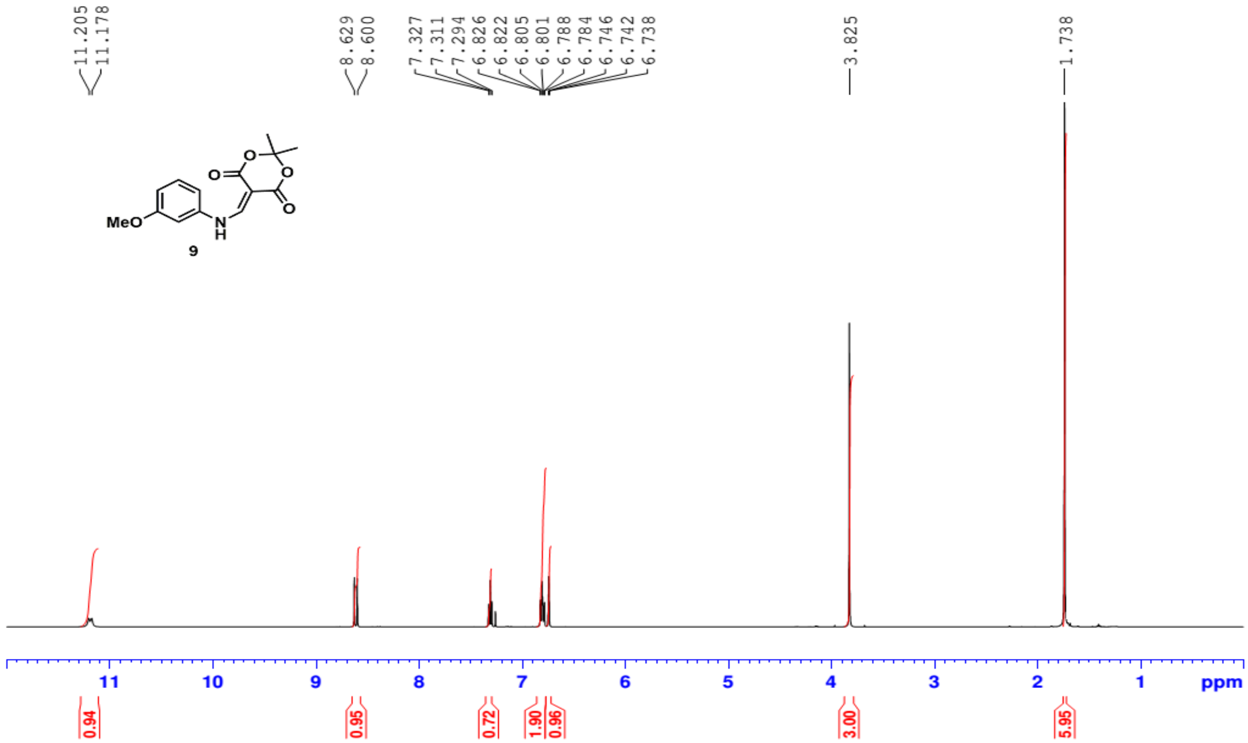


Fig. S37: $^{1}$H NMR spectrum of compound 9

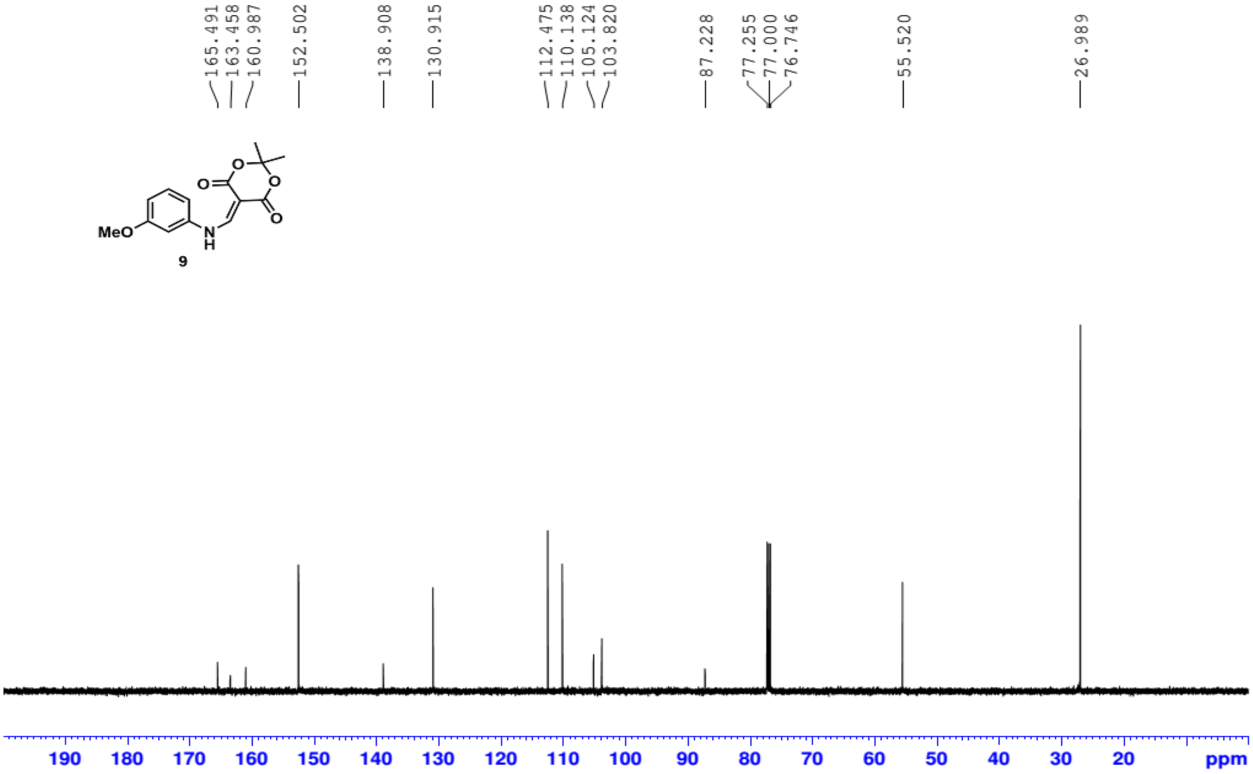


Fig. S38: $^{13}$C NMR spectrum of compound 9

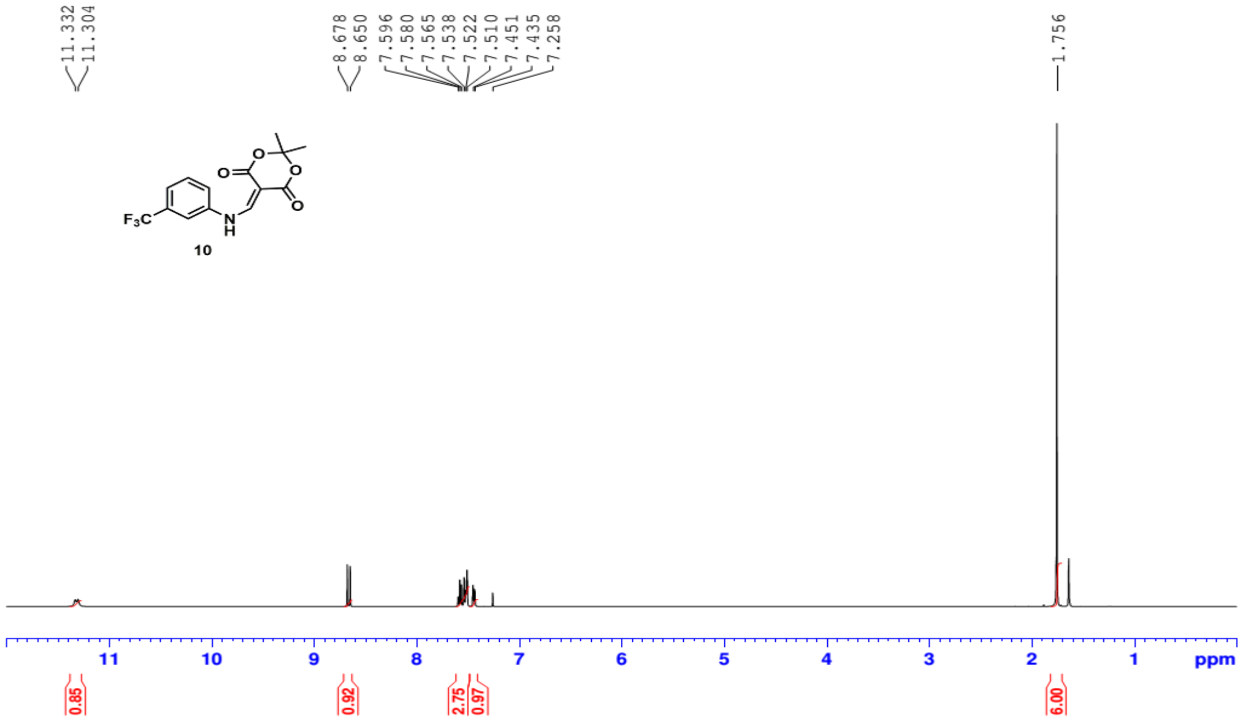


**Fig. S39: $^{1}$H NMR spectrum of compound 10**

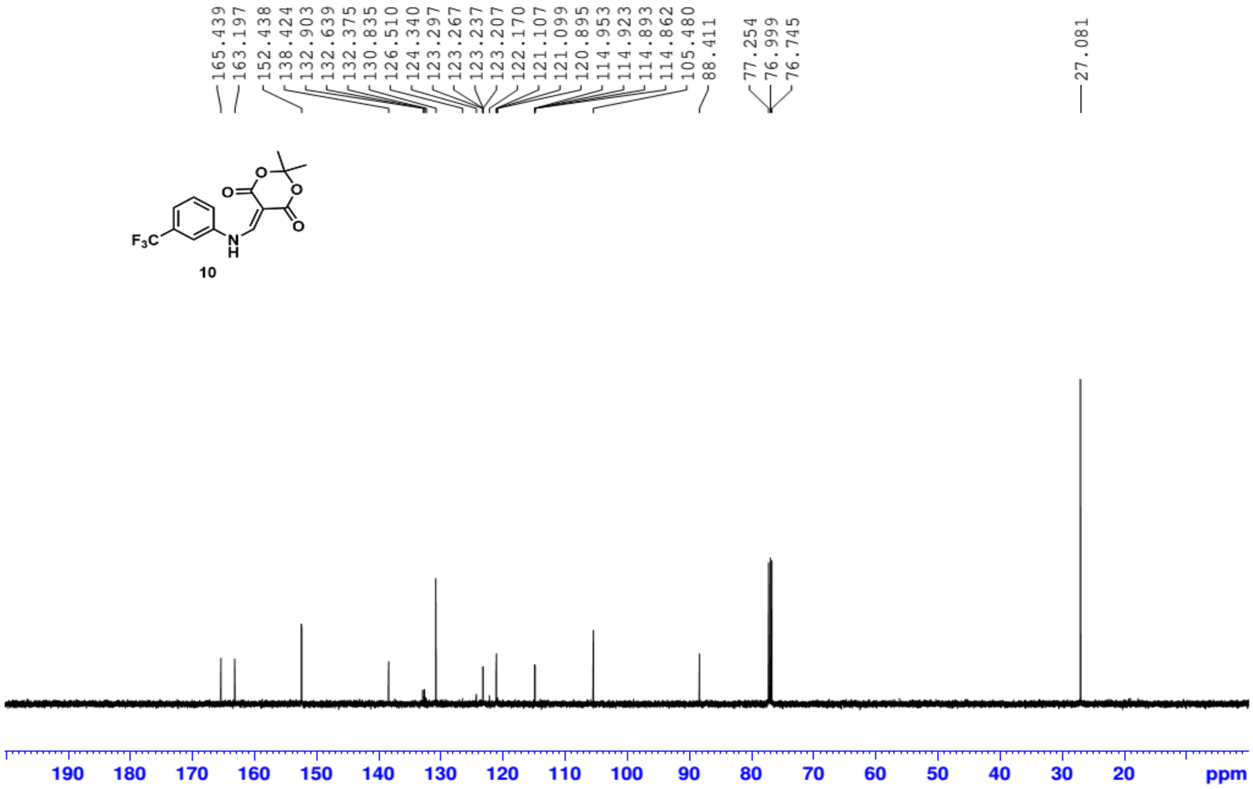


**Fig. S40: $^{13}$C NMR spectrum of compound 10**

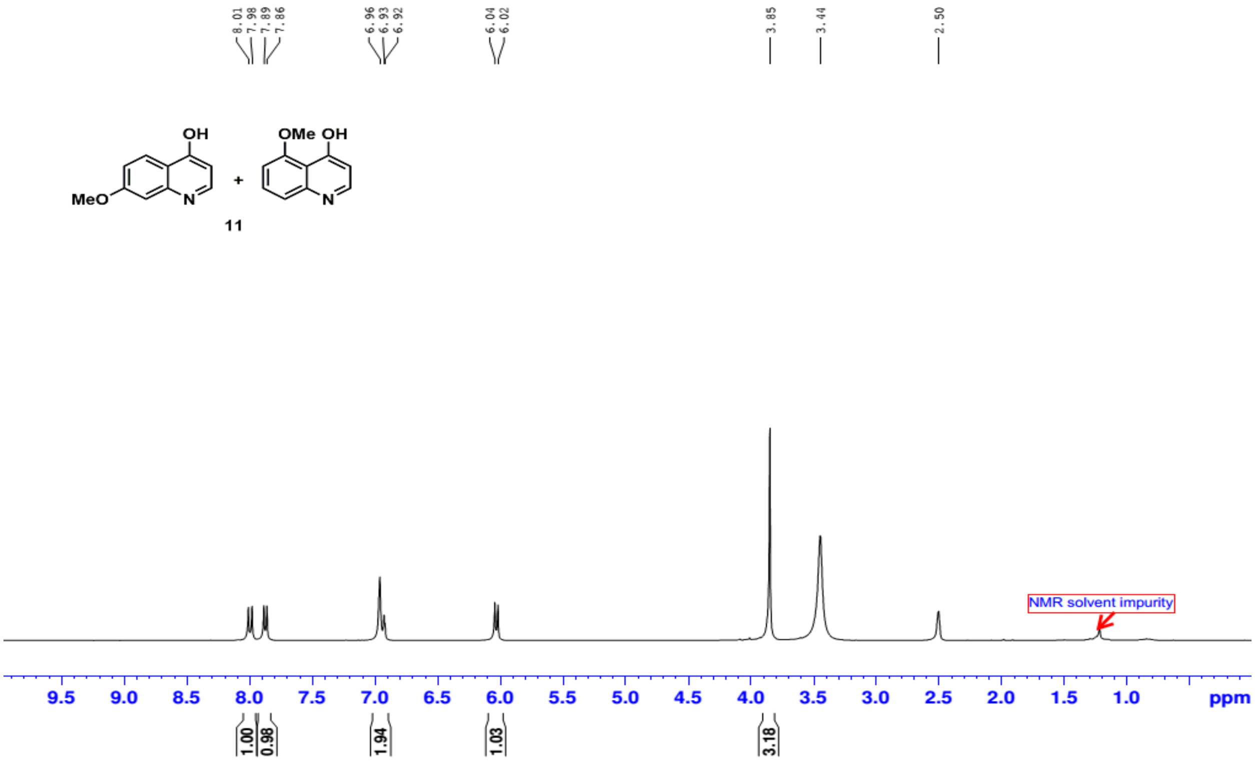

Fig. S41: $^{1}$H NMR spectrum of compound 11

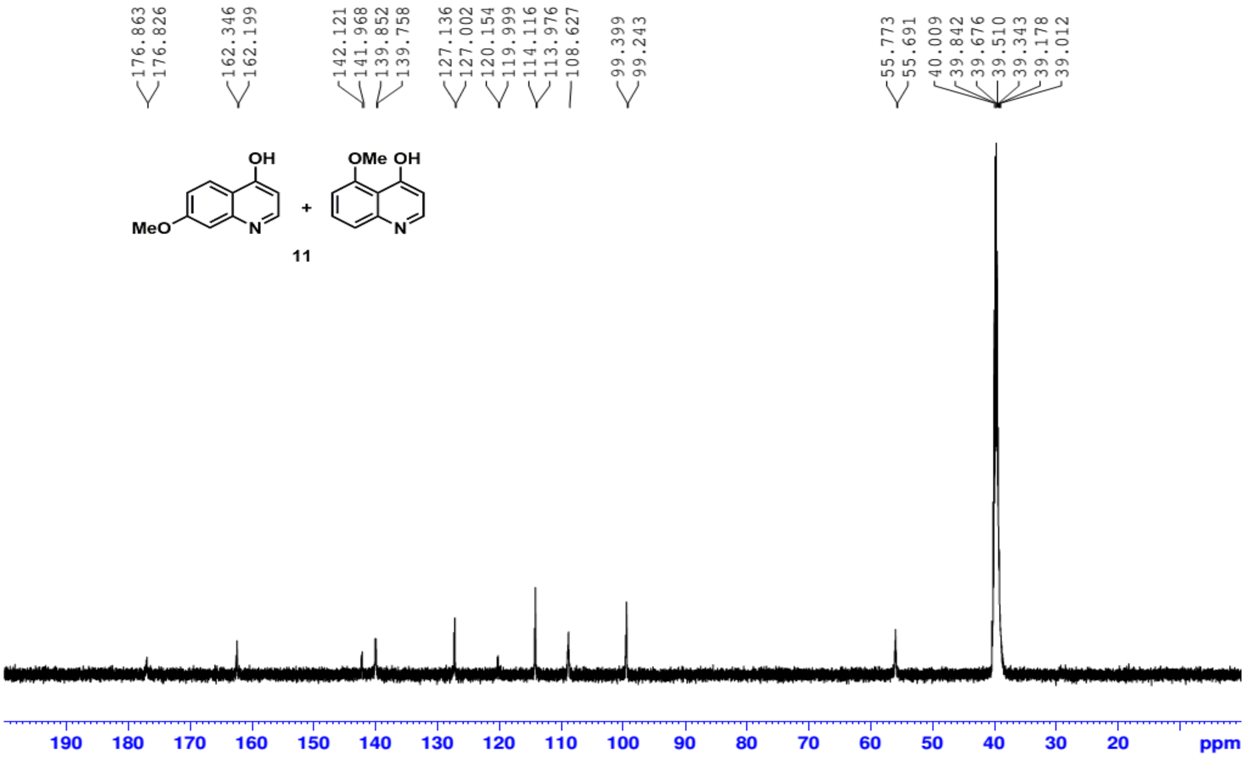

Fig. S42: $^{13}$C NMR spectrum of compound 11

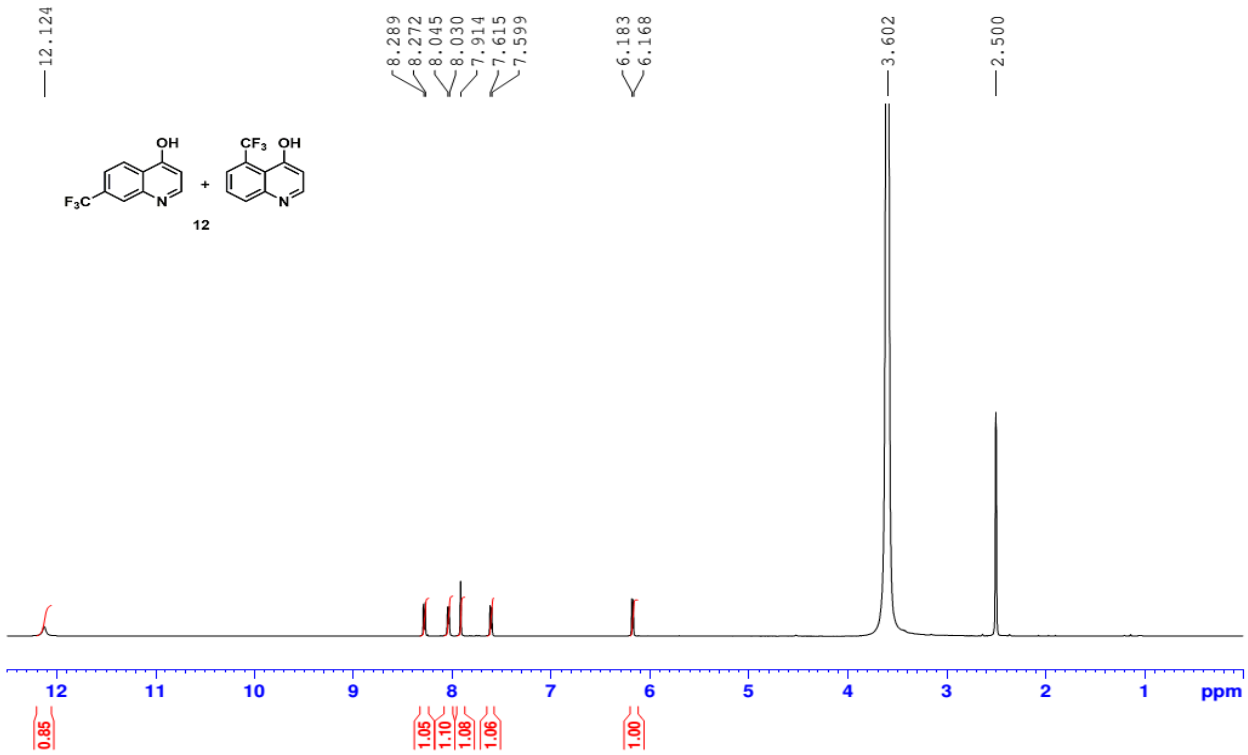


**Fig. S43: $^{1}$H NMR spectrum of compound 12**

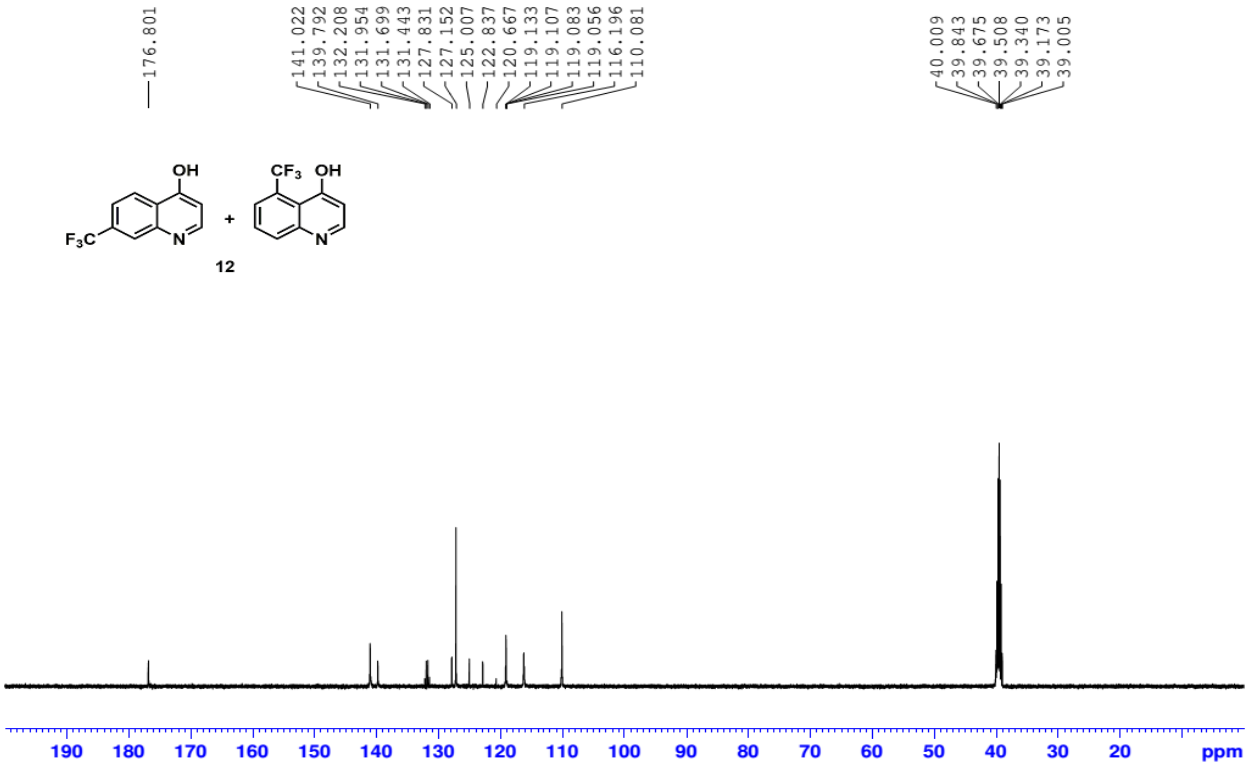


**Fig. S44: $^{13}$C NMR spectrum of compound 12**

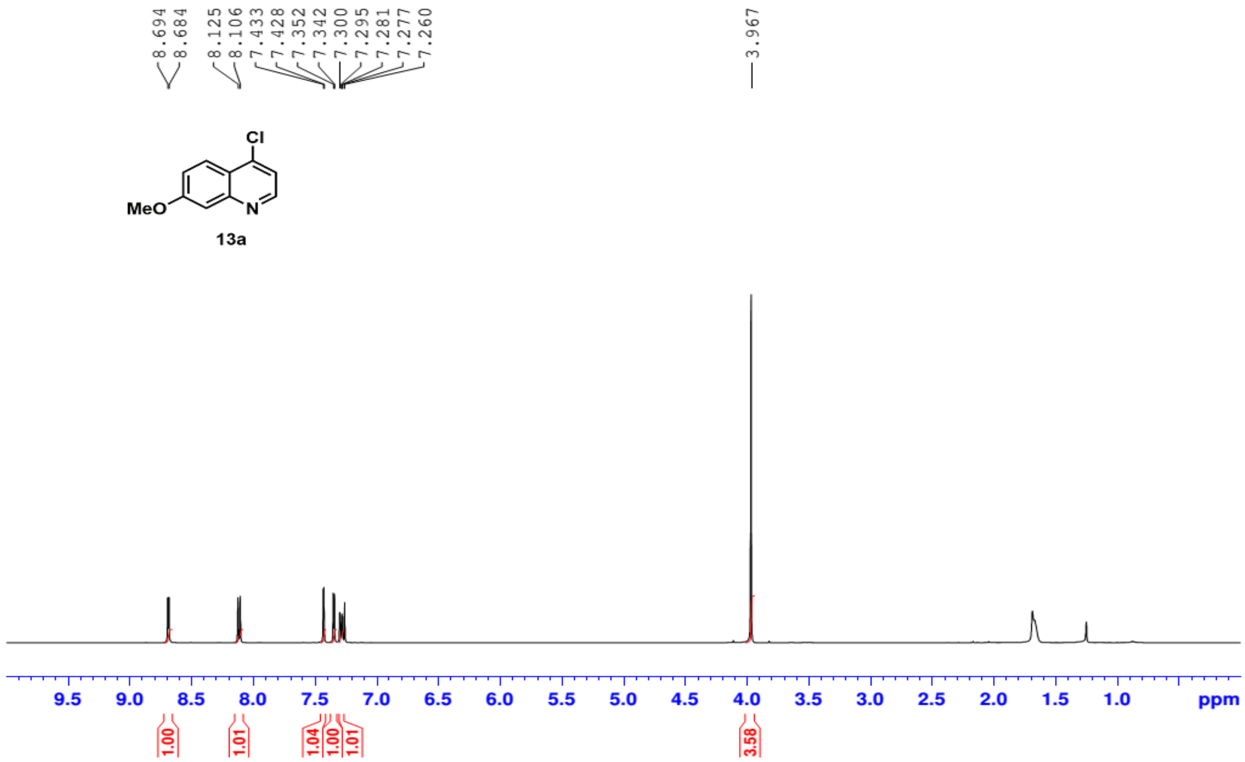


**Fig. 45: $^{1}$H NMR spectrum of compound 13a**

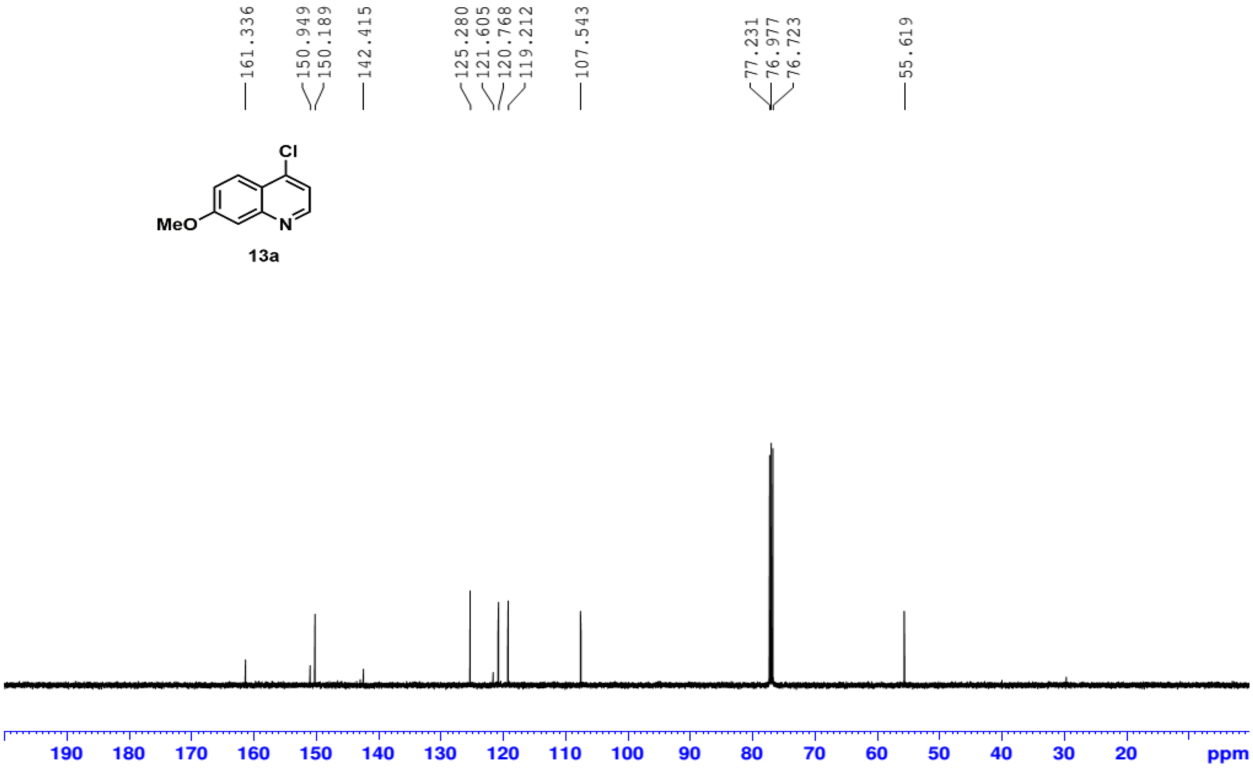


**Fig. 46: $^{13}$C NMR spectrum of compound 13a**

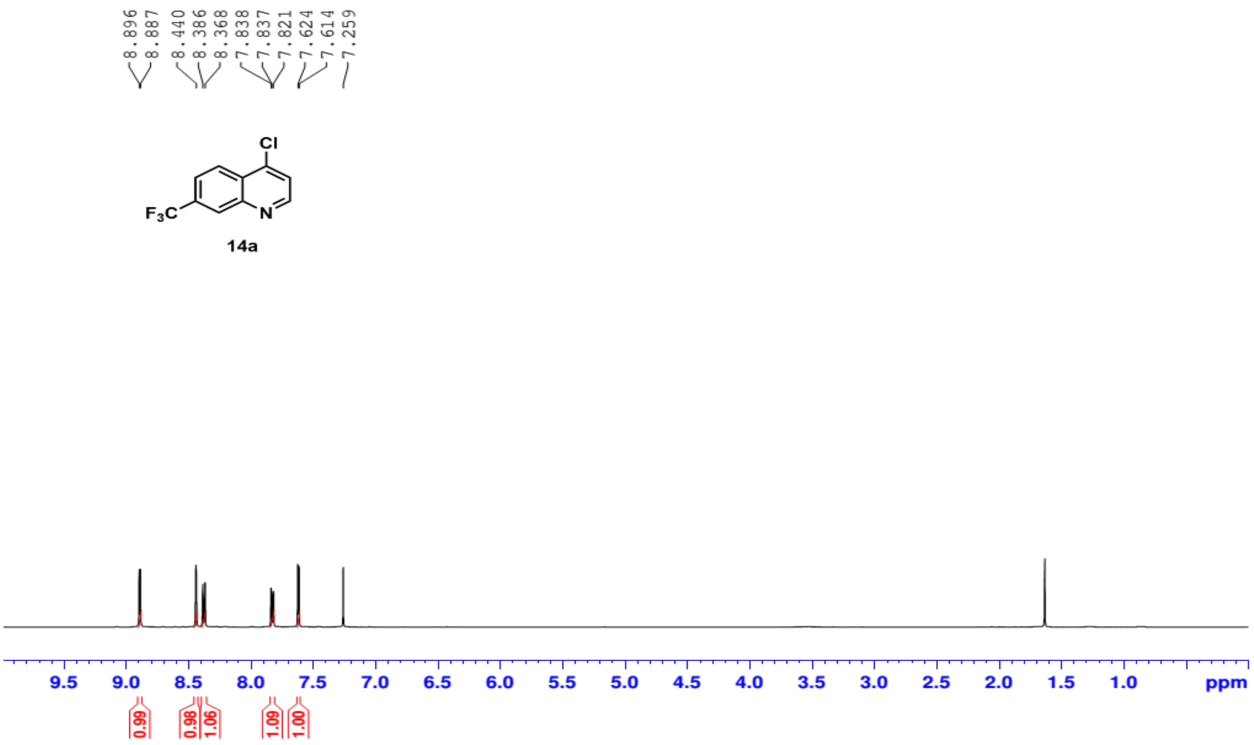


Fig. 47: $^{1}$H NMR spectrum of compound 14a

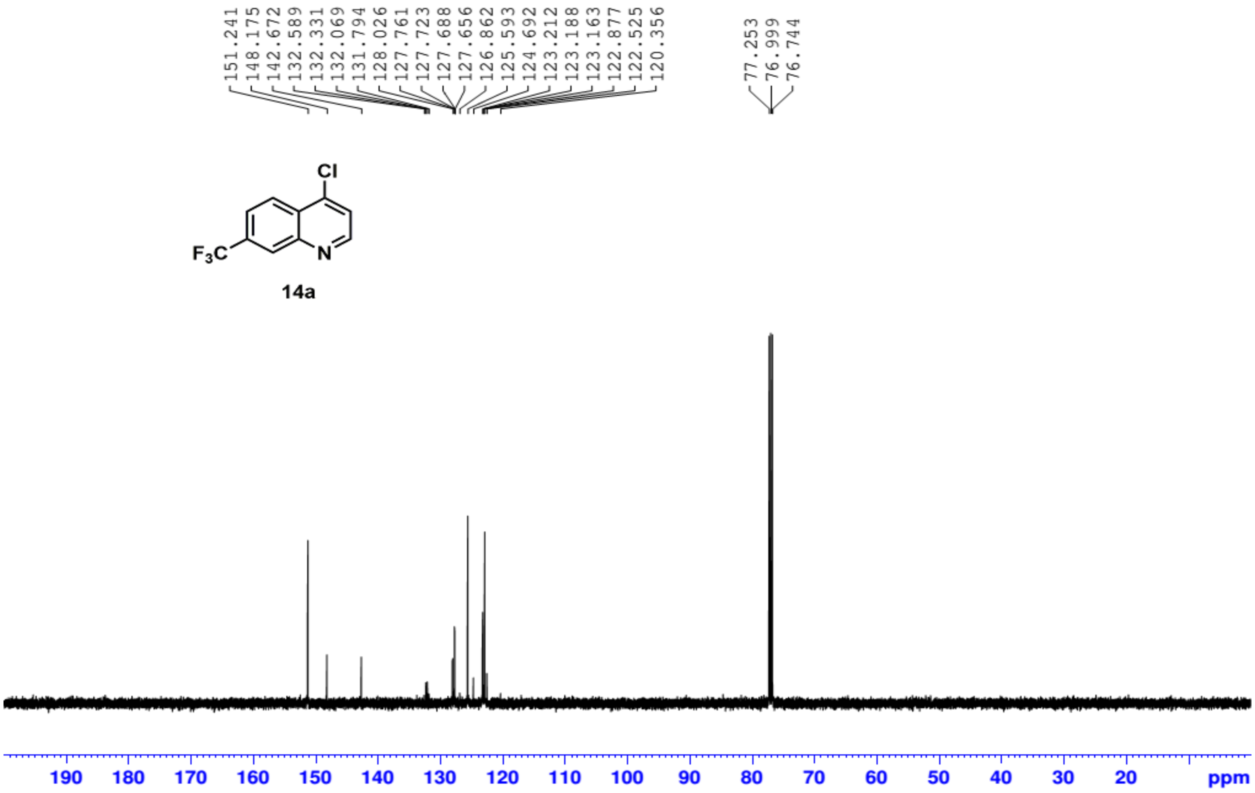


Fig. 48: $^{13}$C NMR spectrum of compound 14a

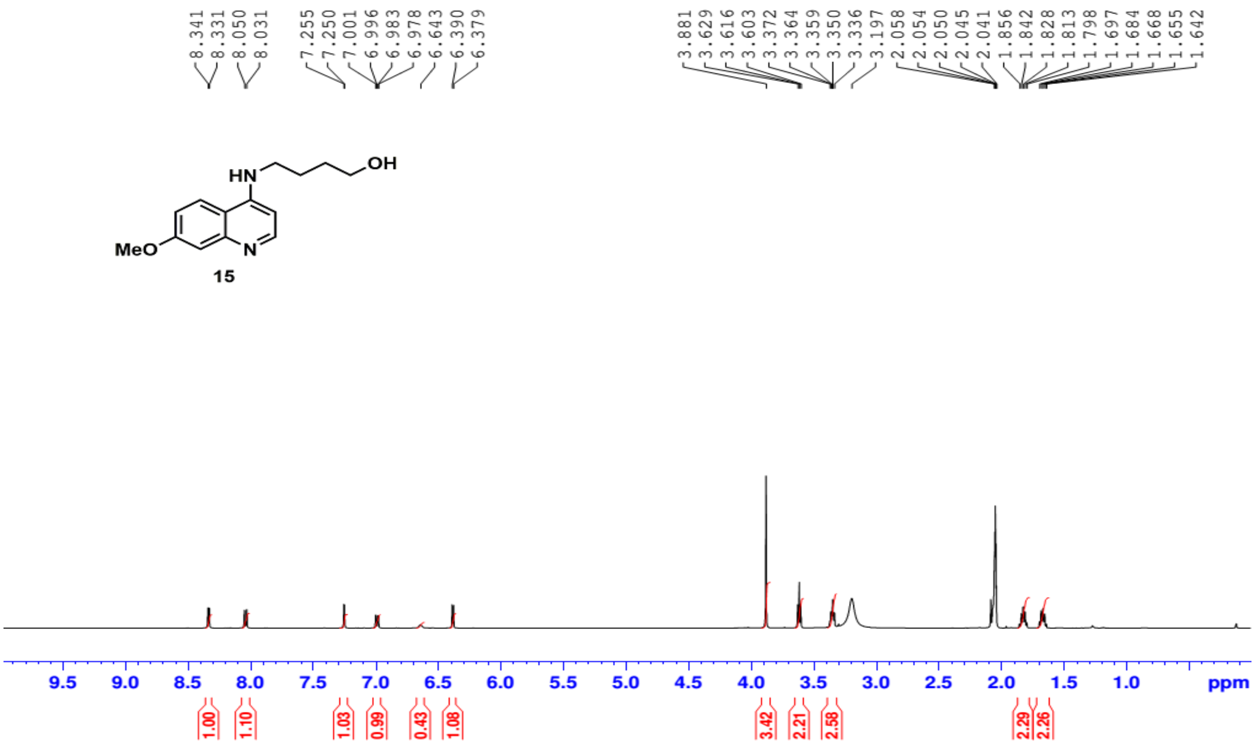

**Fig. S49: $^{1}$H NMR spectrum of compound 15**

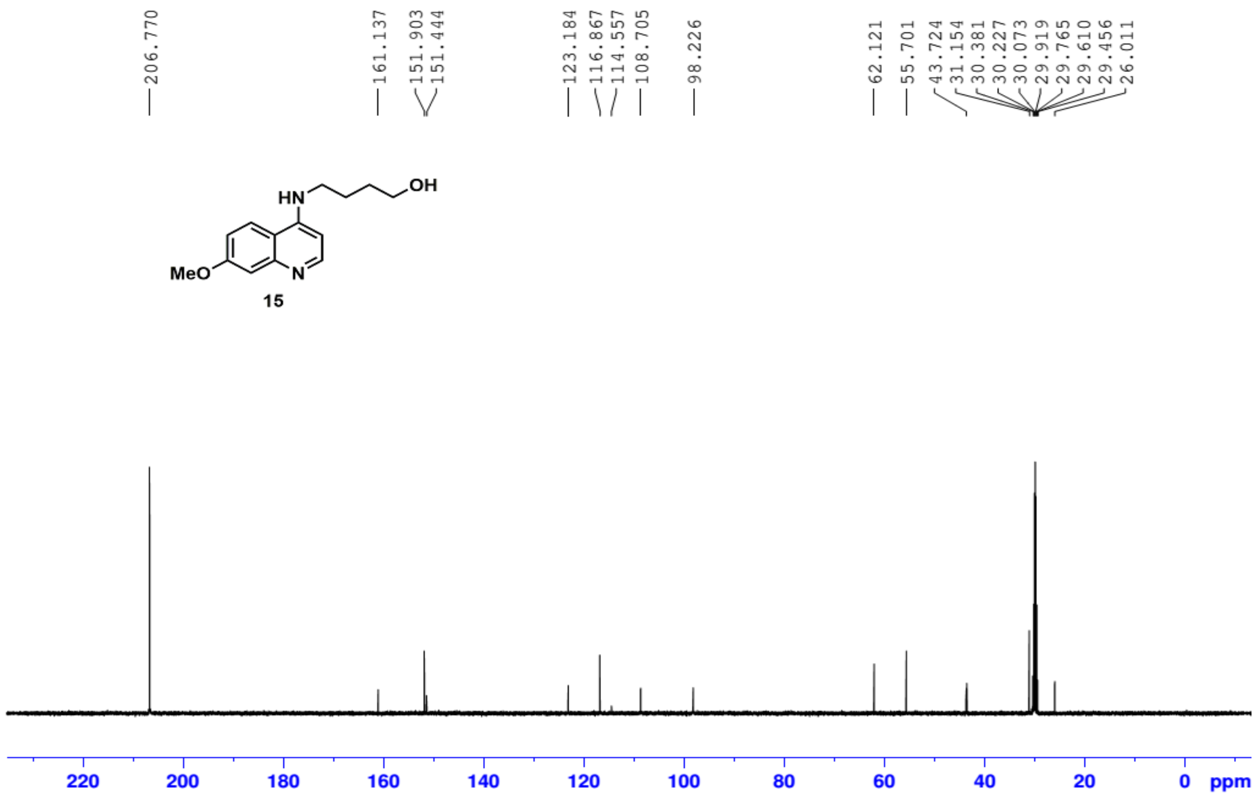

**Fig. S50: $^{13}$C NMR spectrum of compound 15**

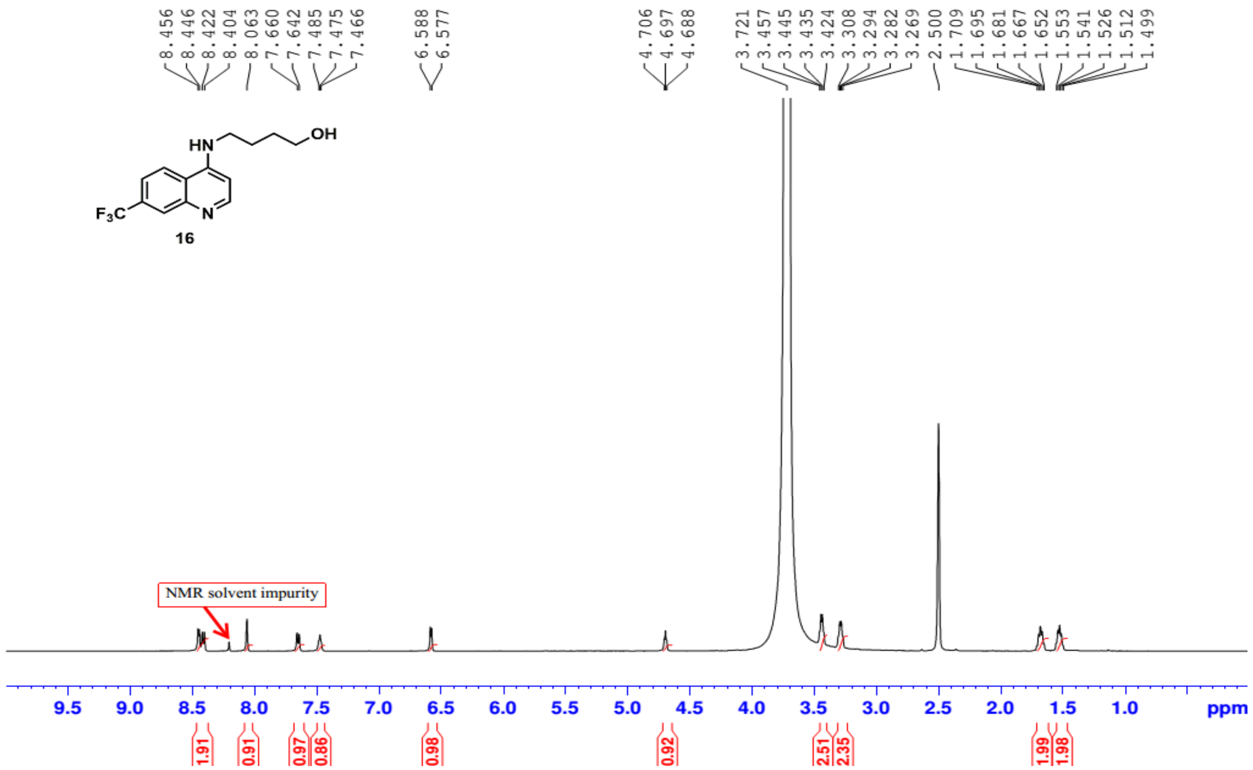


**Fig. S51: $^{1}$H NMR spectrum of compound 16**

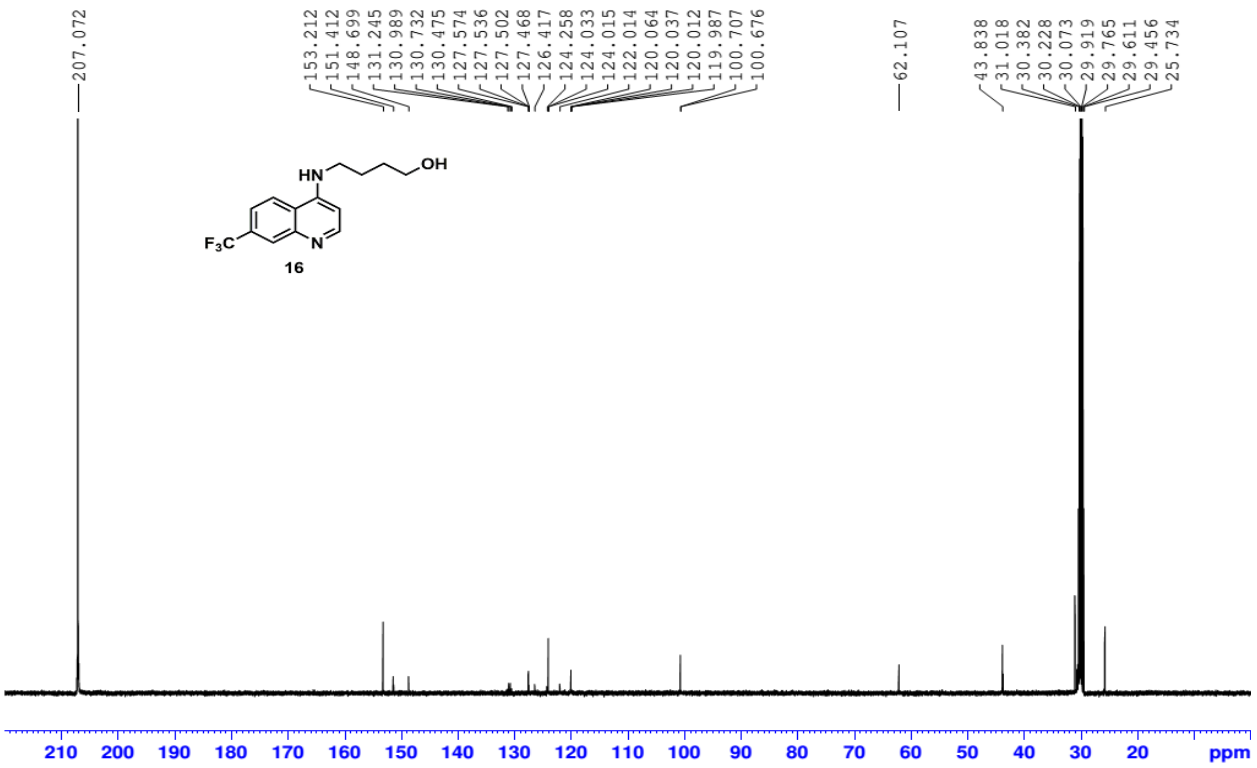


**Fig. S52: $^{13}$C NMR spectrum of compound 16**

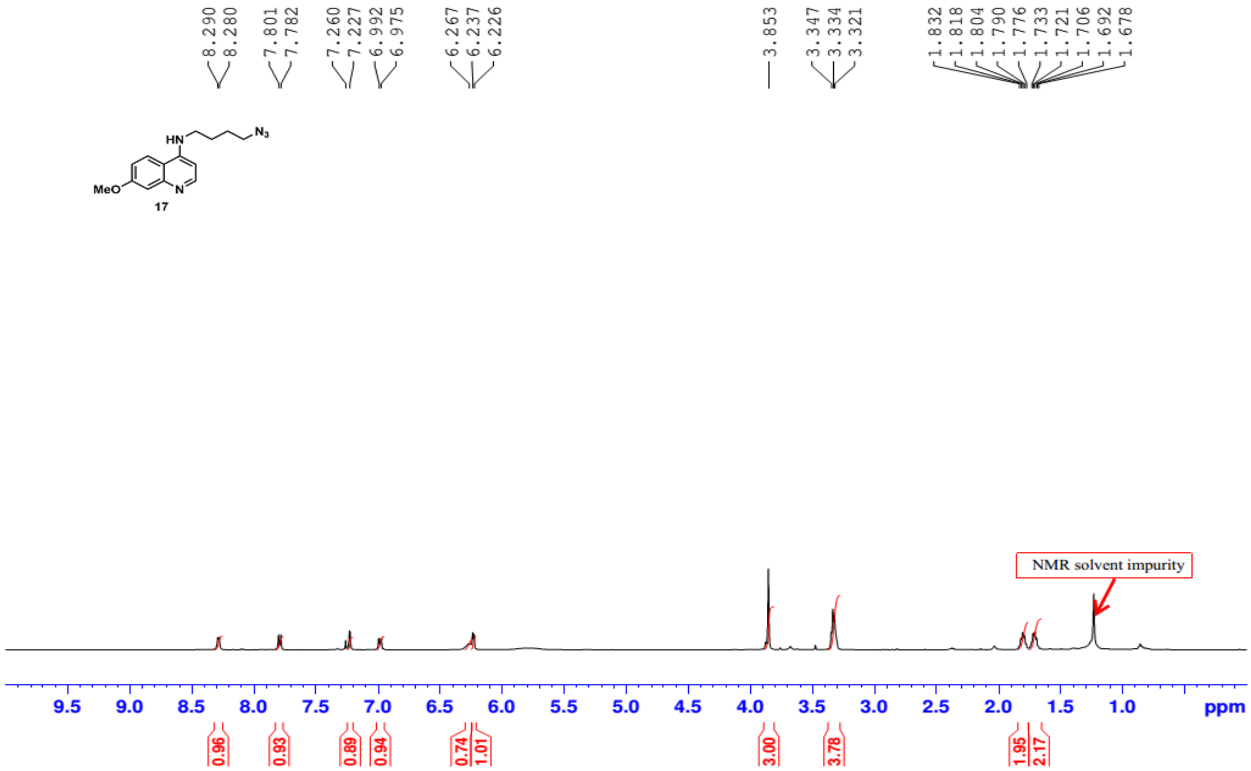


**Fig. 53: $^{1}$H NMR spectrum of compound 17**

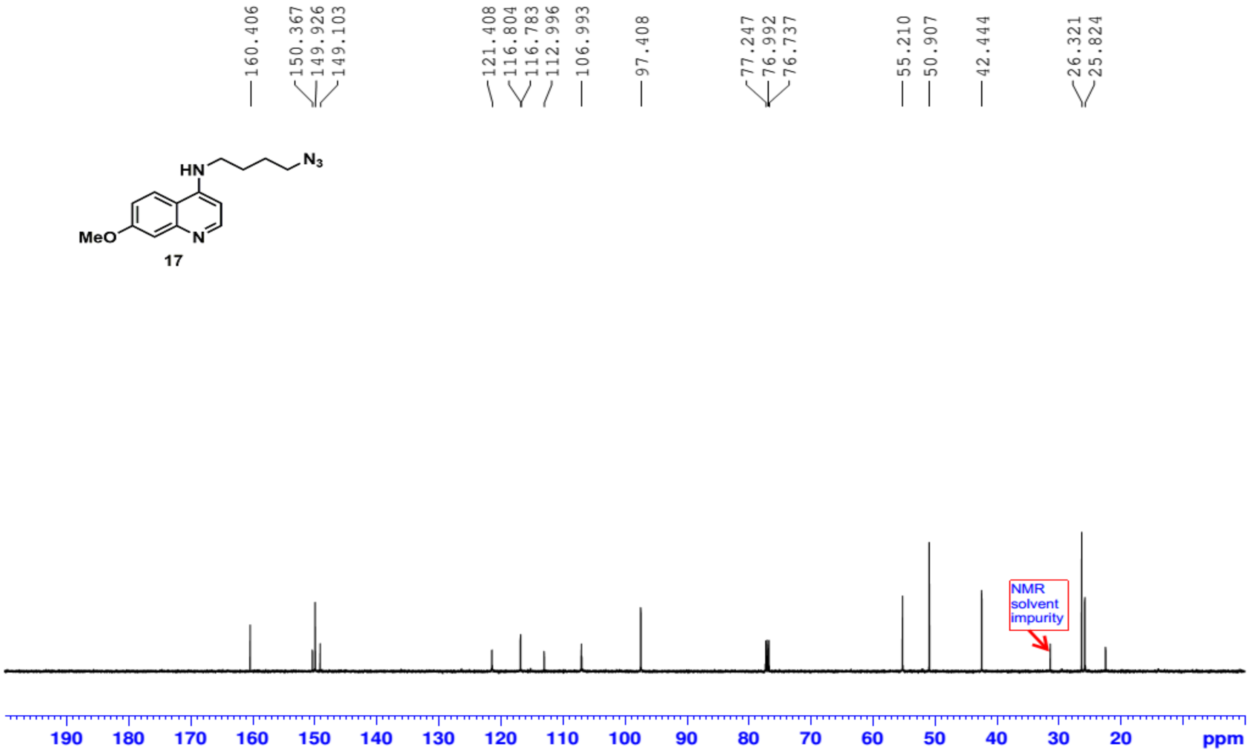


**Fig. 54: $^{13}$C NMR spectrum of compound 17**

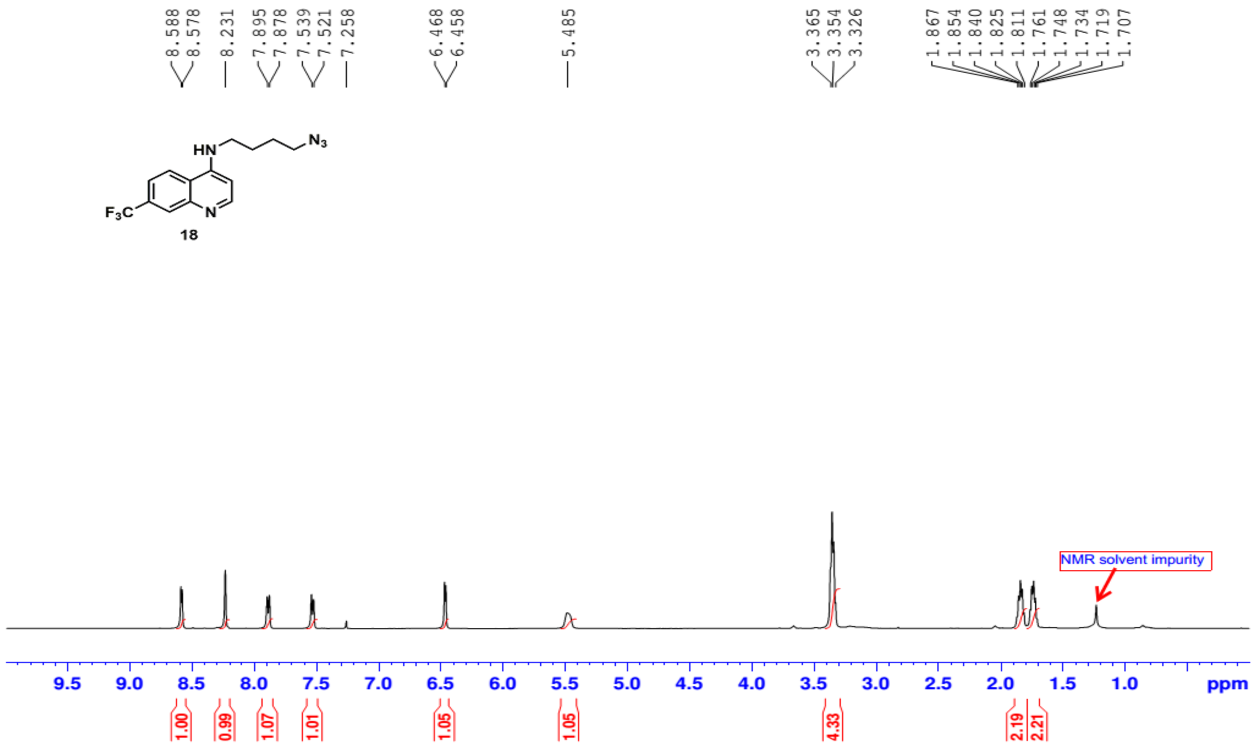


**Fig. S55: $^{1}$H NMR spectrum of compound 18**

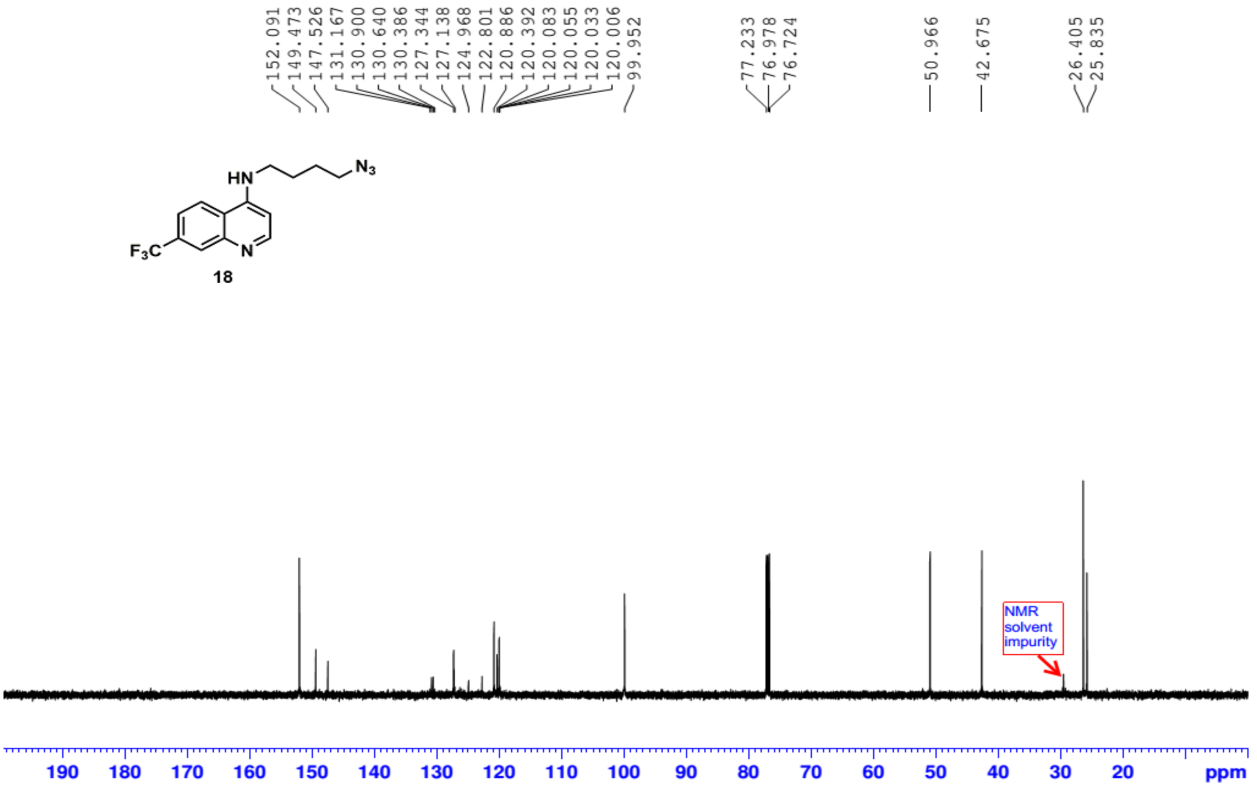


**Fig. S56: $^{13}$C NMR spectrum of compound 18**

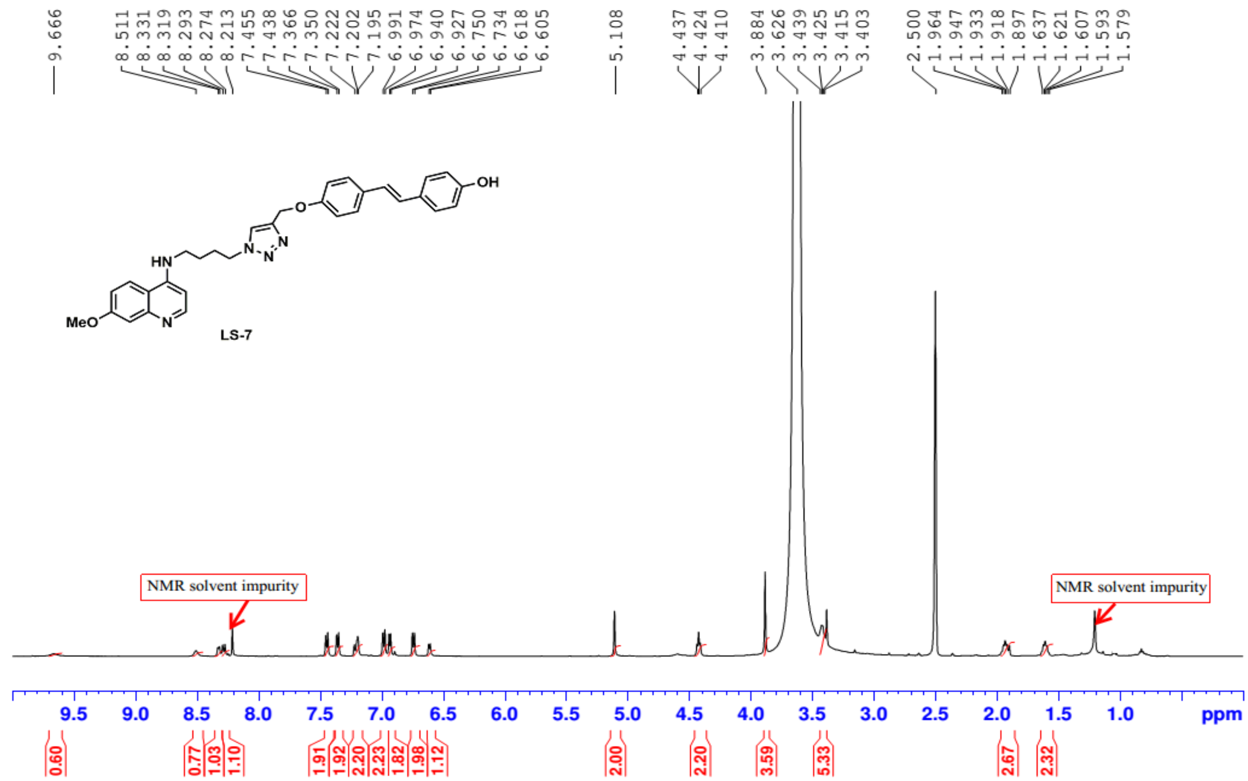


**Fig. S57: $^{1}$H NMR spectrum of compound LS7**

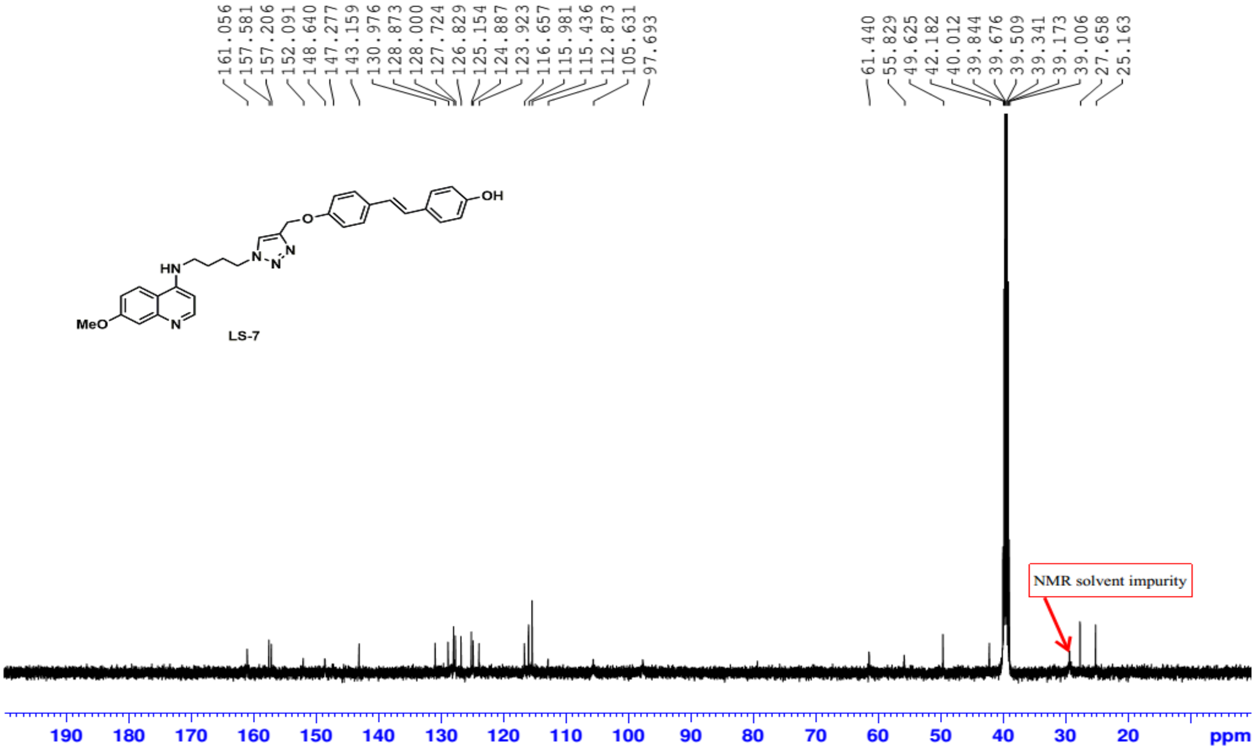


**Fig. S58: $^{13}$C NMR spectrum of compound LS7**

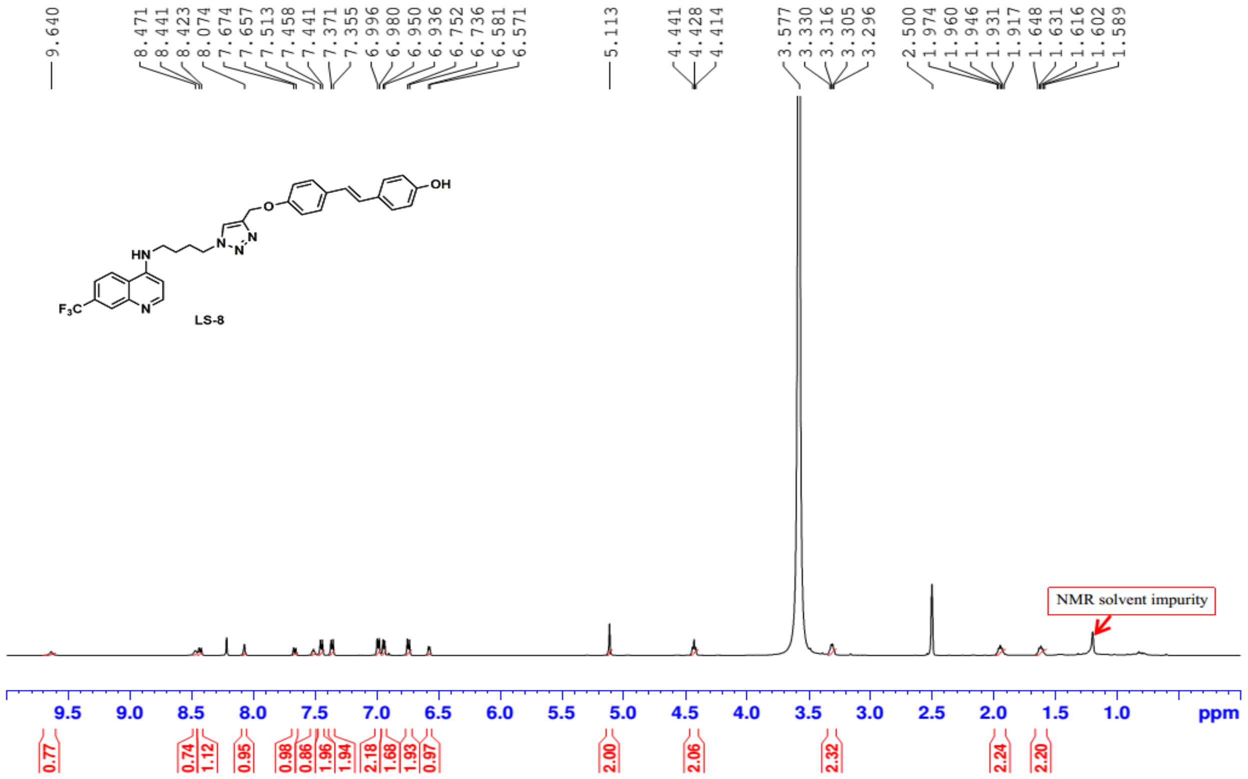

Fig. S59: $^{1}$H NMR spectrum of compound LS8

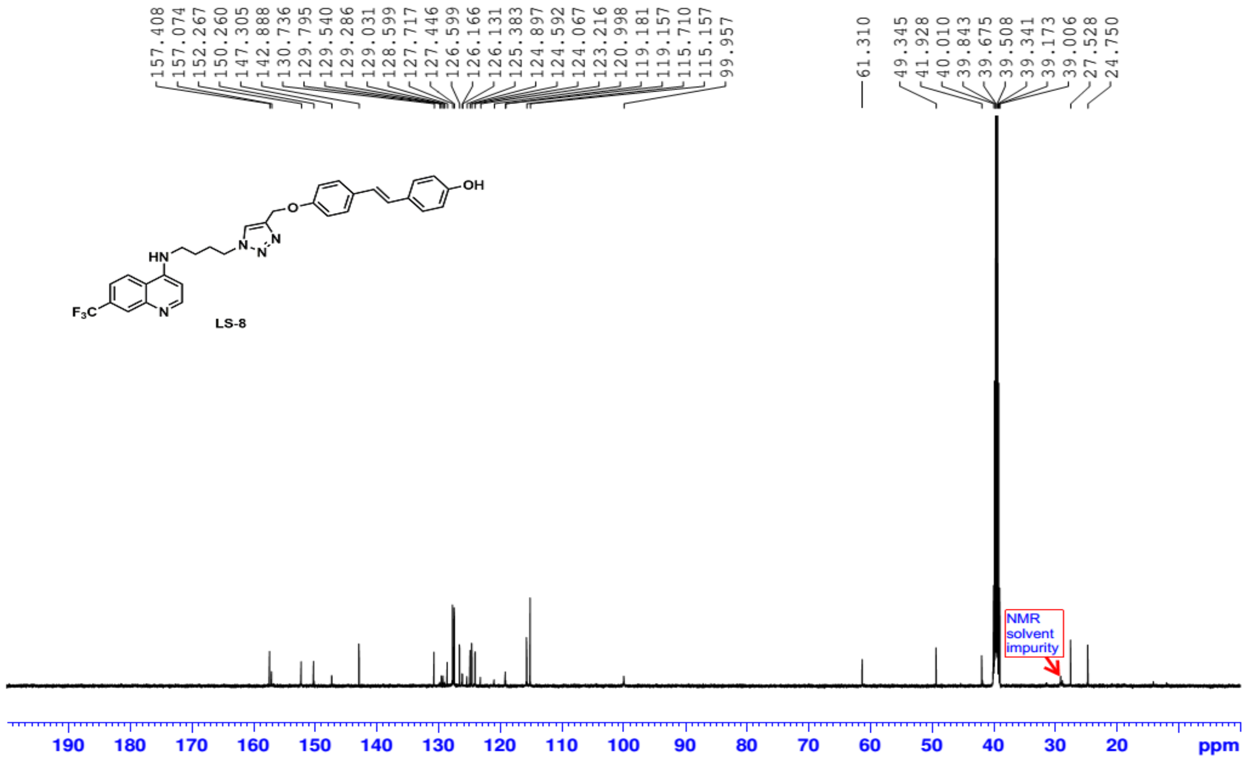

Fig. S60: $^{13}$C NMR spectrum of compound LS8

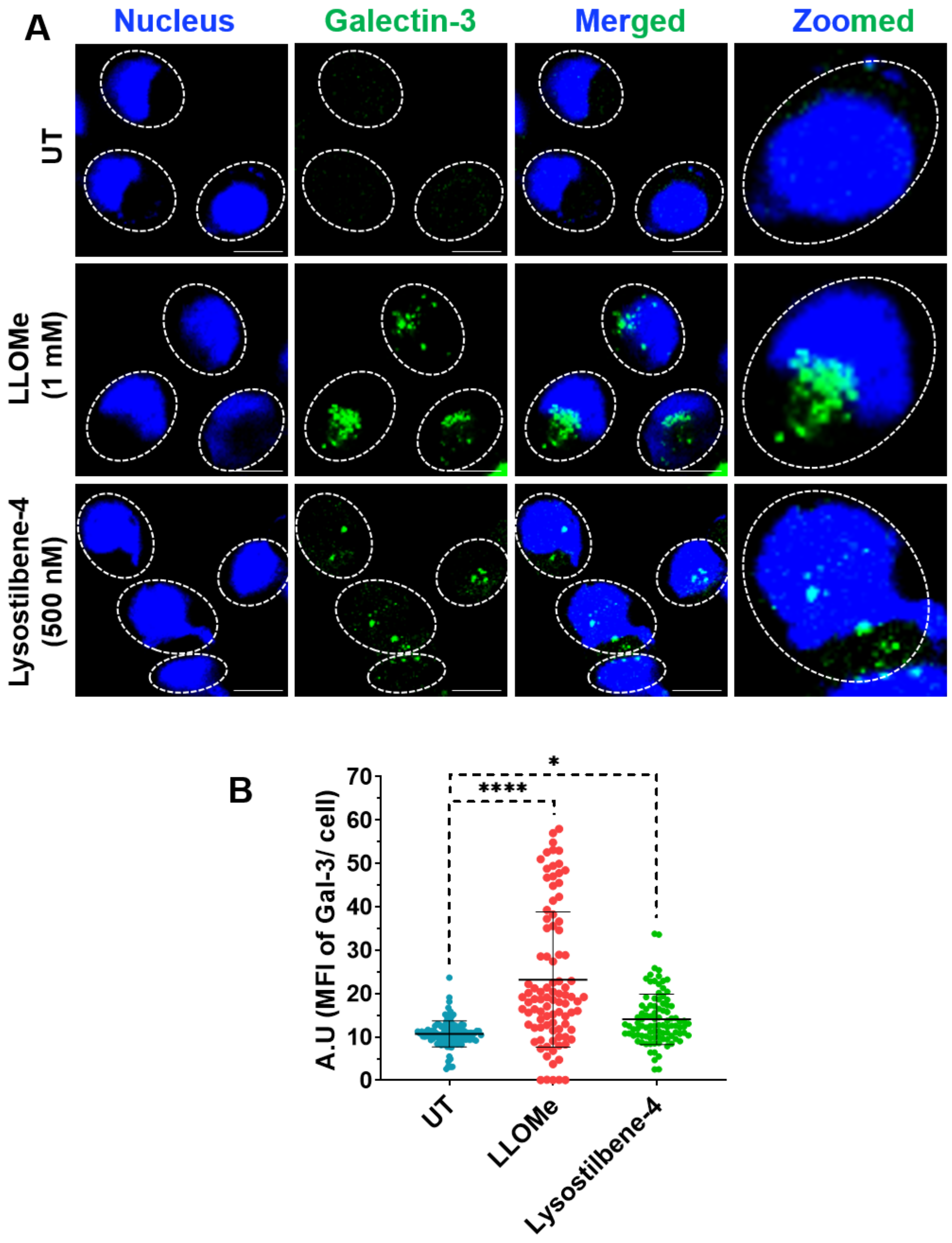


**Figure S61. Lysostilbene-4 induces moderate lysosomal damage.** (A) MIA PaCa-2 cells were treated with LLOMe or Lysostilbene-4 for 2 h. Representative confocal microscopy images show Galectin-3 (green) and nuclei stained with Hoechst (blue) under the indicated conditions. Scale bars: 10 µm. (B) Quantification of mean fluorescence intensity (MFI) of Galectin-3 per cell. Error bars represent mean ± SD, n= >120 cells per sample. (*$p < 0.05$; ****$p < 0.0001$).

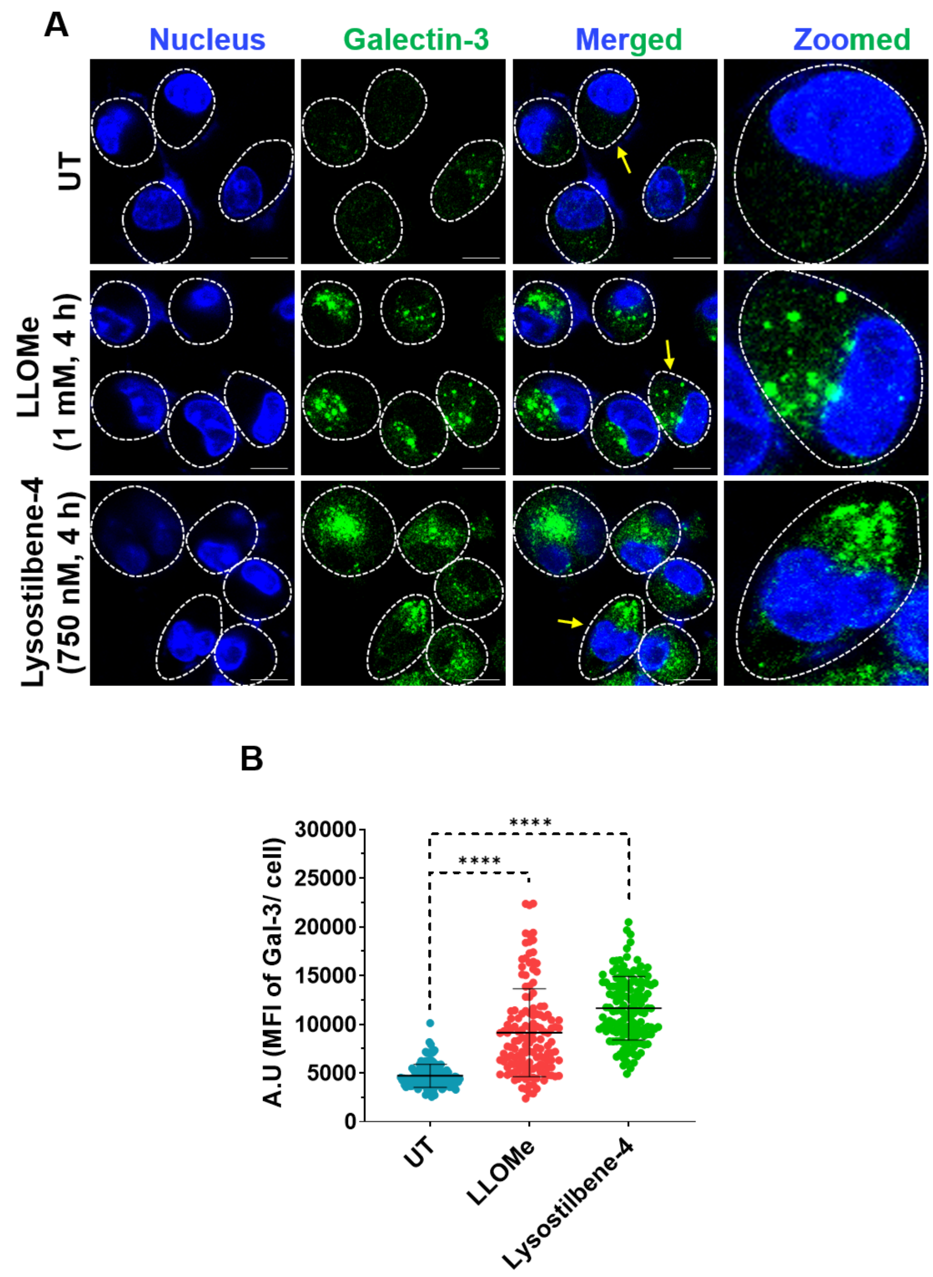


**Figure S62. Lysostilbene-4 (750 nM) induces robust lysosomal membrane permeabilization (LMP).** (A) Cells were treated with LLOMe or Lysostilbene-4 for 4 h. Representative confocal microscopy images show Galectin-3 (green) and Hoechst-stained nuclei (blue) under the indicated conditions. Scale bars: 10 μm. (B) The graph shows the quantification of Galectin-3 MFI per cell. Error bars represent mean ± SD, n= >130 cells per sample (****$p < 0.0001$).

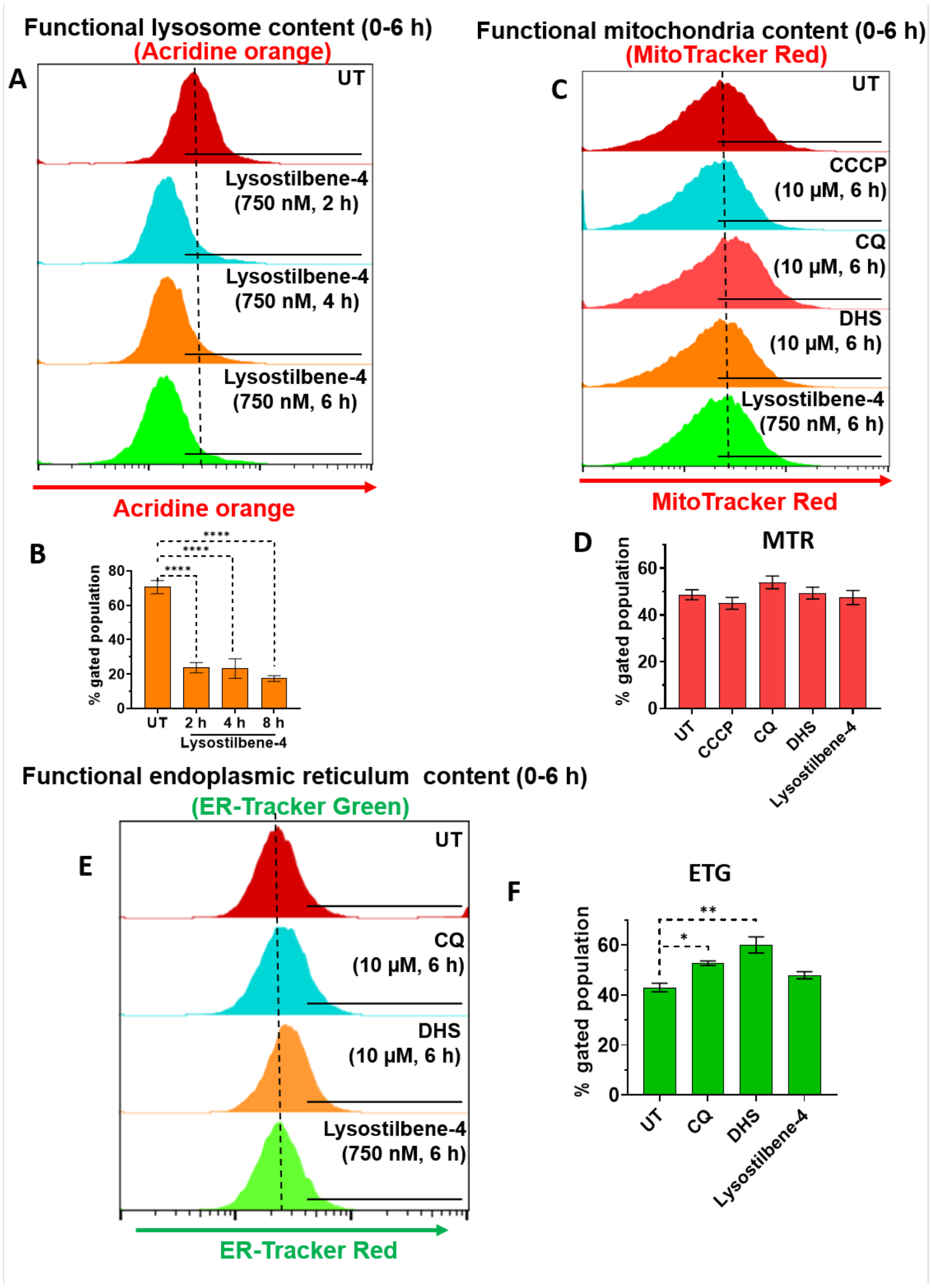


**Figure S63. Lysostilbene-4 induces selective damage to lysosomes.** (A, B) Lysosome damage: Cells were treated with Lysostilbene-4 for indicated time points and LMP was analyzed with reduced staining of lysosomes with AO (red). (C, D) Mitochondria damage: Cells were treated with Lysostilbene-4, CQ, DHS or CCCP for 6 h and loss of mitochondrial membrane potential was assessed by staining with Mitotracker Red, followed by flow cytometry. (E, F) Endoplasmic reticulum damage: Cells were treated with Lysostilbene-4, CQ or DHS for 6 h and ER content was assessed by staining with ERtracker Green, followed by flow cytometry. Error bars represent mean ± SD, N= 3 (*$p < 0.1$, **$p < 0.01$, ****$p < 0.0001$).

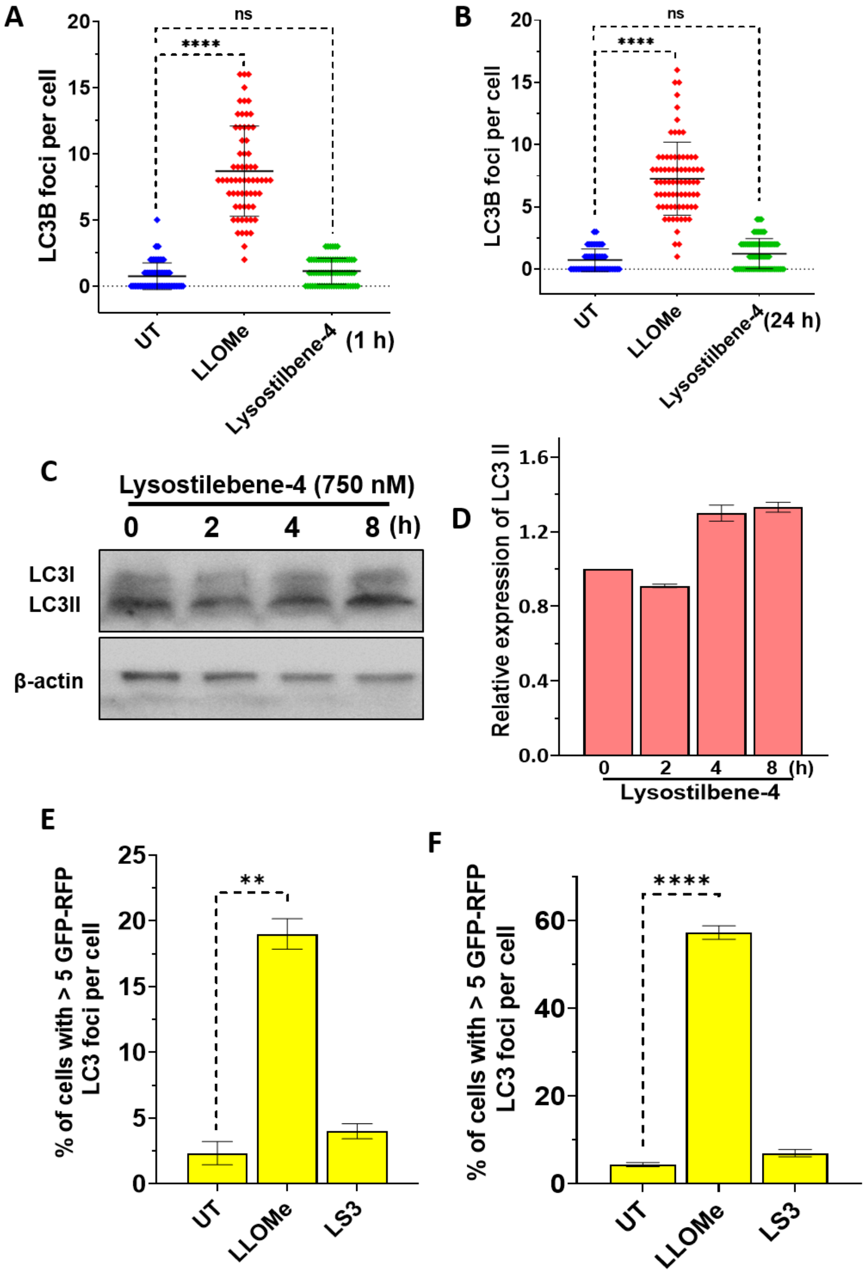


**Figure S64. Lysostilbene-4 inhibits autophagy initiation.** (A, B) The graphs show the quantification of LC3B puncta per cell from Figure 5E in UT, LLOMe, or Lysostilbene-4-treated cells for 1 h and 24 h. Error bars represent mean ± SD, n= >100 cells per sample. (C, D) Immunoblotting of LC3-I/II levels in cells treated with Lysostilbene-4 (750 nM) for the indicated time points, with β-actin as a loading control. Error bars represent mean ± SD, N= 3 biological replicates. (E, F) The graphs show the quantification from Figure 5G for the percentage of cells with >5 GFP-RFP-LC3 colocalized puncta per cell in untreated or LLOMe or LS3-treated conditions, at early (2 h) and late (24 h) time points. Error bars represent mean ± SEM, N= 3 biological replicates. (ns: not significant; **p < 0.01; ****p < 0.0001).

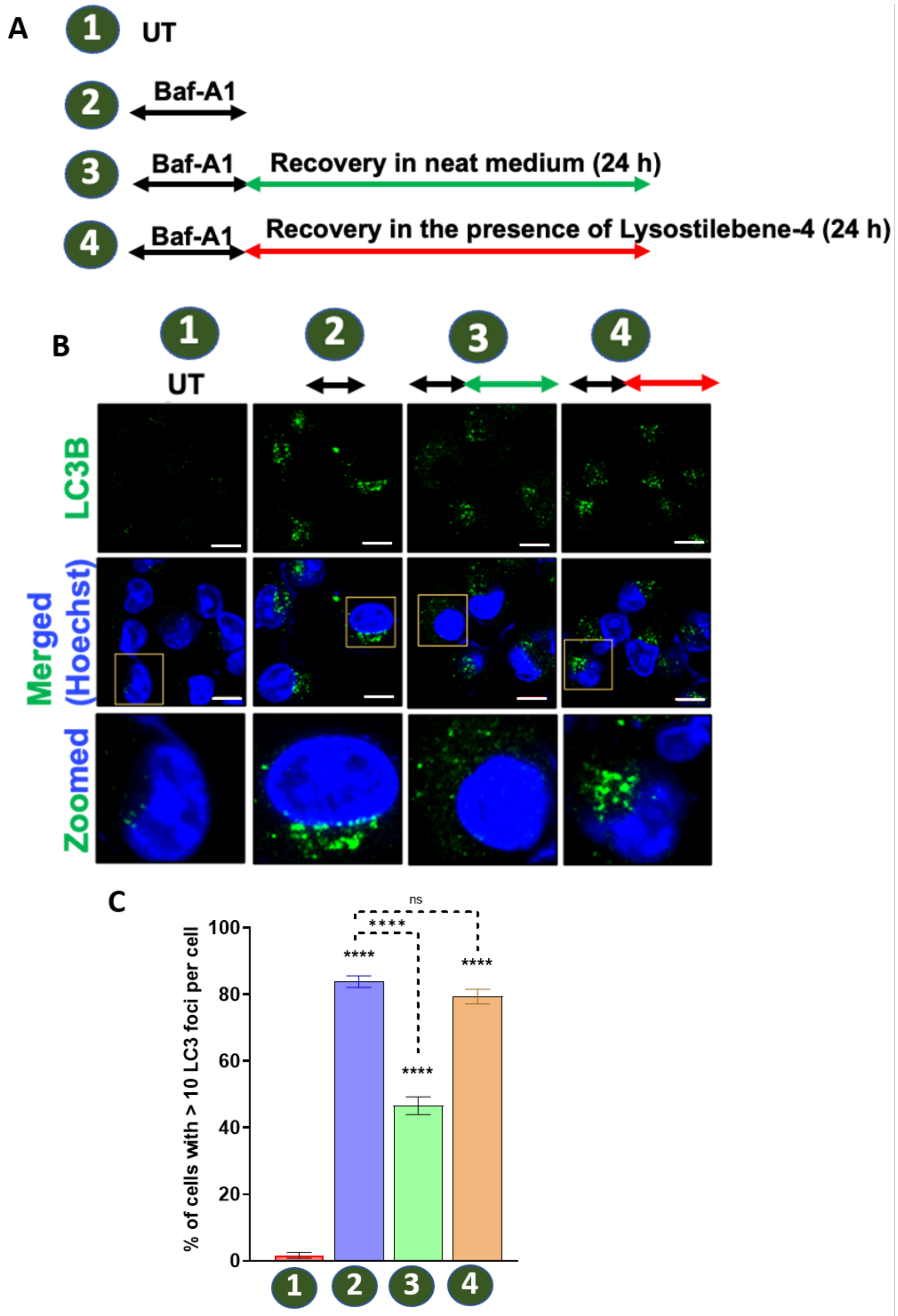


**Figure S65. Lysostilbene-4 inhibits autophagy-mediated clearance.** (A) Schematic of the experimental workflow to evaluate autophagosome clearance following bafilomycin A1-induced accumulation. (B, C) Representative confocal images showing LC3B puncta (green) and nuclei (Hoechst, blue) under the indicated conditions. Scale bars: 15 µm. Quantification of the percentage of cells with >10 LC3B puncta per cell across treatments was shown. Error bars represent mean ± SEM, N= 3 biological replicates. (ns: not significant; ****p < 0.0001).

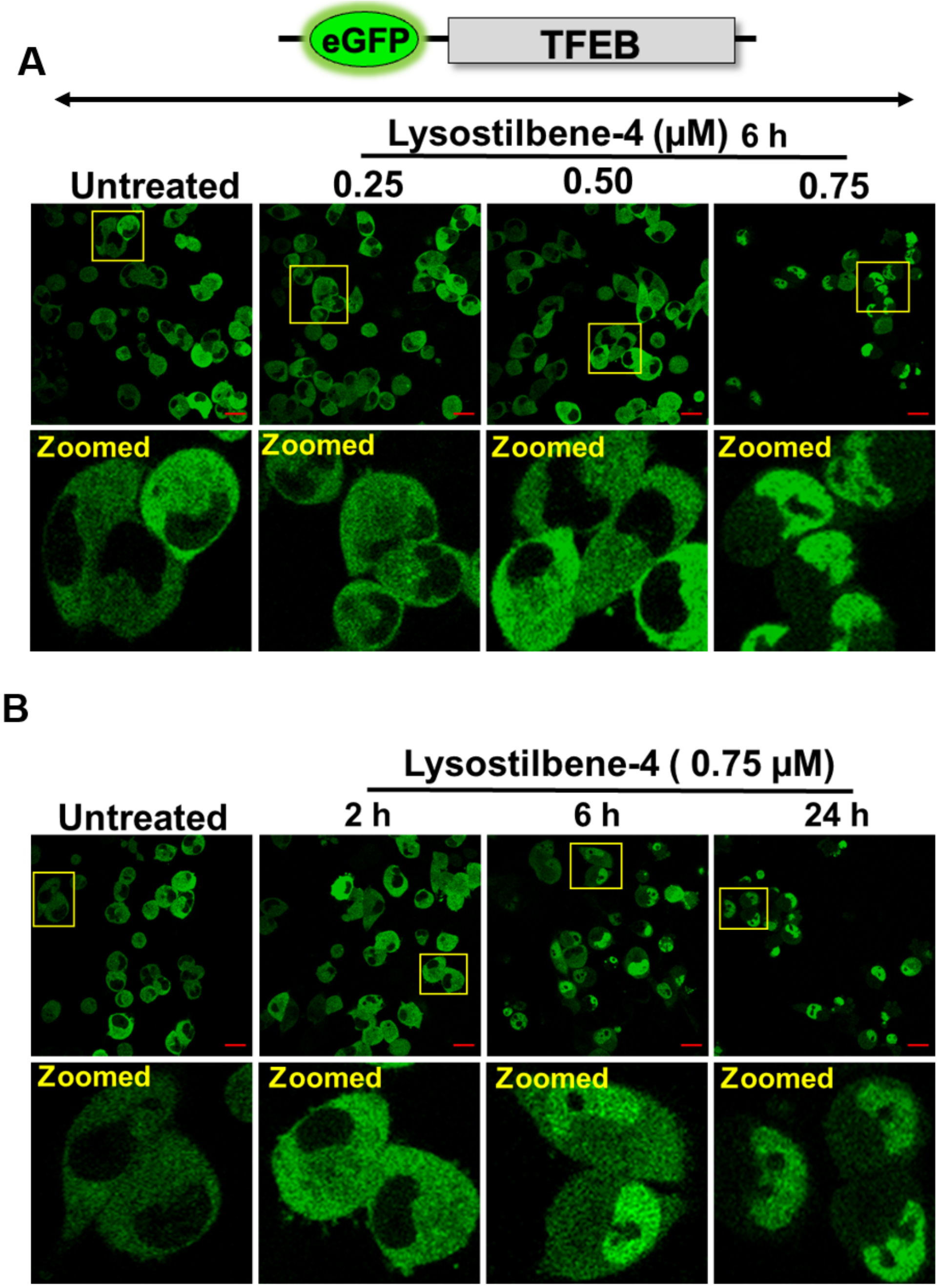


**Figure S66. Lysostilbene-4 induces TFEB nuclear translocation in a concentration- and time-dependent manner.** (A) Representative live-cell microscopy images of MIA PaCa-2 cells expressing eGFP-tagged TFEB upon treatment with increasing concentrations of Lysostilbene-4 (0.25–0.75 µM) for 6 h. (B) Representative images of cells treated with 0.75 µM Lysostilbene-4 for 2, 6, or 24 h. Yellow boxes indicate magnified regions shown in the lower panels. Scale bars:15 µm. (Note: Single cell zoomed images from these experiments are presented in Figure 6A, C).
